# A cholinergic eligibility trace facilitates amygdala plasticity in flavour avoidance learning

**DOI:** 10.64898/2026.07.02.734913

**Authors:** Weston Fleming, Jordan L. Pauli, Kentaro K. Ishii, Marta Trzeciak, Cassidy T. Burke, Sekun Park, Ruohe Liu, Adam G. Gordon, Larry S. Zweifel, Richard D. Palmiter, Garret D. Stuber

## Abstract

When an animal feels ill after consumption of a novel food, it rapidly learns to avoid this food in the future, a form of learning termed conditioned flavour avoidance (CFA). Postingestive malaise often occurs long after consumption, necessitating a neural mechanism that can facilitate plasticity between temporally distant events. Neuromodulators, acting through G-protein-coupled receptors (GPCRs) that can influence neuronal excitability on extended timescales, may facilitate this process. The projection of parabrachial (PB) *Calca* neurons to the central amygdala (CeA) promotes formation of CFA. Here, we demonstrate that these neurons overlap with a PB population that releases the neuromodulator acetylcholine (ACh) in the CeA during both consumption of a novel solution and subsequent visceral malaise, consistent with a role in CFA acquisition. Calcium imaging in brain slices reveals that ACh widely activates CeA neurons and enhances glutamatergic responsivity on a timescale consistent with CFA learning. CRISPR-Cas9-mediated genetic knockdown and optogenetics demonstrate that ACh from PB facilitates CFA acquisition. Loss of ACh signalling to CeA blocks key signatures of CFA-associated plasticity. Together, these data reveal that cholinergic input from PB to central amygdala is a critical neuromodulatory signal that links activity over long timespans to facilitate associative learning.

## Introduction

Animals evaluate novel foods based on their flavour and postingestive effects. Toxic foods can cause nausea (or visceral malaise), vomiting, and diarrhoea with a delay of minutes to hours after consumption. Despite these delays, animals are able to form robust food-malaise associations with just one or two pairings, a highly adaptive form of learning referred to as conditioned flavour avoidance (CFA) or conditioned taste/flavour aversion^1–10^.

How animals learn to correctly associate an outcome to the causal event that preceded it remains a central question in associative learning. Long delays should impede this process: intervening events and the decay of the causal memory interfere with the association^11^. CFA defies these conventions.

The neural mechanisms that facilitate long-delay associative learning are not well understood. One candidate mechanism for CFA acquisition involves neuromodulatory signalling during novel food consumption. Such a neuromodulator, acting through GPCRs, could prime neurons for preferential reactivation and plasticity processes later during visceral malaise^12^. The correlates of such a process– activation of the same neurons both during novel food consumption and during later malaise–have been observed in the central amygdala (CeA)^13^, but the mechanism that permits this selective reactivation is unknown. The CeA is well-positioned as a site of CFA plasticity: it receives a convergence of excitatory inputs from regions including the insular cortex^14–16^, basolateral amygdala^17^, and PB that transmit gustatory, visceral, and aversive information^18–20^. In turn, the CeA regulates appetitive consumption^21–25^, inhibiting consumption if danger is detected^17,26^.

Notably, the CeA receives a dense projection from PB *Calca* neurons^27^, which are active during malaise and critically important for CFA learning^13,28–34^. PB*^Calca^* neurons release calcitonin gene-related peptide (CGRP) and several other neuropeptides^27^, but CGRP itself is largely dispensable for CFA learning^30,35^, raising the possibility that another neuromodulator may have a primary role in CFA acquisition.

One candidate neuromodulator is acetylcholine (ACh). We recently observed that many PB*^Calca^* neurons express *Chat*^27^, suggesting they may also release ACh^36^. Whether PB*^Calca^* neurons are functionally cholinergic is not known, and the role of PB-derived ACh in CeA in associative learning has received limited attention^36,37^. In the CeA, ACh binding at G_q_-linked muscarinic receptors could increase excitability of these neurons on an extended timescale, bridging novel flavour experience and the delayed visceral malaise that drives CFA learning. ACh is important for various forms of associative learning^38,39^, and drug infusion studies in insular cortex and amygdala suggest ACh plays a role in CFA acquisition^40–42^. We hypothesised that a functional cholinergic projection from PB facilitates associative plasticity in CeA to produce CFA learning.

## Results

### ACh is released in the CeA in response to novel flavour and visceral malaise

We first sought to observe endogenous ACh release dynamics in CeA across stages of CFA learning. We adapted CFA to a head-fixed paradigm using an open-source system^43^. This head-fixed approach employs a “multi-spout” system, wherein solution-dispensing spouts rotate, extend, and retract from the mouse, producing rich, trial-structured behaviour that facilitates repeated sampling of multiple solutions in a session. Following water-only habituation sessions, mice underwent a conditioning session in which they had access to a novel solution (sweetened grape Kool-Aid; conditioned stimulus, CS). Afterward, mice received either a malaise-inducing lithium chloride (LiCl) injection^44^ or a control saline injection. After a recovery day, mice underwent another round of conditioning and recovery^30^. Finally, on Test days, mice received an equal number of water and CS trials (**Fig. 1a-c**). Preference was measured by dividing the number of licks for the CS by total licks, as well as by the last-lick time on CS trials, since mice with CFAs are expected to terminate licking for the CS early relative to controls^45^. We observed that LiCl-conditioned mice formed robust CFAs that persist even 3 weeks after conditioning (**Fig. 1d-g, Extended Data Fig. 1**), consistent with traditional, freely moving CFA models^30^.

**Fig. 1.**
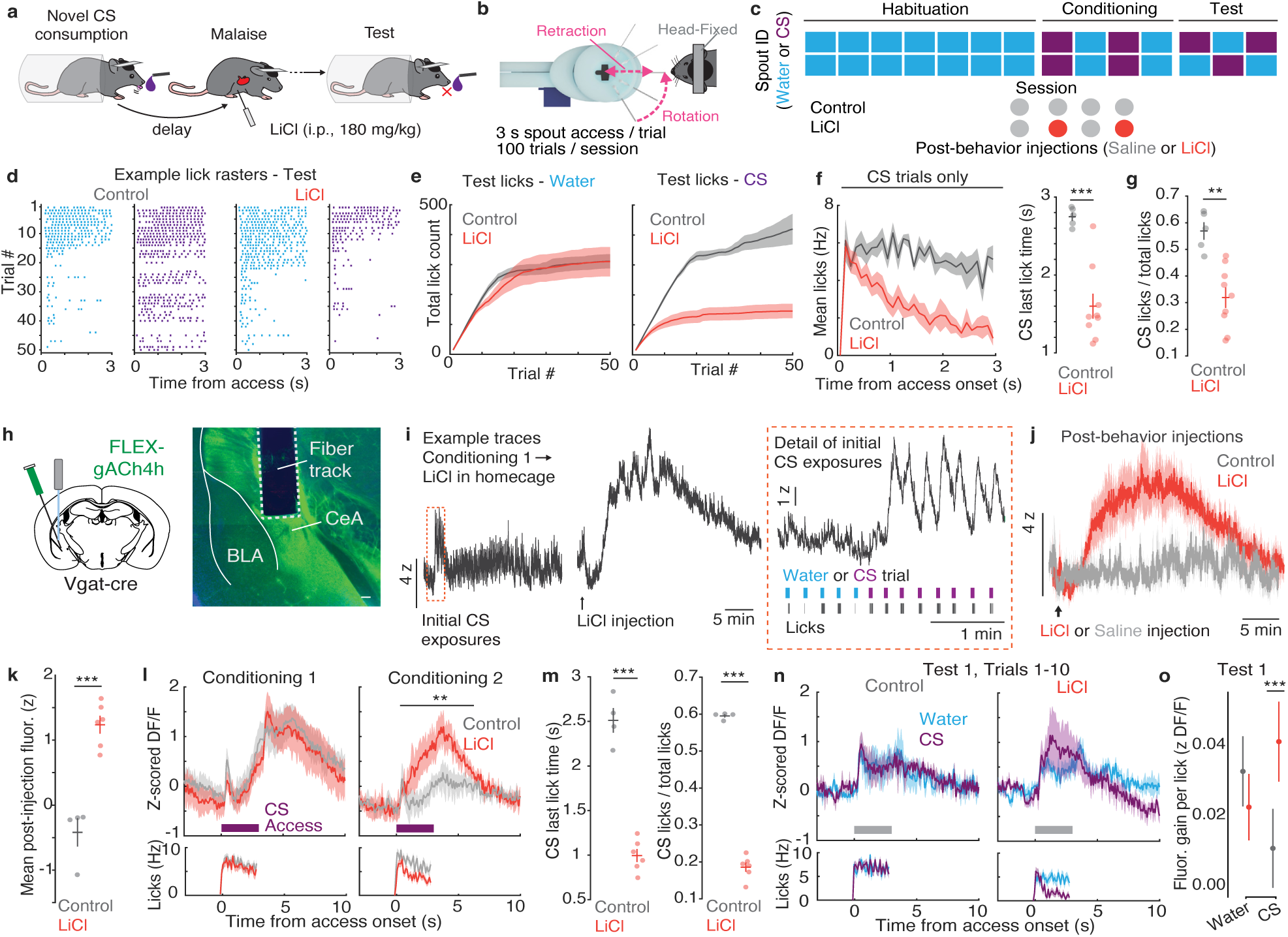
ACh is released in CeA at key timepoints of CFA acquisition. **a,** Schematic of experimental design for head-fixed CFA. **b,** Schematic of multi-spout system. **c,** Timeline of CFA experiment. **d,** Example lick raster plots during Test for example Control (left) and LiCl (right) mice. **e,** Mean cumulative licks for Water and CS during the Test session. **f,** Left: Mean lick rate during the 3-s access period for CS trials at Test. Right: Mean last-lick time during CS trials at Test. **g,** CS preference at Test (n = 5 Control; 9 LiCl). **h,** Left: Viral schematic for ACh photometry in CeA. Right: Example ACh sensor expression and photometry fibre placement. Scale bar is 100 µm. **i,** Left: Example traces during initial Conditioning session and subsequent LiCl injection in a LiCl mouse. Right: Detail of Conditioning trace showing initial CS exposures. **j,** Mean ACh sensor fluorescence after injection of LiCl (LiCl, n = 6) or saline (Control, n = 4). **k,** Mean post-injection fluorescence by group. **l,** Top: Mean ACh sensor fluorescence during first 10 CS presentations on Conditioning 1 (left) and 2 (right). Bottom: Mean lick rates. **m,** Left: Last-lick time during CS trials at Test. Right: CS preference at Test. **n,** Top: Mean ACh sensor fluorescence during water and CS trials for Control (left) and CFA (right) mice. Bottom: Mean lick rate. Gray bar indicates access period. **o,** ACh sensor fluorescence gain per lick for water and CS for data in **n**. Bars and shaded regions represent SEM, except for o, where bars represent SD. \**P*<0.05, \*\**P*<0.01, \*\*\**P*<0.001. See **Supplementary Table 1** for details of statistical tests and exact *P* values.

To observe ACh dynamics in CeA across CFA, Vgat-cre mice were injected with a Cre-dependent ACh sensor (FLEX-gACh4h^46^), which effectively prevented expression in the adjacent, primarily glutamatergic lateral and basolateral amygdalae (**Fig. 1h**). Mice showed strong, phasic ACh release in response to the novel CS on initial exposures (**Fig. 1i**), consistent with experiments demonstrating that a novel food induced calcium transients in PB*^Calca^* neurons^28,34^. Strikingly, mice that received a LiCl injection after conditioning also showed a large, extended increase in ACh not seen in saline-injected mice (**Fig. 1j-k**). Interestingly, by the second conditioning day, saline-conditioned mice had significantly smaller responses to the CS during initial trials, indicating that as a flavour becomes familiar, phasic ACh release in CeA is attenuated (**Fig. 1l**), as has been observed in insular cortex^47^. On Test days, LiCl-conditioned mice displayed robust CFAs (**Fig. 1m**). While ACh release during CS trials at Test was modest and similar between LiCl-conditioned and control mice, when accounting for the difference in licking behaviour between groups, LiCl mice showed significantly more gain in ACh levels per lick than did controls (**Fig. 1n-o**).

Thus, using this head-fixed CFA paradigm, we observed that ACh is released in the CeA both in response to a novel solution and during visceral malaise, two key timepoints of CFA acquisition. These dynamics led us to hypothesize that ACh facilitates the CeA plasticity required for CFA.

### PB*^Calca^* neurons send a functional cholinergic projection to CeA

Prior single-cell RNA sequencing experiments revealed that many PB*^Calca^* neurons also express *Chat*, which encodes choline acetyltransferase (ChAT), the biosynthetic enzyme for synthesis of ACh, and that PB*^Calca^* and PB*^Chat^* neurons are both largely restricted to expression in the external lateral segment of the PB^27^. To visualize the overlap of PB*^Calca^*and PB*^Chat^* neurons and their projections, we used a double transgenic *Calca^Flp^*::*Chat^Cre^* mouse model to virally label both populations. We observed the canonical PB*^Calca^*projections to the capsular region of CeA, whereas PB*^Chat^* projections innervated CeA more broadly (**Fig. 2a**). Using a fluorescence *in situ* hybridization (FISH) approach, we observed that 57% of *Chat*+ PB neurons co-express *Calca* (**Fig. 2b**).

**Fig. 2.**
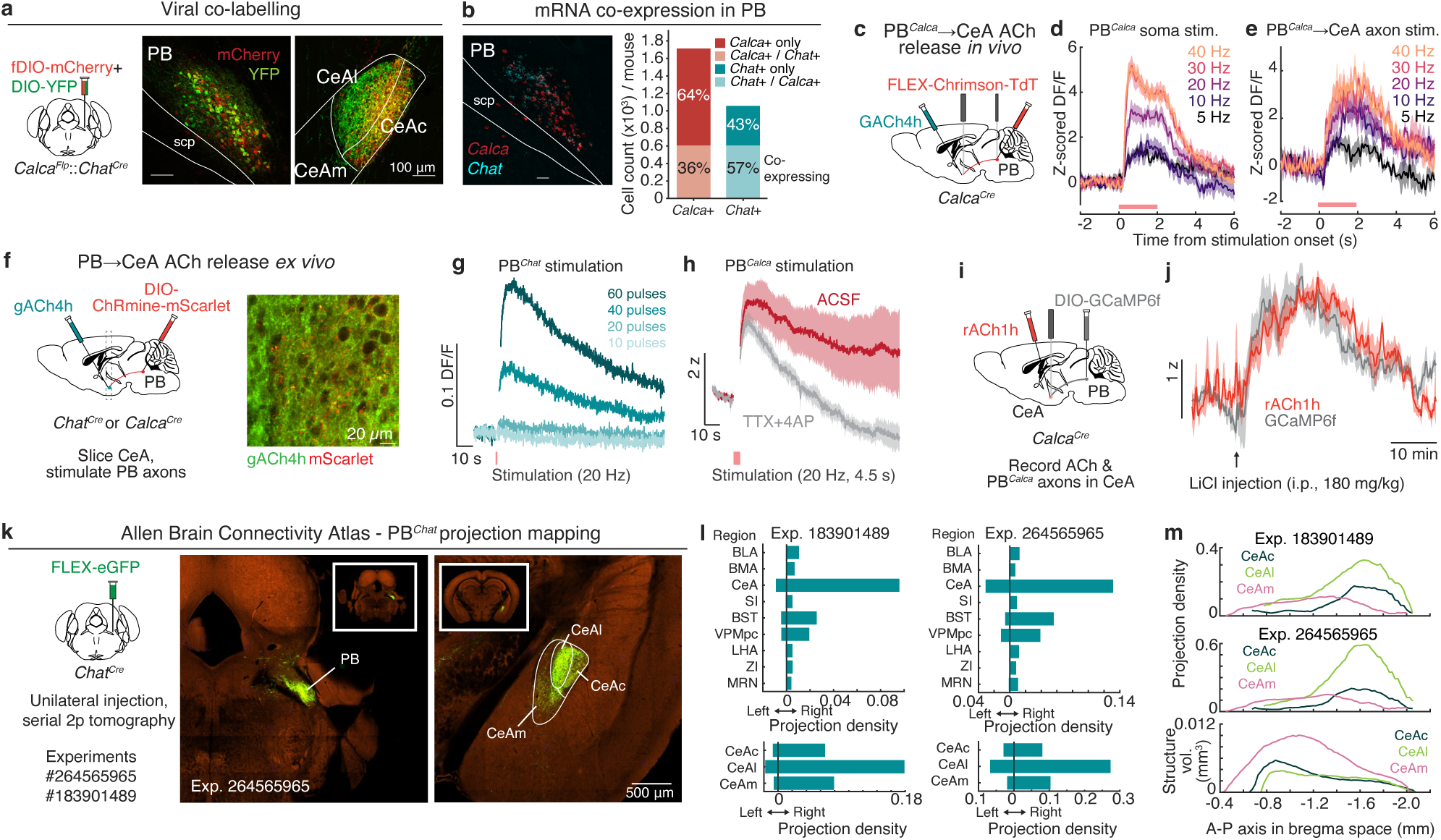
PB*^Calca^* neurons release ACh in CeA. **a,** Left: Viral approach for fluorescent labelling of PB *Calca*+ and *Chat*+ neurons. Middle: Overlap of *Calca*+ (mCherry) and *Chat*+ (YFP) neurons in PB. Right: *Calca*+ and *Chat*+ projections from PB in CeA, with approximate CeA subregions labeled (lateral, CeAl; capsular, CeAc; medial, CeAm). **b,** Left: Example image from FISH experiment. Right: Quantification of mean PB *Calca*+ and *Chat*+ cell counts per mouse, with shading indicating percentage of overlap between mRNA expression for a given marker (n=5 mice). **c,** Viral schematic for optogenetically evoked ACh release in CeA from PB*^Calca^*neurons. **d,** Mean ACh sensor response in CeA to 2 s pulsed stimulation at PB*^Calca^* cell bodies. **e,** Mean ACh sensor response in CeA to 2 s pulsed stimulation at PB*^Calca^* axons in CeA (n=2 mice). **f**, Left, Viral schematic for optogenetic stimulation of PB axons in CeA slices during 2-photon imaging. Right, Example image of ACh sensor and ChRmine-expressing PB*^Chat^* axons in CeA. **g**, ACh sensor response to 20 Hz stimulation of PB*^Chat^* axons across range of pulse counts. **h**, Mean ACh sensor response to PB*^Calca^* stimulation in ACSF and TTX+4AP (n=4 samples, 2 mice). **i,** Viral strategy for simultaneous photometry recording of ACh and PB*^Calca^* axon signalling. **j,** Average trace of simultaneous recording ACh and PB*^Calca^* axon activity following LiCl injection (n=6 mice). **k,** Left, Viral approach for anterograde tracing used in Allen Brain Connectivity experiments. Right, Example images of injection site and projections in CeA, with subregions drawn from corresponding Common Coordinate Framework atlas. **l,** Top, Projection density in select target areas for two separate experiments (BLA, basolateral amygdalar nucleus; BMA, basomedial amygdalar nucleus; SI, substantia innominata; BST, bed nucleus of the stria terminalis; VPMpc, ventral posteromedial nucleus of the thalamus, parvicellular part; LHA, lateral hypothalamic area; ZI, zona incerta; MRN, midbrain reticular nucleus). Bottom, Projection density by CeA subregion. **m,** Top and middle, Projection density by CeA subregion as a function of position on A-P axis in bregma space for either experiment. Bottom, Subregion structure as a function of position on A-P axis in bregma space for Common Coordinate Framework atlas. Shaded regions represent SEM.

Given previous findings that PB*^Calca^* activity during malaise and novel CS exposure are necessary for CFA acquisition^13,30,33,34^, we sought to determine whether PB*^Calca^* neurons release ACh in CeA. To test this, we injected a Cre-dependent, red-shifted excitatory opsin^48^ in the PB of *Calca^Cre^* mice and expressed an ACh sensor in CeA. Fibres were implanted over PB for optogenetic stimulation and over CeA for combined stimulation and ACh sensor photometry. Stimulation of either PB*^Calca^* cell bodies or their axons in CeA drove ACh release in CeA in a stimulation frequency-dependent manner (**Fig. 2c-e**). To rule out polysynaptic ACh release, we used a similar viral approach^49^ while performing 2-photon *ex vivo* imaging of ACh sensor fluorescence in CeA during optogenetic stimulation of PB axons (**Fig. 2f**). We confirmed PB*^Chat^* axon stimulation evoked ACh release with this approach (**Fig. 2g**). Stimulating PB*^Calca^*axons in presence of tetrodotoxin (TTX) and 4-aminopyridine (4AP) drove robust ACh release, demonstrating a functional, monosynaptic cholinergic projection to the CeA (**Fig. 2h**).

Since PB*^Calca^* neurons respond to noxious stimuli across modalities^28^, we reasoned that the same stimuli should drive ACh release in CeA. We used ACh sensor photometry to observe phasic ACh release in response to noxious thermal, mechanical, and electrical stimuli (**Extended Data Fig. 2**). Additionally, we injected viruses for a red-shifted ACh sensor^50^ in CeA and a Cre-dependent calcium indicator in the PB of *Calca^Cre^*mice to simultaneously record ACh release and PB*^Calca^* axon activity in CeA. Following injection of LiCl, we observed large, coincident increases in ACh and PB*^Calca^*axon calcium fluorescence (**Fig. 2i-j**).

Since the PB*^Calca^* projection had only partial overlap in CeA with the PB*^Chat^* projections (**Fig. 2a**), we used the Allen Brain Atlas Connectivity Map to better understand the projection pattern of PB*^Chat^* neurons (**Fig. 2k**). Two experiments with primary injection sites in the PB of *Chat^Cre^* mice both indicated that the CeA is the main target of these neurons, with notable other projections to the bed nucleus of the stria terminalis (BST) and ventral posteromedial nucleus (VPMpc; **Fig.2l**). In both experiments, all CeA subregions received projections, with the CeAl receiving the densest projection (**Fig. 2l**). We measured the density of the PB*^Chat^* projection as a function of anterior-posterior position in the CeA and found that PB*^Chat^* neurons projected primarily to the posterior portions of the CeAc and CeAl (**Fig. 2m**).

Together, these data establish that PB*^Calca^* neurons send a functional cholinergic projection to the CeA. Moreover, the broader PB*^Chat^* population sends dense projections to the CeA, particularly to the posterior CeAc and CeAl.

### ACh activates CeA neurons through muscarinic receptors

How does ACh affect CeA neuronal activity? The neuromodulatory role of ACh has been hypothesised to be mediated through a family of G_i_-or G_q_-coupled muscarinic receptors, which produce long-lasting, opposing effects on neuronal excitability^51^. Using single-nucleus RNA sequencing, we observed widespread expression of genes encoding G_q_-coupled M1, M3, and M5 muscarinic receptors in CeA (**Extended Data Fig. 3**), consistent with previous observations^52^. Thus, we reasoned that ACh may have a predominantly excitatory effect in CeA.

To test this, we injected Vgat-cre mice with a Cre-dependent calcium indicator and performed 2-photon *ex vivo* calcium imaging in the CeA (**Fig. 3a**), an approach that yielded calcium activity dynamics from hundreds of neurons per recording and over 10,000 neurons across experiments. Bath application of carbachol (Cch), a cholinomimetic drug, drove calcium transients in neurons throughout the CeA (**Fig. 3b-c**). Pre-treating slices with a cocktail of antagonists for glutamatergic receptors (NBQX, targeting AMPA receptors; AP5, targeting NMDA receptors) and nicotinic receptors (mecamylamine) did not attenuate the calcium signal. The general muscarinic antagonist scopolamine abolished this effect, as did the M3-and M5-preferring antagonists 4DAMP and VU 6019650^53^, implicating M3 and M5 muscarinic receptors as primary drivers of this activation (**Fig. 3d-e**).

**Fig. 3.**
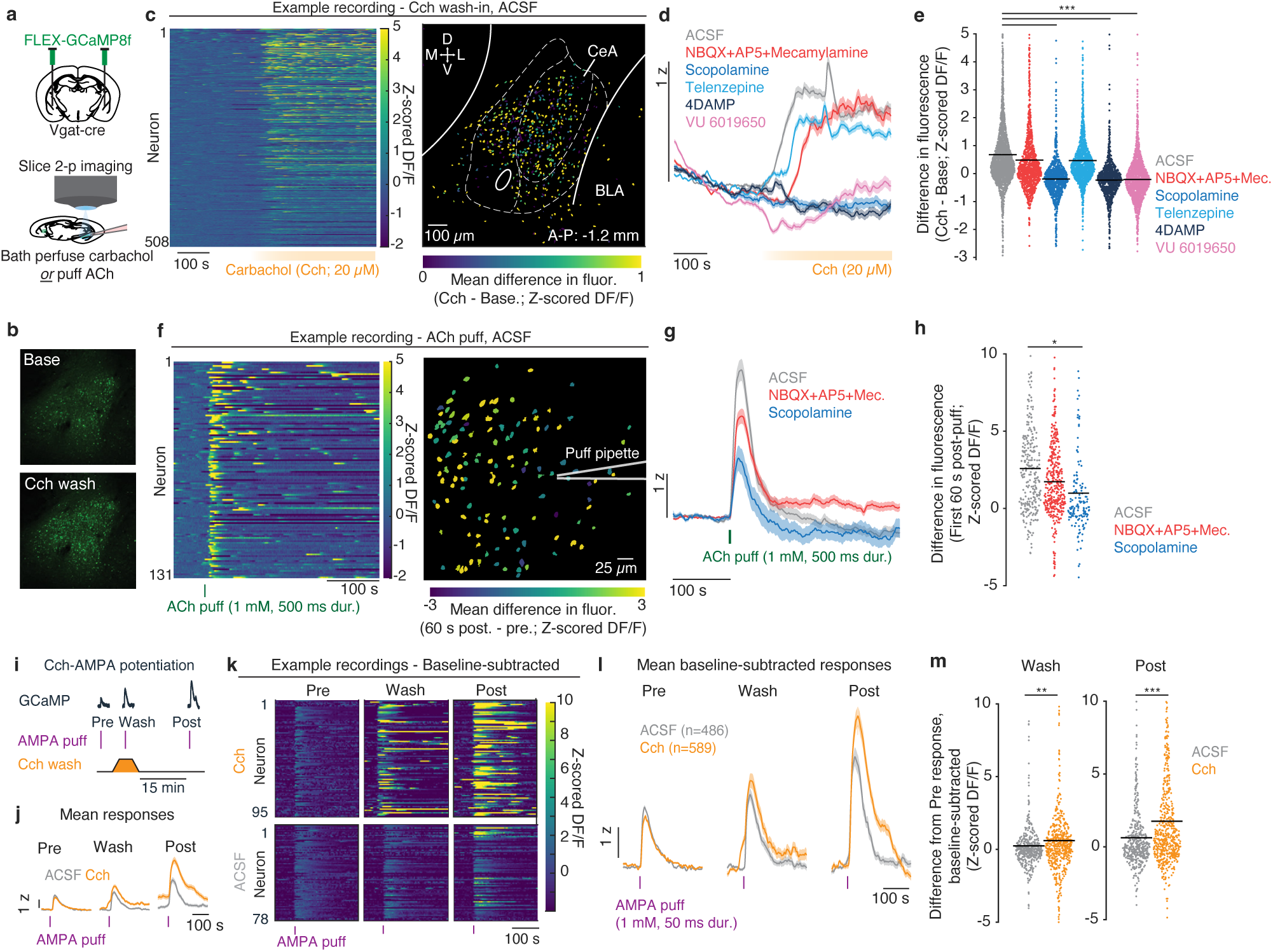
ACh drives widespread activation of CeA neurons and enhances glutamatergic responsiveness on a minutes-long timescale. **a,** Top: Schematic of approach for *ex vivo* 2-photon calcium sensor imaging in CeA. **b,** Mean fluorescence before (Top, “Base”) and after (Bottom) bath perfusion of carbachol (Cch). **c,** Left: Example heatmap of calcium sensor fluorescence during Cch wash-in experiment. Right: Example mean baseline-subtracted fluorescence during Cch wash by neuron. **d,** Mean fluorescence across all neurons by treatment group during Cch wash (ACSF, n=2154 neurons, 7 slices; NBQX (20 µM) + AP5 (50 µM) + Mecamylamine (10 µM), n=1197 neurons, 5 slices; Scopolamine (10 µM), n=1031 neurons, 5 slices; Telenzepine (0.1 µM), n=1482 neurons, 4 slices; 4DAMP (1 µM), n=956 neurons, 9 slices; VU 6019650 (1 µM), n=1664 neurons, 6 slices). **e,** Mean baseline-subtracted fluorescence during Cch wash by group. Bars denote significant effects for Scopolamine, 4DAMP, and VU 6019650. **f,** Left: Example heatmap of calcium sensor fluorescence during ACh puff experiment. Right: Example mean baseline-subtracted fluorescence following ACh puff. **g,** Mean fluorescence across all neurons by treatment around ACh puff (ACSF, n=274 neurons, 4 slices; NBQX+AP5+Mecamylamine, n=388 neurons, 3 slices; Scopolamine, n=153 neurons, 3 slices). **h,** Mean baseline-subtracted fluorescence 60 s following ACh puff. ACh puff experiments were performed in the presence of physostigmine (1 µM). **i,** Timeline for Cch-AMPA puff potentiation experiment. **j,** Non-baseline-subtracted mean calcium fluorescence before (Pre), during (Wash), and 15 min after (Post) Cch wash by group. **k,** Top: Example heatmaps of baseline-subtracted calcium sensor fluorescence response to AMPA puffs in Cch wash experiment. Bottom: Same as Top, but in ACSF control. **l,** Mean baseline-subtracted fluorescence across all neurons by treatment group. **m,** Difference in mean response to AMPA puff from Pre period during Wash (Left) and after wash-out (Post, Right) (ACSF, n=486 neurons, 10 slices; Cch, n=569 neurons, 9 slices). All experiments were performed in the presence of picrotoxin (100 µM). Shaded regions represent SEM. \**P*<0.05, \*\**P*<0.01, \*\*\**P*<0.001. See **Supplementary Table 1** for details of statistical tests and exact *P* values.

We observed phasic ACh signals in CeA *in vivo* in response to a novel solution (**Fig. 1i,l**). To test whether phasic ACh signalling was also sufficient to activate CeA neurons, we micro-puffed ACh onto slices. We found that a single micro-puff of ACh could drive robust increases in calcium fluorescence (**Fig. 3f**), an effect that was significantly attenuated by scopolamine, but not by a mix of glutamatergic and nicotinic antagonists (**Fig. 3g-h**). In separate patch-clamp recordings, bath-applied Cch or micro-puffed ACh drove action potentials in CeA neurons (**Extended Data Fig. 4a-b**), indicating that the observed calcium increases reflect spiking.

One mechanism for increasing neuronal excitability via muscarinic receptors is inhibition of M-currents, voltage-sensitive and non-inactivating potassium currents that effectively resist membrane depolarization near spike threshold^54,55^. With patch-clamp electrophysiology, we observed M-currents in CeA neurons that were inhibited by Cch (**Extended Data Fig. 4c-i**). Following muscarinic activation, in the absence of ACh, recovery of M-currents can take minutes^54^. These currents offer one channel-specific mechanism, among other possible mechanisms^56^, by which ACh can increase CeA excitability.

Additionally, exposure to a novel CS has been shown to activate protein kinase A (PKA) in CeA^13^, evidence of a biochemical eligibility trace that may promote later reactivation during malaise. Whether such PKA activation could be due to a muscarinic mechanism is unclear: the canonical G_q_ signalling pathway does not involve PKA, but G_q_-linked muscarinic receptors activate PKA in the hippocampus^57^. To test whether muscarinic activation can stimulate PKA activity in CeA, we injected Vgat-cre mice with a Cre-dependent PKA sensor (DIO-ExRai-AKAR2^58^) in CeA and performed *ex vivo* 2-photon PKA sensor recordings in the presence of TTX. Carbachol increased PKA sensor fluorescence, and this effect was blocked by scopolamine (**Extended Data Fig. 5**), indicating that muscarinic receptors can drive non-canonical PKA activation in CeA.

Together, these experiments reveal a previously unreported capacity for ACh to broadly increase CeA neuronal excitability through Gq-linked muscarinic receptors, including the rare M5 subtype^59^. The combined results indicate that PB-derived ACh increases CeA neuronal excitability at key timepoints of CFA acquisition.

### ACh enhances glutamatergic responsiveness on a timescale consistent with CFA

Associative plasticity underlying CFA must bridge activity at the CS consumption and malaise stages, which occur minutes to hours apart. We reasoned that ACh could increase excitability and potentiate glutamatergic responsiveness of CeA neurons on a minutes-long timescale, which would provide a means of linking activity at consumption and malaise to facilitate plasticity.

To test this idea, we performed an additional set of *ex vivo* 2-photon calcium imaging experiments where CeA neurons received micro-puffs of α-Amino-3-hydroxy-5-methyl-4-isoxazolepropionic acid (AMPA) intended to generate suprathreshold responses detectable via calcium imaging. AMPA puffs were delivered before, during, and 15 minutes after a brief exposure to Cch (**Fig. 3i-k**). To measure the effect of Cch on each neuron’s response to AMPA puff, we compared the difference between its baseline-subtracted response in the pre-Cch period (“Pre”) to its response during Cch wash-on (“Wash”) and 15 min after wash-out^12^ (“Post”) periods. Cch-treated neurons showed a significant increase in AMPA response at Wash and that persisted at least 15 min later, as measured at the Post stage (**Fig. 3l-m**), consistent with reports in other brain regions that muscarinic activation is sufficient to enhance glutamatergic responsiveness on an extended timescale^60–62^. In the context of our CFA study, this result supports the possibility that ACh released onto CeA neurons during novel CS consumption could support later reactivation of these neurons during visceral malaise, providing a GPCR-based mechanism for associative plasticity.

### PB→CeA ACh facilitates CFA

Based on the previous results, we hypothesised that ACh released from PB neurons may be necessary for CFA learning. To test the importance of ACh release from PB neurons, we used *Chat^Cre^*mice to target the entire PB *Chat*+ population, reasoning that *Chat*+/*Calca*-neurons may also participate in CFA. Mice were injected in the PB with Cre-dependent CRISPR-Cas9 viral constructs targeting *Slc18a3*, which encodes the vesicular acetylcholine transporter (VAChT), and *Chat*, which encodes choline acetyltransferase (ChAT) (sg*Chat*+). In unilaterally injected mice, we observed qualitatively fewer ChAT+ PB cell bodies on the side ipsilateral to sg*Chat*+ injection (**Fig. 4a**).

**Fig. 4.**
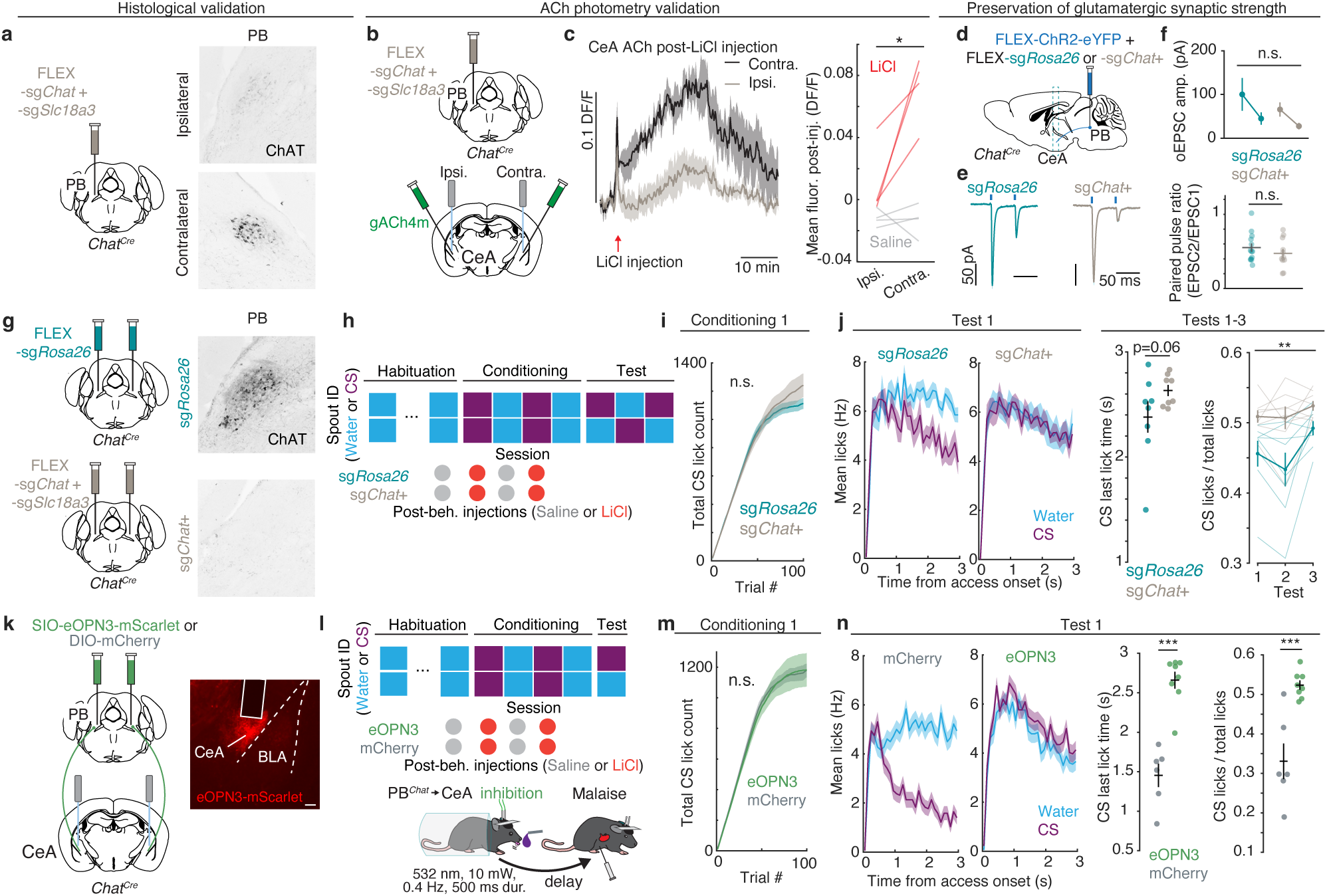
PB^ACh^ and PB*^Chat^*→CeA input facilitate CFA acquisition. **a,** Left: Viral strategy for unilateral knockdown of *Chat* and *Slc18a3* in PB*^Chat^* neurons. Right: Example ChAT expression in PB ipsi- and contralateral to site of viral injection. **b,** Viral strategy for bilateral ACh photometry for functional validation of *Chat* and *Slc18a3* knockdown in PB. **c,** Left: Mean ACh sensor fluorescence during LiCl injection in CeA ipsilateral (Ipsi.) and contralateral (Contra.) to site of knockdown viral injection. Right: Mean post-injection ACh sensor fluorescence by recording site and injection type (n=4 mice). **d,** Viral strategy for validation of preservation of glutamatergic signalling following sg*Chat*+ knockdown. **e,** Example optically evoked excitatory postsynaptic currents (oEPSCs) at CeA neurons in response to paired-pulse PB*^Chat^* axon stimulation. **f,** Top: Mean oEPSC amplitudes by group during first and second pulse. Bottom: Mean paired-pulse ratio (PPR) by group (sg*Rosa26*, n=13 neurons, 2 mice; sg*Chat*+, n=11 neurons, 2 mice). **g,** Viral strategy for bilateral knockdown of *Chat* and *Slc18a3* in PB*^Chat^* neurons for CFA behavioural testing. **h,** Timeline of CFA behavioural experiment. **i,** Mean cumulative licks for CS by group during the first conditioning session. **j,** Left: Mean lick rates during lick trials by solution and group during Test 1. Middle: Last-lick time on CS trials during Tests 1-3. Right: CS preference by group across Tests 1-3 (sg*Rosa26*, n=8 mice; sg*Chat*+, n=8 mice). **k,** Left: Viral strategy for inhibition of PB*^Chat^*terminals in CeA during CFA acquisition. Right: Example histology and fibre placement in CeA. **l,** Top: Timeline of CFA terminal inhibition experiment. Bottom: Detail of laser delivery timing during Conditioning sessions. **m,** Mean cumulative lick for CS by group during the first conditioning session in the terminal inhibition experiment. **n,** Left: Mean lick rates during lick trials by solution and group during Test. Middle: Last-lick time on CS trials during Test. Right: CS preference during Test (mCherry, n=6 mice; eOPN3, n=8 mice). Shaded regions represent SEM. \**P*<0.05, \*\**P*<0.01, \*\*\**P*<0.001. See **Supplementary Table 1** for details of statistical tests and exact *P* values.

To confirm whether this approach reduces ACh released in CeA during malaise, we performed unilateral injections of sg*Chat*+ in the PB and injections of an ACh sensor in CeA for bilateral fibre photometry measurements (**Fig. 4b**). While saline injection had no effect on ACh signal in CeA, we observed LiCl-driven ACh release in the CeA that was largely abolished at the CeA site ipsilateral to the PB sg*Chat*+ injection (**Fig. 4c, Extended Data Fig. 6a-c**), demonstrating that the CRISPR approach is sufficient to suppress ACh release during visceral malaise. We also measured locomotion in an open-field chamber before and after acute LiCl injection, as reduced locomotor activity is an established indicator of malaise^63^. We observed that sg*Chat*+ mice locomoted significantly more than did controls following LiCl injection (**Extended Data Fig. 6d-g**). Importantly, glutamatergic transmission from PB to CeA neurons was not affected by CRISPR treatment (**Fig. 4d-f**).

Having established that sg*Chat*+ treatment reduces ACh release from PB neurons, we tested whether PB^ACh^ was required for CFA behaviour. Using *Chat^Cre^* mice, we bilaterally injected either the Cre-dependent sg*Chat*+ virus mixture or a Cre-dependent control virus targeting the non-coding *Rosa26* gene^64^ (sg*Rosa26*; **Fig. 4g**). All mice received LiCl injections following conditioning sessions (**Fig. 4h**). Viral ACh knockdown had no effect on CS consumption during the initial conditioning session (**Fig. 4i**); however, following conditioning, across Test sessions, sg*Chat*+ mice licked for the CS significantly more compared to sg*Rosa26* controls (**Fig. 4j**), indicating that PB^ACh^ facilitates CFA behaviour.

Because this CRISPR approach permanently attenuates ACh release by PB*^Chat^* neurons and, thus, cannot reveal when PB*^Chat^* neuron activity is important for behaviour, or whether the PB*^Chat^* projection to the CeA is specifically involved, we performed a complementary projection-specific optogenetic inhibition experiment. We injected a Cre-dependent inhibitory opsin specialized for terminal inhibition (SIO-eOPN3-mScarlet^65^) bilaterally into the PB of *Chat^Cre^* mice and implanted optical fibres over CeA (**Fig. 4k**). Control mice received injections of a Cre-dependent fluorophore-only virus (DIO-mCherry). We administered brief pulses of laser throughout the head-fixed CS consumption period of both Conditioning sessions to inhibit PB*^Chat^* terminals during CS consumption. We then injected LiCl 5 to 7 min after termination of the conditioning session to reduce the likelihood that eOPN3 inhibition persisted into the malaise period (**Fig. 4l**). Inhibition of PB*^Chat^***→**CeA terminals did not affect CS consumption during the first conditioning session (**Fig. 4m**). However, at the Test, mice that received PB*^Chat^***→**CeA inhibition during CS consumption showed significantly longer latency to terminate licking for the CS and a greater preference for the CS (**Fig. 4n**), indicating that PB*^Chat^***→**CeA activity during novel CS exposure contributes to CFA acquisition.

Thus, knocking down PB^ACh^ release or inhibiting PB*^Chat^***→**CeA activity during CS exposure attenuates acquisition of CFA. Together, these results establish that PB^ACh^→CeA facilitates CS-malaise associative learning.

### PB^ACh^→CeA promotes reactivation of CS-preferring neurons in malaise

We next asked how loss of PB^ACh^ input affects the neural correlates of CFA in CeA. Associative plasticity in CFA likely requires neurons responsive to a novel CS to be selectively reactivated during later malaise experience. At Test, such plasticity would result in enhanced neuronal responses to the CS. We hypothesized that PB^ACh^ would be required for CS-malaise plasticity during acquisition, and that CeA neurons would fail to develop enhanced CS responses in its absence.

To record from CeA neurons across CFA behaviour, we injected a calcium indicator in the CeA and implanted newly developed prism lenses^66^, which permit longitudinal, deep-brain recording of hundreds of neurons per mouse. In one experiment (“Base”), we injected Vgat-cre mice with a Cre-dependent calcium indicator to help restrict expression to the CeA. In another experiment (“CRISPR”), using *Chat^Cre^* mice, we injected Cre-dependent sg*Rosa26* or sg*Chat*+ viruses bilaterally into the PB, and used a pan-neuronal calcium indicator in the CeA (**Fig. 5a**). In the Base experiment, mice received either saline (“Control”) or LiCl injections after CS conditioning sessions. In the CRISPR experiment, all mice received LiCl injections (**Fig. 5b**). After these experiments, we registered coronal sections containing the imaging plane to a reference atlas, permitting identification of CeA neurons and their membership in the capsular (CeAc), lateral (CeAl), or medial (CeAm) subregions of CeA (**Fig. 5c-d, Extended Data Fig. 7**).

**Fig. 5.**
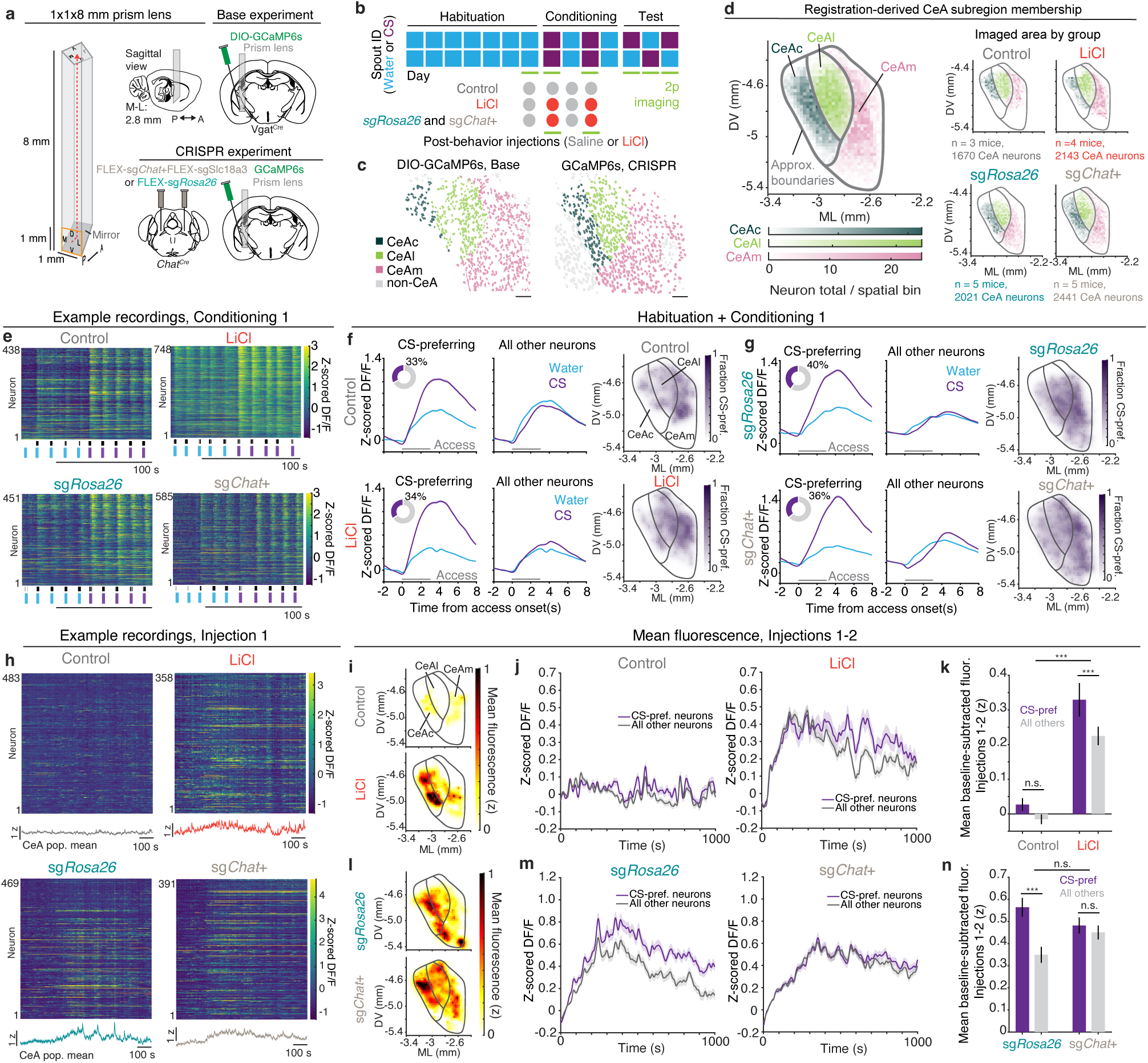
PB^ACh^ promotes preferential reactivation of CS-preferring neurons during malaise. **a**, Schematic of microprism lens, and viral and implant strategy for Base and CRISPR experiments. **b,** Timeline of 2p CFA experiments. **c,** Example fields-of-view with neurons coloured by CeA subregion, as determined through post-hoc atlas registration procedure in **Extended Data** Fig. 7**. d**, Left, Number of total imaged neurons per spatial bin in bregma space across all groups, colour-coded by CeA subregion. Right, Imaged neurons per spatial bin for each group. **e,** Example fluorescence heatmaps during Conditioning 1. Blue and purple horizontal lines denote periods of water or CS access. Vertical black ticks denote lick times. **f,** Left: Peristimulus time histograms for water and CS trials for CS-preferring and all other CeA neurons by group over Habituation and Conditioning 1 sessions in Base experiment. Inset donut represents proportion of CS-preferring neurons by group. Right: Mean fraction of CS-preferring neurons in bregma space. **g**, Same as **f**, but for CRISPR experiment. **h,** Example fluorescence heatmaps during Injection 1. Bottom line plot depicts mean population trace for the recording. **i**, Mean post-injection fluorescence during injections 1 and 2 in bregma space. **j,** Mean fluorescence during injection traces across CS-preferring and all other neurons in the Base experiment. **k**, Mean fluorescence by neuron class and group in the Base experiment (Control, n=3 mice, 1670 CeA neurons; LiCl, n=4 mice, 2143 CeA neurons). **l-n**, Same as **i-k**, for the CRISPR experiment (sg*Rosa26*, n=5 mice, 2021 CeA neurons; sg*Chat*+, n=5 mice, 2441 CeA neurons). Bars and shaded regions represent SEM. \**P*<0.05, \*\**P*<0.01, \*\*\**P*<0.001. See **Supplementary Table 1** for details of statistical tests and exact *P* values.

During the initial conditioning session, across groups, we observed robust neuronal responses to the CS (**Fig. 5e**). Using a linear model, we identified CS-preferring CeA neurons and found a similar fraction of CS-preferring neurons across groups, with CS-preferring neurons in each CeA subregion (**Fig. 5f**). This was true even for neurons in sg*Chat*+ mice, indicating that knockdown of PB^ACh^ input does not diminish overall CeA neuronal responsiveness to the CS (**Fig. 5g**). Following LiCl injection, we observed widespread increases in calcium fluorescence (**Fig. 5h**), particularly in the CeAc, to which PB*^Calca^* neurons send a robust projection^30^, and the CeAl (**Fig. 5i, Extended Data Fig. 8a-b**). In the Base experiment, when comparing the post-injection activity of identified CS-preferring neurons against all other neurons, we observed that these CS-preferring neurons underwent preferential reactivation during malaise (**Fig. 5j-k**). While this effect held true in LiCl-injected sg*Rosa26* control mice, sg*Chat*+ treatment significantly diminished the preferential reactivation of CS-preferring neurons (**Fig. 5l-n**). Together, these experiments reveal that the PB^ACh^ input facilitates preferential reactivation of neurons that were responsive to the novel CS, a key neural correlate of CS-malaise associative learning in CFA.

### PB^ACh^→CeA facilitates CFA-associated plasticity

We hypothesised that CeA neurons in LiCl-conditioned mice would have enhanced responses to the CS during Test sessions, and that sg*Chat*+ treatment would diminish these effects. At Test, LiCl-conditioned Control and sg*Rosa26* mice exhibited strong CFAs, whereas the sg*Chat*+ group did not (**Fig. 6a-b**), consistent with the behaviour-only experiment (**Fig. 4j**). In response to the CS, neurons in the LiCl group had an earlier peak time, despite marked decreases in licking (**Fig. 6c**). To measure how responses to the CS changed following conditioning, we compared peak statistics of each neuron at Conditioning 1 and Test 1 (**Fig. 6d**) and found that LiCl conditioning significantly decreased the time-to-peak (**Fig. 6e**) and increased the slope-to-peak (**Fig. 6f**) in response to the CS relative to Controls.

**Fig. 6.**
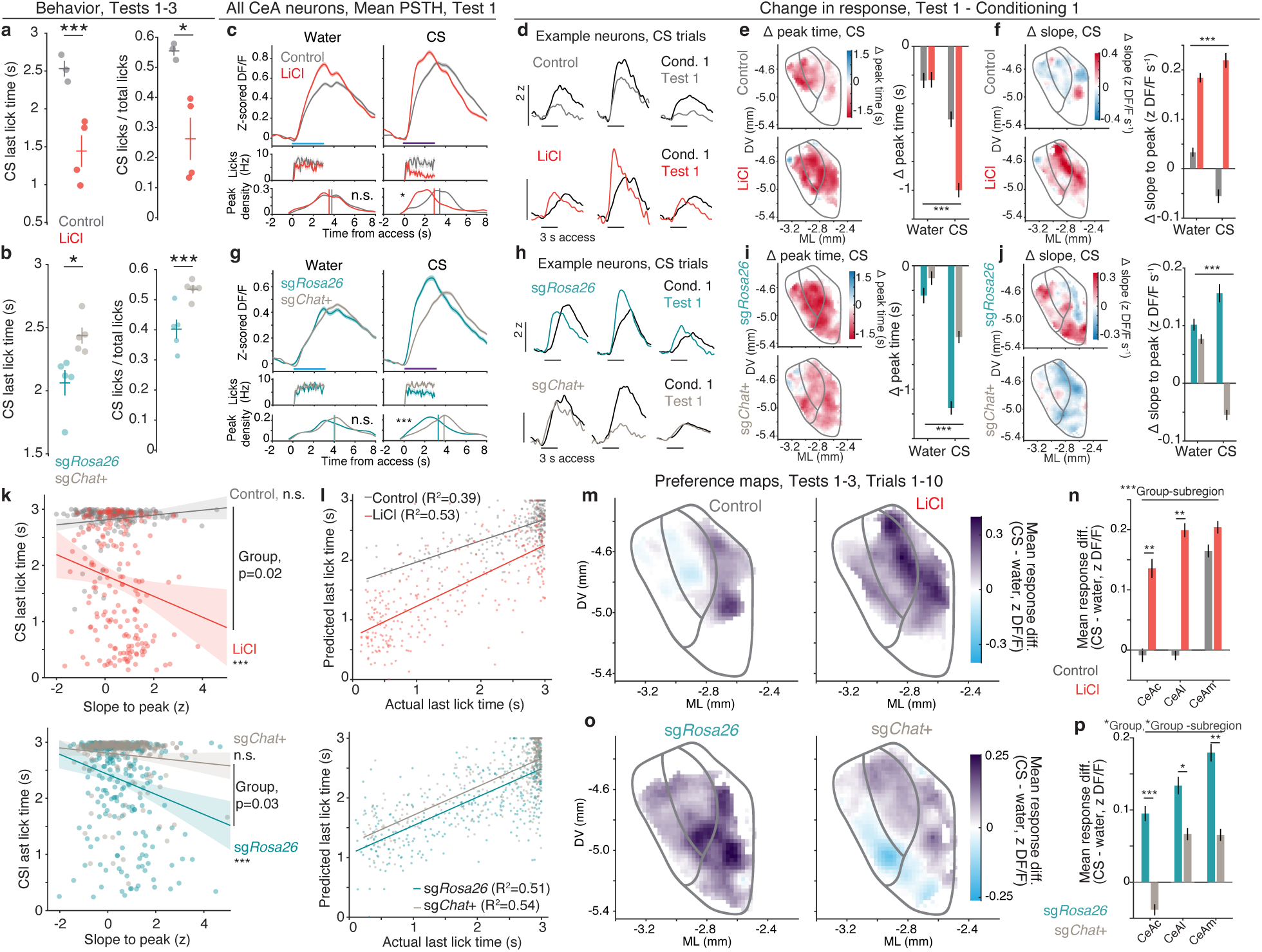
PB^ACh^ is required for CFA-associated plasticity in CeA. **a,** Left, Mean last-lick time on CS trials across Tests 1-3 in the Base experiment. Right, CS preference in the Base experiment. **b**, Same as **a**, for CRISPR experiment. **c,** Top, Mean neural responses across all CeA neurons during water and CS lick trials at Test 1 in Base experiment. Middle, Mean lick rates during lick trials at Test 1. Bottom, Distribution of mean peak time across neurons. **d,** Example neurons’ mean CS responses at Conditioning 1 (black) and Test 1 (group colour). Horizontal bars represent access periods. **e,** Left, Median change in neuronal peak times from Conditioning 1 to Test 1 by spatial bins. Right, Group mean change in peak time from Conditioning 1 to Test 1 in Base experiment. f, Left, Median change in slope-to-peak from Conditioning 1 to Test 1 by spatial bins. Right, Group mean change in slope-to-peak from Conditioning 1 to Test 1 in Base experiment. **g,** Top, Mean neural responses across all neurons during water and CS lick trials at Test 1 in CRISPR experiment. Middle, Mean lick rates during lick trials at Test 1. Bottom, Distribution of mean peak time across neurons. **h,** Example neurons’ mean CS responses at Conditioning 1 (black) and Test 1 (group colour). Horizontal bars represent access periods. **i,** Left, Mean change in neuronal peak times from Conditioning 1 to Test 1 by spatial bins. Right, Group mean change in peak time from Conditioning 1 to Test 1 in CRISPR experiment. **j,** Left, Mean change in slope-to-peak from Conditioning 1 to Test 1 by spatial bins. Right, Group mean change in slope-to-peak from Conditioning 1 to Test 1 in CRISPR experiment. **k,** Top, Scatter plot of z-scored slope-to-peak of population PSTH vs last-lick time on CS in Base experiment. Bottom, Same, for CRISPR experiment. **l,** Actual last-lick time vs predicted last-lick time using supervised dimensionality reduction approach on neural data in Base (top) and CRISPR (bottom) experiments. **m,** Spatial maps of group mean difference in neuronal response to CS versus water (from 0 to 1.5 s after access onset) during trials 1-10 on Tests 1-3 in Base experiment. **n,** Group mean difference in neuronal response to CS versus water by CeA subregion in Base experiment (Control, n=3 mice, 1670 CeA neurons; LiCl, n=4 mice, 2143 CeA neurons). **o-p**, Same as **m-n**, but for CRISPR experiment (sg*Rosa26*, n=5 mice, 2021 CeA neurons; sg*Chat*+, n=5 mice, 2441 CeA neurons). Shaded regions and bars represent SEM. \**P*<0.05, \*\**P*<0.01, \*\*\**P*<0.001. See **Supplementary Table 1** for details of statistical tests and exact *P* values.

In the CRISPR experiment, in CS trials at Test, we also observed a significant difference in time-to-peak between neurons in sg*Rosa26* and sg*Chat*+ groups, with a relative lag in peak in sg*Chat*+ neurons (**Fig. 6g**). Neurons in the sg*Rosa26* group had a pronounced decrease in time-to-peak and increase in slope-to-peak during CS trials from Conditioning 1 to Test 1, like what was observed in LiCl-conditioned mice in the Base experiment. This effect was significantly attenuated in the sg*Chat*+ group (**Fig. 6h-j**). Together, these data show that PB^ACh^ knockdown limits the plasticity of CeA neurons’ responses to CS following LiCl conditioning.

As calcium indicator fluorescence can be conceptualized as a leaky integration of underlying firing, we reasoned that this increase in slope-to-peak in LiCl and sg*Rosa26* mice represented increased firing rates in CeA neurons upon initial exposure to the CS at Test^67^. Given the CeA’s involvement in the regulation and termination of consumption^17,21–24,68^, we wondered whether a robust initial response to the CS may reflect detection of danger and promote the early termination of CS licking observed in our CFA experiment (**Fig. 6a,b**). To test this, we calculated the population mean peri-stimulus time histogram on a trial-by-trial basis and related the slope of these population responses to the last-lick times. We found that population slope-to-peak predicted last-lick time on CS trials in LiCl and sg*Rosa26* groups, but not in Control or sg*Chat*+ groups (**Fig. 6k**), consistent with the idea that an enhanced early CeA response to CS could be involved with early termination of licking.

To further test the idea that CeA activity is related to lick termination, for each mouse, we used a supervised dimensionality reduction approach to produce a “last-lick time (LLT) axis” score, which was used to predict the timing of the last lick using only neuronal data on a trial-by-trial basis. For every mouse, LLT axis-based estimates far outperformed the same analysis applied to trials with shuffled true last-lick times (**Extended Data Fig. 9d**) and produced similarly accurate estimates across experimental groups (**Fig. 6l**), indicating that CeA neuron activity can be used to predict lick termination.

Finally, we asked whether knockdown of PB^ACh^ affected the selectivity of CeA neuronal responses to the CS at Test. To test this, we measured the difference in each CeA neuron’s mean fluorescence during lick trials for either solution over the first 10 trials of Test sessions. In the Base experiment, we found significantly stronger CS preference in LiCl-conditioned mice than controls in the CeAc and CeAl (**Fig. 6m-n**). Interestingly, many neurons in the CeAm in both groups exhibited preferential responses to the CS. In the CRISPR experiment, we found significantly weaker CS preference throughout the CeA in the sg*Chat*+ group relative to the sg*Rosa26* group (**Fig. 6o-p**, **Extended Data Fig. 10**). Together, these data provide further evidence that, in the absence of PB^ACh^ input, CeA neurons do not undergo the plasticity processes that produce prototypical CFA-associated CS responses at Test.

Together, these experiments show that CeA neurons exhibit enhanced responses to the CS following its learned association with malaise. Following conditioning, many neurons fire robustly in immediate response to the CS, and this activity predicts early termination of CS consumption. These changes are significantly attenuated when PB^ACh^ is knocked down, consistent with our hypothesis that PB^ACh^→CeA is critical for plasticity processes relating CS consumption to malaise and producing CFA behaviour.

## Discussion

Associative learning is the basis for flexible, adaptive behaviour. Causally linked events are sometimes separated by considerable time gaps. How do neural systems bridge such gaps to promote learning? Here, using CFA, a rapid, strong, and phylogenetically widespread form of learning, we show that ACh, released from PB neurons into the CeA, is a key facilitator of this process. Previous work established the PB**→**CeA circuit in CFA acquisition^30^ and proposed that GPCR signalling in CeA may link novel consumption to malaise^13^; our results identify ACh as a neuromodulator that bridges this gap.

Neuromodulators can prime the induction of associative plasticity^12^. We propose that ACh, binding G_q_-linked muscarinic receptors on CS-responsive CeA neurons during consumption, establishes a functional eligibility trace that increases the likelihood of these neurons being reactivated during subsequent malaise, creating conditions for CS-malaise plasticity. Consistent with this idea, brief cholinergic treatment enhanced CeA responsiveness to excitatory input for at least 15 min, a timescale consistent with our CFA experiments. In the absence of PB-derived ACh, CFA acquisition is impaired and CeA neurons do not exhibit CFA-associated plasticity.

The large ACh release observed following LiCl injection implies an additional role of ACh in plasticity induction during malaise. That PB^ACh^ knockdown did not affect overall CeA neural activation during malaise suggests that the role of ACh is not a simple increase in excitatory drive. One possibility is that ACh during malaise also acts at G_i_-coupled muscarinic receptors to inhibit glutamatergic terminals in CeA, as described in striatum^69^. Under this model, PB^ACh^ released during malaise could filter excitatory input to CeA, reducing the likelihood that CFA-unrelated sensory information is associated with malaise. Likewise, net loss of PB^ACh^ during malaise would not yield less activity overall, but less organized, CFA-specific activity.

Our results are consistent with the literature showing that activation or inhibition of PB*^Calca^* neurons can promote or inhibit CFA formation^13,30,33,34^, as either manipulation would similarly affect PB^ACh^ release in CeA. The observation that PB*^Calca^* neurons can have heterogeneous responses to consumption and malaise^34^ raises the question of whether PB*^Chat^* neurons are similarly heterogeneous, or whether they are activated by both novel CS and malaise.

Prior research on CFA pursued the idea that flavour memory resided in the taste pathway, e.g., within the insular cortex, and that coincidence of the persistent taste signal and the malaise signal occurred in the basolateral amygdala^70,71^. However, the role of the PB*^Calca^* neurons, which do not project directly to the basolateral amygdala, was unknown then, and the idea that the novel CS signal and the malaise signal could arise from the same population of PB neurons was not considered. Moreover, ablation of the gustatory thalamus does not block CFA^72^.

While our data indicate that ACh is critical for CFA acquisition, other neuromodulators are likely involved. For example, the CeA receives dopaminergic innervation from the ventral tegmental area and substantia nigra pars compacta and has widespread expression of dopamine receptors^17,73^. Our finding that muscarinic receptor activation can stimulate PKA activity alludes to the possible convergence of dopamine, acting through D1 receptors^74^, and ACh signals during novel CS consumption to facilitate plasticity during later malaise. Additionally, PB*^Calca^*neurons express neuropeptides (CRH, TAC1, NTS, and PACAP) that can also activate G_q_-linked GPCRs; however, they would be in dense-core vesicles, whereas ACh is transported into synaptic vesicles. Because the contents of synaptic vesicles are preferentially released at low frequencies, ACh is likely to be released to a greater extent by weaker CS-mediated neuronal activation than neuropeptides^75^. Given that neuropeptides can remain in the extracellular space for extended time periods, they may be important for CFAs when the CS-malaise interval is hours-long, which we did not examine here. How neuromodulators work in concert in CFA remains an open question.

Because it has been suggested that neither canonical genetic markers for CeA cell types nor CeA subregions reliably capture all CFA-involved CeA neurons^13^, we used a pan-neuronal approach in our calcium imaging studies to image CeA neurons. Our *in vivo* imaging data suggest that many CeAc and CeAl neurons have enhanced responses to a CS following LiCl conditioning, an effect that is diminished with PB^ACh^ knockdown. Our ability to predict lick termination with CeA activity suggests that, consistent with literature, a subset of CeA neurons may be causally involved in early lick termination in CFA. CeA neurons positive for protein kinase C-δ, which are localized to the CeAl/c, express muscarinic receptors (**Extended Data Fig. 3**), and can rapidly suppress food intake^22,24^, likely make up a considerable portion of CFA-involved neurons. Interestingly, a subpopulation of CeAm neurons retained a preferential response to the CS in saline-conditioned control mice, neurons that may be activated via flavour-encoding insular cortex inputs^16^, involved in promotion of ongoing consumption^23,68^, or both.

While CFAs are known to form more readily with novel foods, we did not formally compare the formation of CFA in novel vs familiar flavours. One untested hypothesis is that diminished ACh as a CS becomes familiar decreases the likelihood of CFA formation, and that artificially driving ACh release in CeA during consumption of a familiar food would restore the ability for an animal to form a CFA. Such a result has already been observed with Cch infusion in insular cortex^41^.

Here, we provide evidence that neuromodulation facilitates associative plasticity over minutes-long delays in the CeA. This type of neuromodulatory priming may be commonplace in other forms of reward and aversive learning, and may be central to proposed models of retrospective causal learning, wherein a causal CS can be retroactively associated with an outcome due to the salience or novelty of the CS^76^.

## Methods

### Mice

This study was performed in strict accordance with the recommendations in the Guide for the Care and Use of Laboratory Animals of the National Institutes of Health. All animal procedures were pre-approved by Animal Care and Use Committees (IACUC) at the University of Washington (#4450-01). Transgenic mice on a C57BL/6J background were used for experiments. *Calca^Cr^*^e^^32^, *Calca^Flp^*::*Chat^Cre^*, *Chat^Cre^*, and Vgat-cre (*Slc32a1^Cre^*) mice were bred in the lab to obtain heterozygous offspring used for experiments. Mice used for behavioural experiments were singly housed to prevent damage to implants and avoid possible effects of socially observed malaise following LiCl injections. All other mice were group-housed. Mice were at least 8 weeks old prior to surgery and all groups included males and females of similar proportions. Mice were assigned to groups randomly and the experimenter was not blinded to group identity. Mice were kept on a reverse 12 hr light/dark cycle and behavioural experiments were conducted within the dark cycle.

### Head-fixed conditioned flavour avoidance (CFA) behaviour

Mice entered a water restriction protocol 3 days prior to head-fixed behaviour. On an initial head-fixed habituation day, mice were head-fixed and presented with free access to the water spout for 10 min. Beginning the next day, mice underwent 7 daily water-only habituation sessions, wherein mice were head-fixed and received 100 water trials. For all head-fixed behaviour sessions, spouts were presented for 3 s access periods per trial, with inter-trial intervals sampled randomly from an 8 to 13-s distribution. Following the final water-only habituation session, mice received a single saline injection (0.1 mL, 0.9%, i.p.) and returned to their home cage to habituate mice to injections.

On the conditioning day, in behaviour-only experiments, mice were presented with 100 trials of the conditioned stimulus (CS), sweetened Kool-Aid (0.06% unsweetened grape Kool-Aid powder in 5% sucrose^30^). In fibre photometry and *in vivo* calcium imaging experiments, mice were presented with 5 water trials followed by 95 CS trials, both to compare water and CS responses and ensure recording conditions were stable prior to the first CS trial. Following the head-fixed behaviour sessions, mice received either a saline injection or LiCl injection (180 mg/kg, i.p.). The exact timing of the LiCl injection varied by experiment, but in all cases occurred approximately 2-10 min following the final conditioning trial. Mice were returned to their home cage after injection, except during in vivo calcium imaging experiments, in which case mice were head-fixed for a 15-min post-injection recording before being returned to their home cage. On the following day, mice underwent a water-only recovery session. Conditioning and recovery sessions were repeated. The day after the final recovery session, mice underwent CFA Test, wherein they were presented with 50 trials each of water and CS, pseudo-randomly presented such that each block of 10 trials contained 5 water and 5 CS trials, for a total of 100 trials. For the CRISPR behaviour, photometry, and in vivo calcium imaging experiments, mice underwent 3 consecutive days of Tests. For the optogenetic terminal inhibition experiment, mice underwent 1 Test. In the initial behaviour-only experiment, following Test 1, mice were removed from water restriction for approximately 17 days before returning to water restriction, and undergoing an additional CFA Test 21 d after the initial Conditioning period.

### Fibre photometry

Mice were injected unilaterally with 500 nL of AAV9-hSyn-gACh4h (BrainVTA) in the right CeA (M-L, 2.90 mm; A-P, -1.50 mm; D-V, -4.50 mm). Optical fibres (400 µm core diameter, 0.48 NA, Doric) were implanted unilaterally to target the injection region (M-L, 2.90 mm; A-P, -1.50 mm; D-V, -4.50 mm). For ACh sensor recordings during PB*^Calca^*cell body and terminal stimulation *Calca^Cre^* mice were injected with 500 nL of AAV9-hSyn-gACh4h in the right CeA (M-L, 2.90 mm; A-P, -1.50 mm; D-V, - 4.50 mm) and 500 nL of AAVDJ/Syn-FLEX-Chrimson-tdTomato (Addgene; Lot #av169; titer 9.6e12) in the right PB (M-L, 1.40 mm; A-P, -4.80 mm; D-V, -3.50 mm). Optical fibres (400 µm core diameter, 0.50 NA, Inper) were implanted unilaterally to target the CeA (M-L, 2.90 mm; A-P, -1.50 mm; D-V, - 4.50 mm). An additional fibre (200 µm core diameter, 0.39 NA, RWD) was implanted to target the PB (M-L, 1.75 mm; A-P, -5.00 mm; D-V, -3.20 mm).

For simultaneous ACh sensor and PB*^Calca^* terminal recordings in CeA, *Calca^Cre^* mice were injected with 500 nL of AAV9-hSyn-rACh1h (BrainVTA; titer 2.6e12) in the right CeA (M-L, 2.90 mm; A-P, -1.50 mm; D-V, -4.30 mm) and 500 nL of AAVDJ/EF1a-DIO-GCaMP6f (Addgene; Lot #av6361; titer 3.9e12) bilaterally in the PB (M-L, 1.40 mm; A-P, -4.80 mm; D-V, -3.50 mm). Optical fibres (400 µm core diameter, 0.50 NA, Inper) were implanted unilaterally to target the CeA (M-L, 2.9 mm; A-P, - 1.50 mm; D-V, -4.50 mm).

To record neural activity with gACh4h expressed specifically in GABAergic neurons in CeA, mice were connected to a lock-in demodulated system (RZ10X; Tucker Davis Technologies (TDT)). The system was outfitted with 3 LEDs with the following configuration: 465 nm at 331 Hz / 70 mA for green sensors (gACh4; GCaMP8f); 560 nm at 523 Hz / 100 mA for red sensors (rACh1h), and 405 nm at 211 Hz / 8 mA. We used a 6-port minicube (FMC6_IE(400-410)_E1(460-490)_F1(500-540)_E2(555-570)_F2(580-680)_S, Doric Lenses) coupled to a 3 m patch cable (0.57 NA, Doric Lenses), which was coupled via ceramic split sleeve (2.5 mm diameter, Precision Fiber Products) to the optic fibre implanted in the mouse brain. Behavioural events were synchronized with fibre photometry signals using TTL inputs from the OHRBETS behaviour system^43^ to the TDT system.

Data were converted into a uniform data format and downsampled to 100 Hz. Photobleaching was corrected by first calculating F0, defined by fitting a double exponential decay to the raw signal, rank order filtered for the 5th percentile over a 1 min moving window. This F0 was subtracted from the raw signal, and the absolute value of the minimum value of the resultant signal was added to signal to ensure no values were negative, and the signal was then divided by F0 to calculate DF/F (F-F0/ F0). The z-score of the DF/F signal was calculated across head-fixed behaviour sessions for each mouse. The mean and standard deviation used for z-scoring were calculated using the concatenated DF/F signal across all head-fixed behaviour sessions for a given mouse.

For recordings performed following drug injection, the same procedure was used for calculation of DF/F, except that to best preserve slow fluctuations in signal related to drug effects, all fluorescence values from 30 s to 30 min were not used to fit the double exponential decay model used in the calculation of F0. Additionally, in the CRISPR confirmation experiments, data was not z-scored prior to comparison across animals.

### Fear conditioning and anaesthetized noxious stimuli photometry

Wild-type mice received injections of AAV9-hSyn-gACh4h in CeA and fibre implant as described above. Experiments were performed at least 4 weeks after surgery.

In the fear conditioning experiment, a patch cable was attached to the implanted fibre for photometry recording and the mouse was placed in a 24×24 cm chamber (MedAssociates, ENV-350CW) with a conductive metal floor. On day one, mice underwent 10 trials in which they received a 10 s tone (3 kHz, 80 db) that co-terminated with delivery of a foot shock administered through the metal floor (0.5 s, 0.3 mA). On day two, the mouse was returned to the same chamber, where a floor insert covered the metal grid. Mice received 30 trials of the same 10 s tone with no shock. For each day, the inter-trial interval between tones was selected on a variable schedule (mean, 90 s; range, 67 to 135 s). Timing of behavioural stimuli were recorded via TTL pulse delivered to the photometry acquisition system.

A separate cohort of mice was used for the noxious stimuli experiments. In these experiments, mice were anaesthetized with 1.5-2.0% isoflurane (mixed with 0.8 L per min oxygen)^28^. A patch cable was attached for fibre photometry measurements. In the tail pinch experiment, mice received a tail pinch via micro bulldog clamp (Fine Science Tools) for 5 s, with inter-stimulus intervals of at least 120 s and 5 trials total. In the thermal experiment, the tail was immersed in hot water for 10 s, with inter-trial intervals of at least 120 s.

### Characterization of *Calca* and *Chat* overlap in PB

To visualize overlap of *Calca*+ and *Chat*+ neurons in PB and their projections in CeA, *Calca^Flp^*::*Chat^Cre^*mice were injected bilaterally with a mix of AAV-DIO-eYFP and AAV-fDIO-mCherry in PB (M-L, 1.40 mm; A-P, -4.80 mm; D-V, -3.50 mm).

For the *in situ* hybridization experiment, wild-type mice were anaesthetized with sodium pentobarbital and phenytoin sodium (0.2 mL, i.p.), decapitated, and brains were quickly removed, frozen on crushed dry ice, and stored at -80 °C. 20 µm coronal sections were cut on a cryostat (Leica CM 1950), directly mounted onto glass slides (SuperFrost Plus), and stored at -80 °C. RNAscope fluorescence multiplex assay v2 was performed following the manufacturer’s guidelines. Probes used were: Chat (#408731) and Calca (#578771-C2).

Images were captured on a confocal microscope (Olympus FV-1200) at 20x magnification centreed on the external lateral PB. Probe images were subtracted from one another using the image calculator function in Fiji to remove background autofluorescence. Images were adjusted and regions of interest were drawn in Fiji before transfer to QuPath software for analysis. The subcellular detection function in QuPath was used to define transcript staining, and DAPI signals with 8 or more associated puncta were labeled positive.

### Analysis of Allen Brain Atlas Connectivity data

To identify and compare projection targets of PB*^Chat^* neurons, we analysed data from two experiments using Chat-IRES-Cre-neo mice with PB as a source structure (#265565965 and #183901489) in the Allen Brain Atlas Connectivity map. AAV injection structures for either experiment were almost exclusively PB (89% PB, 2% LC, 7% fibre tracts in #183901489; 95% PB, 5% fibre tracts in #265565965). For projection density by target region comparisons (**Fig. 2l**), pre-calculated projection density values by region and CeA subregion were taken directly from the downloadable .xml datasets.

For analysis of projection density by subregion and anterior-posterior position (**Fig. 2m**), we used volumetric connectivity data using the AllenSDK MouseConnectivityCache (Allen Mouse Brain Common Coordinate Framework, 25 µm isotropic resolution). For a given experiment, we obtained the full 3-dimensional projection density volume, and, for each CeA subregion, a binary structure mask derived from the CCFv3 annotation volume. Each mask was restricted to the hemisphere ipsilateral to the injection site. For each coronal slice, projection density was defined as the mean voxel-wise projection density across all structure-mask voxels at that slice, including voxels with no projection signal. Structure volume at each slice was computed as the voxel count multiplied by the isotropic voxel volume (25 µm), converted to mm. A-P position is reported as an approximate bregma-relative coordinate, obtained by linear offset (5.4 mm) following an established CCF-to-Paxinos correspondence implemented in the cortex-lab/allenCCF toolbox.

### CRISPR-Cas9 viral knockdown of *Chat* and *Slc18a3*

For knockdown of *Chat* and *Slc18a3* in PB, *Chat^Cre^* mice were bilaterally injected with 850 nL of a 1:1 mix of AAV1-CMV-FLEX-SaXCas9-U6-sgChat and AAV1-CMV-FLEX-SaCas9-U6-sgSlc18a3 (425 nL of each virus per injection site; UW NAPE Center MGRC) bilaterally in PB (M-L,1.40 mm; A-P, -4.80 mm; D-V, -3.50 mm). Control mice were bilaterally injected with 500 nL of AAV1-CMV-FLEX-SaCas9-U6-sgRosa26 (UW NAPE Center MGRC; titer 3.1e10).

For the CRISPR photometry validation experiment, in addition to a unilateral injection in either left or right PB using the viral strategy described above, *Chat^Cre^*mice were also injected with 400 nL of AAV1-hSyn-gACh4m (UW NAPE Center MGRC) bilaterally in CeA (M-L, 2.90 mm; A-P, -1.50 mm; D-V, -4.30 mm). CRISPR virus injections were counterbalanced to account for the possibility of different ACh signalling profiles in right vs left CeA. Optical fibres (400 µm core diameter, 0.48 NA, Doric) were implanted bilaterally to target the CeA (M-L, 2.90 mm; A-P, -1.50 mm; D-V, -4.50 mm). 6 weeks after surgery, mice were placed in an empty cage and photometry patch cables were attached to the optical fibre cannulae. After a period of 15 mins, mice were given an injection of lithium chloride (180 mg/kg, i.p) or saline (0.9%, 0.1 mL, i.p.) and recorded for 45 mins. Traces were analysed as described in **Fibre photometry** above.

For the electrophysiology experiment, *Chat^Cre^* mice were bilaterally injected with 1000 nL of a 0.4:0.4:0.2 mix of AAV1-CMV-FLEX-SaXCas9-U6-sgChat (400 nL/injection), AAV1-CMV-FLEX-SaCas9-U6-sgSlc18a3 (400 nL/injection), and AAV1-DIO-ChR2-eYFP (200 nL/injection; UW NAPE Center MGRC) in PB (M-L,1.40 mm; A-P, -4.80 mm; D-V, -3.50 mm). Control mice were injected with a 0.5:0.2 mix of AAV1-CMV-FLEX-SaCas9-U6-sgRosa26 (500 nL/injection) and AAV1-DIO-ChR2-eYFP (200 nL/injection). All experiments were performed 6 wk after viral injection and as described in ***Ex vivo* slice electrophysiology** below.

### Immunohistochemistry

For histology experiments, mice were anaesthetized with sodium pentobarbital and phenytoin sodium (0.2 mL, i.p.) and intracardially perfused with ice-cold PBS followed by 4% PFA. After a 24 h post-fixation in PFA, brains were stored in 30% sucrose for at least 3 days, followed by placement in a mold with OCT, and frozen at -80°C. 35 μm coronal sections were collected for CeA and PB on a cryostat (Leica) and stored in sucrose-based cryoprotectant for long term storage at -20°C

For projection tracing, sections were washed two times in PBS then incubated in a blocking solution consisting of 3% normal donkey serum and 0.2% Triton-X in PBS for 1 h at room temperature. Sections were then moved to blocking solution containing primary antibodies chicken-anti-GFP (1:10000, Abcam, ab13970), and rabbit-anti-DsRed (1:1000, Takara, 632496) and incubated overnight at 4°C. The following day, sections were rinsed 3 times in PBS and then incubated for 1 h in PBS with secondary antibodies Alexa Fluor 488 donkey anti-chicken (1: 500, Jackson ImmunoResearch, AB 2340375) and Alexa Fluor 594 donkey anti-rabbit (1:500, Jackson ImmunoResearch, AB 2340621). After 3 more PBS rinses, sections were mounted on glass slides and coverslipped with Fluoromount-G with DAPI (Southern Biotech).

For ChAT staining in validation of CRISPR constructs, sections were washed two times in PBS then incubated in a blocking solution consisting of 3% normal donkey serum and 0.25% Triton-X in PBS for 1 h at room temperature. Following the block, sections were incubated in sodium citrate buffer for 30 min at 60°C for antigen retrieval. They were then moved to a blocking solution containing the primary antibody goat anti-ChAT (1:500, Millipore, Ab144p) and incubated overnight at RT. The following day, sections were rinsed 3 times in PBS and then incubated for 2 h in blocking solution with the secondary antibody donkey anti-goat Cy5 (1:500, Jackson ImmunoResearch, AB 2340415). After 3 more PBS rinses, sections were mounted on glass slides and coverslipped with Fluoromount-G with DAPI (Southern Biotech).

### Inhibition of PB*^Chat^* terminals in CeA during CFA conditioning

To inhibit PB*^Chat^* terminals in CeA during CS exposure in CFA, *Chat^Cre^* mice were bilaterally injected with AAV1-SIO-eOPN3-mScarlet-WPRE (Addgene; titer 7.0e12) in PB (M-L,1.40 mm; A-P, -4.80 mm; D-V, -3.50 mm). Control mice were bilaterally injected with AAV5-EF1a-DIO-mCherry (Addgene; titer 4e11). Optical fibres (300 µm core diameter, 0.39 NA, RWD) were implanted bilaterally to target the CeA (M-L, 2.90 mm; A-P, -1.50 mm; D-V, -4.50 mm). Experiments were performed approximately 8 weeks after surgery.

Mice underwent the head-fixed CFA behaviour experiment described above, with some modifications. On the final water-only habituation day, patch cables were attached to implanted ferrules to habituate mice to this process. The laser remained off for this session. Then, prior to both conditioning sessions with CS exposure, mice were head-fixed and patch cables were attached to the ferrules. The laser was turned on approximately 1 min before the first spout extension to ensure sufficient terminal inhibition during the initial trials. Photoinhibition continued for the duration of the ∼22 min conditioning session (532 nm laser, 10 mW power, 500 ms laser pulses at 0.4 Hz). The laser was turned off once the behaviour session, and mice were immediately removed from the head-fixed behaviour apparatus and returned to their home cage. After a delay of approximately 5 mins, mice received a LiCl injection (180 mg/kg) and were returned to their home cage. Patch cables were not attached during recovery or the Test session.

### *Ex vivo* 2-photon imaging

For measurement of ACh release following optogenetic stimulation of PB terminals, *Chat^Cre^* or *Calca^Cre^* mice were injected bilaterally with 500 nL of AAV8-Ef1a-DIO-ChRmine-mScarlet-Kv2.1-WPRE (UW NAPE Center MGRC; Lot #7082; titer 6.5e12) in PB (M-L,1.40 mm; A-P, -4.80 mm; D-V, -3.50 mm) and bilaterally with 500 nL of AAV9-hSyn-gACh4h (BrainVTA) in CeA (M-L, 2.90 mm; A-P, -1.50 mm; D-V, -4.50 mm). For *ex vivo* 2-photon calcium imaging experiments, Vgat-cre mice were injected bilaterally with 500 nL of AAV1-pGP-Syn-FLEX-jGCaMP8f in CeA (M-L, 2.90 mm; A-P, -1.50 mm; D-V, -4.45 mm). For PKA sensor experiments, Vgat-cre mice were injected bilaterally with 500 nL of AAV1-ExRai-AKAR2 (Janelia; titer 1.89e13) in CeA (M-L, 2.90 mm; A-P, -1.50 mm; D-V, -4.45 mm).

All *ex vivo* slice experiments were performed on brain slices collected at approximately the same time of day. Mice were anaesthetized with an i.p. Injection of Euthasol (0.06 mL/30 g). Mice were perfused with 10 mL of ice-cold NMDG ACSF (92 mM NMDG, 2.5 mM KCl, 1.25 mM NaH_2_PO_4_, 30 mM NaHCO3, 20 mM HEPES, 25 mM glucose, 2 mM thiourea, 5 mM Na-ascorbate, 3 mM Na-pyruvate, 0.6 mM CaCl_2_, 10 mM MgSO_4_·7H_2_O, and 12 mM N-Acetyl-L-cysteine; pH adjusted to 7.3-7.4). Mice were then decapitated. The brain was extracted and placed in NMDG ACSF solution for 2 min. Afterwards, coronal slices (300 µm) were sectioned using a vibratome (VT1200s, Leica, Germany) in a bath of cold NMDG ACSF solution, and then were incubated in NMDG ACSF at 34°C for approximately 14 min. Slices were then transferred into a holding solution of HEPES ACSF (92 mM NaCl, 2.5 mM KCl, 1.25 mM NaH_2_PO_4_, 30 mM NaHCO_3_, 20 mM HEPES, 25 mM glucose, 2 mM thiourea, 5 mM Na-ascorbate, 3 mM Na-pyruvate, 2.4 mM CaCl_2_, 2 mM MgSO_4_·7H_2_O and 12 mM N-Acetyl-l-cysteine, bubbled at room temperature with 95% O_2_/ 5% CO_2_) for at least 45 mins until recordings were performed.

The recording solution (120 mM NaCl, 3.5 mM KCl, 1.25 mM NaH_2_PO_4_, 26 mM NaHCO_3_, 1.3 mM MgCl_2_, 2.4 mM CaCl_2_ and 11 mM D-(+)-glucose, continuously bubbled with 95% O_2_/ 5%CO_2_) was delivered to slices via superfusion driven by a peristaltic pump (flow rate of 4-5 mL/min) and was held at 30°C. An Olympus 4x UPlanFLN lens was used to localize sensor expression. For recordings, an Olympus 25x XLPLN25XWMP lens (for GAC4h-based terminal stimulation experiments) or Olympus 10x (for GCaMP-based pharmacology experiments) was used. Images were acquired with Olympus FV31S-SW software interfaced with an FVMPE-RS Olympus 2p scope powered by a Mai Tai Ti:sapphire laser with dispersion compensation running at 920 nm. Individual frames were acquired at either 7.5 or 15 Hz with a Galvano resonant scanner with a resolution of 512×512 pixels.

For optogenetic stimulation of PB terminals while recording ACh biofluorescence in CeA, we used a 615 nm LED (CoolLED) controlled by a Master-9 stimulator, programmed for 20 Hz stimulation for 4.5 s with 5 ms pulse durations. LED power was measured at 3 mW. Stimulation was first performed in ACSF only. Then, TTX and 4AP were bath-applied for at least 2 min before another stimulation recording was performed.

For GCaMP-based carbachol wash-in experiments, all recordings were performed with picrotoxin (PTX; 100 μM) in the recording solution. For carbachol (Cch) wash-in experiments, the perfused solution was switched from ACSF to ACSF containing Cch (20 uM) 2 mins after the start of a recording. Travel time to the recording chamber was estimated to be approximately 2 min, with solution in the recording chamber reaching a target steady-state drug concentration an estimated 60-90 s later. In antagonist experiments, slices were pre-treated with ACSF containing antagonists for at least 5 mins before beginning recording. During recordings, ACSF containing Cch and antagonists was perfused beginning 2 min after recording start as described. Antagonists used were: a combination of NBQX (20 µM), D-AP5 (50 µM), and mecamylamine (10 µM); scopolamine (10 µM); telenzepine (0.1 µM); 4-DAMP (1 µM); and VU 6019650 (1 µM). In pharmacology experiments, slices were not reused for additional recordings after Cch wash.

For acetylcholine puffing experiments, all recordings were performed in the presence of PTX (100 µM) and a low concentration of cholinesterase inhibitor physostigmine (1 µM). Glass pipettes with identical dimensions as those used for whole-cell electrophysiology experiments were filled with recording ACSF containing acetylcholine hydrochloride (100 mM), as well as Alexa Fluor 594 hydrazide (13.2 µM) for visualization of the pipette during placement. The pipette was placed above a section of tissue in the imaging field of view. During recording, slight positive pressure (approximately 50 kPa) was briefly applied (500 ms puff duration), and time-locked fluorescence responses were recorded. Recordings were only performed once on a given slice.

For AMPA puffing experiments, all recordings were performed in the presence of PTX (100 µM). Glass pipettes were filled with recording ACSF containing (*RS*)-AMPA hydrobromide (1 mM), as well as Alexa Fluor 594 hydrazide (13.2 µM) for visualization of the pipette during placement. Puffs were applied (50 kPa, 50 ms puff duration) and time-locked fluorescence responses were recorded. Puff responses were recorded before, during, and 15 min after a brief wash-in period, where perfusate was ACSF (control) or ACSF with carbachol (Cch, 20 µM). Experiments were only performed once on a given slice.

For PKA sensor experiments, all recordings were performed in the presence of tetrodotoxin (TTX; 1 µM) to eliminate the possibility of calcium transient-related activation of PKA or polysynaptic signalling, and PTX (100 µM). Scopolamine (10 µM) was used to antagonize muscarinic receptors. In pharmacology experiments, slices were not reused for additional recordings after Cch wash.

### Analysis of *ex vivo* fluorescence traces

To extract calcium fluorescence traces, Suite2p^77^ was used for functional ROI detection. ROIs were manually inspected to remove neurons visibly outside of CeA. In all *ex vivo* experiments, raw fluorescence traces were neuropil-subtracted with a scaling factor of 0.7.

For the Cch wash-in experiment, neuropil-subtracted fluorescence traces were z-scored using fluorescence during a 200 s baseline period before Cch wash began. Z-scored traces were then baseline-subtracted using mean fluorescence over the same baseline window. For each neuron, we calculated the baseline-subtracted mean fluorescence during the Cch wash period (defined as a 240 s window beginning at the 360 s mark of the recording, when Cch concentration should be at or near peak values).

For the ACh puff experiment, neuropil-subtracted fluorescence traces were z-scored using fluorescence during a 70 s baseline period beginning 20 s after recording start and ending 10 s before puff onset. For each neuron, we calculated the puff response as the mean baseline-subtracted fluorescence value over a 30 s window immediately following puff onset.

For the AMPA puff experiment, functional ROI detection was performed in Suite2p across the concatenated recordings from the Baseline, Wash, and Post recordings. Neuropil subtraction was performed. Within a given recording, traces were baseline-subtracted using a 25 s baseline window beginning at the start of the trace and ending 5 s before AMPA puff. For each neuron, z-scoring across the concatenated, baseline-subtracted trace was performed using all fluorescence values from the Baseline recording so that subsequent puff responses could be compared in scale to the initial puff response. Then, for each neuron, for each AMPA puff, we calculated the puff response as the mean baseline-subtracted fluorescence value over a 30 s window immediately following puff onset.

For the PKA sensor experiment, CellPose^78^ was used within Suite2p for anatomical ROI detection, since PKA signals tended to be visibly less dynamic than those observed during calcium imaging experiments. Fluorescence traces were z-scored and baseline-subtracted the same as in the calcium Cch wash-in experiment. For each neuron, we calculated the baseline-subtracted mean fluorescence during the Cch wash period (defined as a 300 s window beginning at the 360 s mark of the recording).

### *Ex vivo* slice electrophysiology

Brain slice collection was performed with identical methods and solutions as in the *ex vivo* slice 2-photon calcium imaging experiments. Following incubation in HEPES ACSF, whole-cell recordings were performed using a Multiclamp 700B (Molecular Devices, Sunnyvale, CA) using pipettes with a resistance of 4-6 MOhm. During recordings, slices were perfused with a recording ACSF solution (same as in ***Ex vivo* 2-photon imaging** experiment, continuously bubbled with 95% O2/5% CO2). Infrared differential interference contrast–enhanced visual guidance was used to select neurons that were 3–4 cell layers below the surface of the slices. The recording solution was delivered to slices via superfusion driven by a peristaltic pump (flow rate of 4-5 mL/min) and was held at 30°C.

In the PB ChR2 experiment (**Fig. 4d-f**), PB projections were visualized by the presence of YFP using a fluorescence microscope (Scientifica SliceScope Pro 1000; LED: SPECTRA X light engine (Lumencore)). Neurons were voltage clamped at −70 mV, and the pipette series resistance was monitored throughout the experiments by hyperpolarizing steps of -10 mV with each sweep. If the series resistance changed by >20 % during the recording, the data were discarded. Whole-cell currents were filtered at 1 kHz and digitized and stored at 10 KHz (Clampex 10; MDS Analytical Technologies). For paired pulse ratio (PPR) measurements, a pair of blue light pulses (3 ms width; 50 ms inter-stimulus interval) was delivered every 30 seconds for 7-10 sweeps. For each neuron, the mean peak optically evoked excitatory postsynaptic current (oEPSC) was taken for each stimulus, and used to calculate the neuron’s mean oEPSC amplitude and its PPR. Two-sample t-tests were performed to test for group differences at each pulse oEPSC amplitude and for PPR.

In the M-current experiments (**Extended Data Fig. 4c-i**), cells were sampled throughout the CeA. Neurons were voltage clamped at -30 mV and given 1-s, 10 mV hyperpolarization steps (voltage clamp at -40 mV) every 30 s. After 5 sweeps, carbachol (20 µM) was added to the recording ACSF and another 5 sweeps of hyperpolarizing steps were performed. The mean trace was calculated for each cell for baseline and carbachol periods. We then used 1-sample t-tests to test for the effects of carbachol on holding current (I_Hold_); deactivation current (I_deactivation_, or the difference in mean holding current in 50 ms window immediately after voltage step onset (I(V_H_)) and steady state holding current (I_SS_), calculated over the final 100 ms of the voltage step), reactivation current (Ireactivation, or the difference in mean holding current in a 50 ms window immediately after voltage step offset (I_ret_) and in the final 100 ms window of the first 1 s following voltage step offset (I_SS_,_ret_)); and time constants τ_deactivation_ and τ_reactivation_. τ values were calculated by fitting a single exponential function to the 1 s windows during and immediately after the voltage step. If there was no detectable sag current (I_deactivation_ or I_reactivation_ < |2 pA|), fitting was skipped and τ was set to 0 ms.

For current clamp recordings (**Extended Data Fig. 4a-b**), pipettes were filled with a potassium-based internal solution containing 120 mM potassium gluconate, 0.2 mM EGTA, 10 mM HEPES, 5 mM NaCl, 1 mM MgCl_2_, 2 mM Mg-ATP and 0.3 mM NA-GTP, with the pH adjusted to 7.2 with KOH. The resting membrane potential was normalised to -70 mV. In some cases, the resting membrane potential was brought to -70 mV by small current injections of <50 pA. If the compensating current was > 50 pA, the recording of the neuron was terminated. For acetylcholine puffing experiments (**Extended Data Fig. 4b**), recordings were performed in the presence of a low concentration of cholinesterase inhibitor physostigmine (1 µM). Glass pipettes with identical dimensions as those used for whole-cell electrophysiology experiments were filled with recording ACSF containing acetylcholine hydrochloride (100 mM). The pipette was placed above a section of tissue in the imaging field of view. During recording, slight positive pressure (approximately 50 kPa) was briefly applied (500 ms puff duration).

All recordings were performed with picrotoxin (100 µM) present in the recording ACSF solution. All slice experiments were completed within 4 hours after slices were made to maximize cell viability and consistency.

### Single-nucleus RNA sequencing

A total of 16 mice were used for the single-nucleus RNA sequencing experiment. Mice were water restricted and injected with either saline or LiCl (180 mg/kg). Approximately 50 mins after injection, mice were perfused with formaldehyde (37%) and brains were extracted. Fixed brains were immediately sectioned on a Vibratome VT1200S, and tissue punches containing CeA were collected. Tissue punches were fixed and dissociated following 10X’s protocol for chromium fixed RNA profiling (CG000553). Probe hybridization; GEM generation and barcoding; GEM recovery and pre-amplification; and gene expression library construction were all completed following 10X’s protocol for chromium fixed RNA profiling for single-plexed samples (CG000691). Dual Index Plate TS Set A (PN: 3000511; LN: 402663365) was used for library construction. Sequencing was performed by Novogene (NovaSeq X Plus Series (PE150)).

All downstream snRNAseq analysis was completed using the R package Seurat, version 4.2. Samples were filtered to exclude nuclei with fewer than 800 or greater than 40,000 reads, or with greater than 3% of reads from mitochondrial genes (which is often associated with low quality in snRNAseq.) The SCTransform algorithm was utilized to normalize each sample prior to and after integration, which we have found to produce superior results when compared to log-normalization strategies. This method uses a GLM-based approach in which sequencing depth for each nucleus is a covariate, improving our ability to make comparisons between samples that were run separately and likely contain technical variation. We utilized the percentage of mitochondrial reads as an additional regression variable to further control for discrepancies in cell quality between samples. Normalization is followed by PCA, which identifies PCs from the gene expression data for each sample, 20 of which were used to generate a UMAP. After nearest-neighbor analysis is completed, each cell is classified into a putative cluster. Clustering resolution can be adjusted upwards or downwards to produce more or fewer clusters respectively, but we observed satisfactory results with resolution = 0.3 for most samples. Doublets were identified and removed using the R package DoubletFinder, after which samples were re-normalised. Samples were integrated together using Seurat’s SCT-based integration strategy.

### *In vivo* 2-photon calcium imaging during behaviour

To perform large scale in vivo calcium imaging in CeA, in the Base experiment, Vgat-cre mice were injected with 500 nL/site of AAV1-Syn-FLEX-GCaMP6s.WPRE.SV40 (Addgene; titer 2.1e13) in right CeA at two sites along the same dorsal-ventral path (M-L, 2.90 mm; A-P, -1.40 mm; D-V, -4.80 and - 4.20 mm). In the CRISPR experiment, *Chat^Cre^* mice were injected with 500 nL/site of AAVDJ-hSyn-GCaMP6s (Stanford Vector Core; titer 1.5e13) in right CeA at the same coordinates. In this experiment, mice were also bilaterally injected with 850 nL of a 1:1 mix of AAV1-CMV-FLEX-SaXCas9-U6-sgChat and AAV1-CMV-FLEX-SaCas9-U6-sgSlc18a3 (UW NAPE Center MGRC) bilaterally in PB (M-L,1.40 mm; A-P, -4.80 mm; D-V, -3.50 mm). Control mice were bilaterally injected with 500 nL of AAV1-CMV-FLEX-SaCas9-U6-sgRosa26 (UW NAPE Center MGRC; titer 3.1e10) in PB.

In both experiments, mice were implanted with an 8 mm-long prism lens with a 1.0 mm^2^ imaging face (manufacturer, OptoSigma). Briefly, a square-shaped section of skull of an area of approximately 1.4 mm^2^ over and anterior to the site of viral injection was removed to allow for implantation of the lens anterior to the injection site. Care was taken to place the lens in a holder such that the imaging face would be parallel to the coronal plane of the brain. The lens was lowered at a speed of approximately 2 mm/min until reaching a depth of -4.00 mm D-V. From here, the lens was lowered at a speed of 0.2 mm/min. The final targeted position of the rightmost corner of the bottom of the lens was (M-L, 3.30 mm; A-P, -1.40 mm; D-V, -5.00 mm), such that the centre of the imaging face would be positioned immediately adjacent to the injection site (M-L, 2.80 mm; A-P, -1.40 mm; D-V, - 4.50 mm). The skull was covered with Metabond, and superglue was used to secure the lens prior to removal of the lens holder. A custom headbar was positioned above the skull around the lens, and dental cement was used to secure the headbar and complete the implant.

All *in vivo* imaging was performed on a Bruker Investigator two-photon microscope through a Cousa objective (20 mm WD)^79^. Functional imaging was performed using an Insight X3 laser at 920 nm. Data were acquired at 15 Hz on resonant galvo mode (2-frame averaging). Images were acquired at 512×512 pixel resolution. An OHRBETS system was used during imaging experiments. The OHRBETS stage was mounted on a Thorlabs goniometer platform (TTR001/M) to facilitate fine adjustments of the imaging plane. To register a consistent imaging plane across days, we collected images of the imaging plane using 32-frame imaging. These images were used to adjust the imaging position and depth on the following session to ensure anatomical landmarks and neuronal positions were consistent across sessions.

Following recording, we tracked imaged neurons across sessions by concatenating all recordings and performing functional ROI detection with Suite2p. This provided motion detection within days and accounted for small misalignments between sessions. Neurons were verified as tracked if they had transients across multiple days.

To extract fluorescence traces for analysis, we started with the raw, concatenated fluorescence trace for a given neuron. First, neuropil subtraction was performed with the neuropil trace calculated by Suite2p, scaled by 0.1. The resulting neuropil-subtracted trace was low-pass filtered using a zero-phase second-order Butterworth filter with a cutoff frequency of 2 Hz (implemented via Matlab’s filtfilt function). To calculate DF/F, the 8th percentile of fluorescence values across the concatenated trace was calculated; this value was used as F0 in a standard DF/F calculation. Then, within each session, traces were passed through a 1-s Gaussian filter, and the 8th percentile of fluorescence values within the session was subtracted from the within-session trace to normalize baseline values across sessions. We concatenated traces from head-fixed behaviour sessions and used the mean and standard deviation of the concatenated trace for z-scoring all traces. Z-scoring statistics were derived from behaviour sessions only to avoid possible confounds from group differences during injection sessions.

Behavioural data was synchronized with imaging data using voltage signals sent from the OHRBETS system to the Bruker system during acquisition, where voltage steps marked each spout extension and retraction. Licks were acquired by the OHRBETS system, and their timing in relation to the spout extension onset time was used to relate lick timing to imaging data.

### Analysis of *in vivo* calcium traces

For analysis of calcium signals during behavioural sessions, a peristimulus histogram was calculated for each trial using the 2 s before and 8 s following spout extension. Traces were baseline corrected using the last 1 s of the period before spout extension.

To visualize neuronal metrics in space, we used estimated bregma space values determined during atlas registration (see **Atlas registration of 2-photon *in vivo* imaged neurons**). Values in the anterior-posterior axis were collapsed to create a 2-D visualization in bregma space. For visualization purposes only, CeA subregion boundaries were hand-drawn based on the apparent boundaries when plotting by registration-determined subregion (**Extended Data Fig. 7**). To produce heatmaps, neurons were binned into a fixed 2-D grid in coronal space with a bin width of 0.025 mm. Within each bin, the metric of interest was summarized as the either the median (for changes in peak statistics) or mean across neurons, computed separately per group. Each group’s binned grid was then spatially smoothed by convolution with an isotropic, exponentially decaying, distance-dependent kernel. The kernel was normalized once, globally, to a sum of 1. Finally, each group’s smoothed grid was masked by that group’s neuron-occupancy pattern, such that unoccupied bins were removed if >=5 of its Moore-neighborhood neighbors were also unoccupied, limiting plots from expanding into unsampled areas for a given group.

To assess neuronal preference at Conditioning 1 (**Fig. 5f-g**), a linear mixed effects (LME) model was used. Only trials with >=1 lick were used, and analysis was restricted to the first 40 trials of Habituation and Conditioning 1 sessions. Mean fluorescence over a 1-5 s window following spout extension was predicted by the following fixed effects: flavour (water or CS); total lick count for the trial; and session number. Solution identity and session number were treated as categorical variables. After running the LME model, the Benjamini-Hochberg procedure was used to adjust p-values based on the number of effects terms in the model, with a threshold of 0.05. For neurons with significant flavour encoding, valence of the beta weight for flavour was used to label neurons as water- or CS-preferring.

For analysis of post-injection traces, all traces were baseline subtracted using the 8^th^ percentile of fluorescence values within the first 30 s of the recording. We plotted these baseline-subtracted fluorescence traces separately for CS-preferring neurons, as assessed with the LME approach, vs all other neurons. To quantify preferential reactivation of CS-preferring neurons in post injection traces (**Fig. 5i-n**), we calculated the mean post-injection trace for each neuron across the two injection sessions. This value was used in a linear mixed effect model, where mean injection fluorescence was predicted by fixed effects for group (ie, Saline vs LiCl), CS preference (0 or 1), and their interaction, with a random effect for neuron ID. In this model, group, CS preference, and neuron ID were all treated as categorical variables. Estimated marginal means were used for post-hoc comparisons to assess group differences in activation based on CS preference during Conditioning.

For peak amplitude and timing analysis, for a given neuron, we determined its peak fluorescence value over an 8 s window post spout access onset over a defined trial range on Test 1 (trials 1-10 for **Fig. 6**). The baseline-subtracted amplitude of this peak value and its time relative to spout onset access were recorded. These values were averaged across trials and Test sessions to calculate mean peak time and amplitudes for CS trials. The slope-to-peak was calculated by dividing the peak amplitude by the time-to-peak. For each peak metric, values were used in an LME model, where peak statistic was predicted by fixed effects for group, solution, and their interaction, with a nested random effect for neuron ID within mouse ID. Estimated marginal means were used for post-hoc comparisons to assess group differences by solution.

To measure the change in peak statistics from Conditioning 1 to Test 1 (**Fig. 6e,f,i,j**), peak statistics were calculated as described above. For each neuron and solution, the difference in peak metric between the two sessions was calculated. These difference metrics were used in a linear fixed effects model, where the change in peak metric (peak time or slope) was predicted by fixed effects for group, solution, and their interaction, with a nested random effect for neuron ID within mouse ID. Estimated marginal means were used for post-hoc comparisons to assess group differences by solution.

To predict last-lick time using neural data, we examined trials 1-40 over Tests 1-3. First, to quantify the relationship between slope-to-peak values and CS last-lick times (**Fig. 6k**), population PSTH slope-to-peak values were calculated for each trial and z-scored within individual subjects. Then, using an LME model, last-lick times were predicted by fixed effects for population slope-to-peak value, group, solution, and their interactions, with a random effect for mouse ID. Estimated marginal means of linear trends of the slope-to-peak value on last-lick time for CS were performed to determine whether slope-to-peak values significantly predicted last-lick times, and whether the relationship varied by group.

To predict last-lick time using neural data (**Fig. 6l**), we first summarized population activity using a multi-bin temporal basis, where the 6 s period post access onset was divided into 30 non-overlapping, ∼0.13 s bins. For every neuron and every bin, we calculated the mean baseline-subtracted fluorescence. We concatenated values across bins, yielding a 3-dimensional feature matrix of trials x neurons x 30 bins. We used supervised dimensionality reduction to find a single axis (LLT axis) whose projection covaried maximally with actual, trial-by-trial last lack times. This LLT axis was defined as the coefficient vector of a partial least squares (PLS) regression of true last-lick time on the z-scored temporal-basis features, using 4 latent components (a value fixed across all mice and chosen via nested, cross-validated search that maximized held-out predicted performance), normalised to unit length. LLT axis scores used for evaluation were strictly held out. We used 5-fold cross validation, where for each training fold, 80% of trials were used to estimate the PLS axis; the training-fold scaler was then applied to the remaining, held-out trials, which were projected onto the training-fold axis. This approach gives one held-out LLT axis score per trial. Performance was quantified per mouse as the squared Pearson correlation (R^2^) between the held-out LLT axis scores and the actual last-lick times. Significance was assessed with a shuffle null distribution approach. Actual last-lick time labels were randomly permuted across a given mouse’s trials and the cross-validation procedure was re-run on the permuted labels 100 times, forming a null distribution of held-out R^2^ values. The per-mouse permutation p-value was the fraction of shuffled R^2^ values that equaled or exceeded the non-shuffled R^2^ value.

To determine how CS preference varied by neuron based on CeA subregion (**Fig. 6-p**), for each neuron, we calculated the mean PSTH during water and CS trials over the first 10 trials of Tests 1-3. We calculated the mean CS and water responses using the first 1.5 s of access, reasoning that licking differences across the full 3 s access period render neural data less comparable. Response differences were calculated by subtracting the mean water response from the mean CS response. Using an LME model, response differences were predicted using group, CeA subregion (CeAc, CeAl, or CeAm), and their interaction as fixed effects, with a random effect for mouse ID. Pairwise contrasts between groups by subregion were performed with estimated marginal means.

### Atlas registration of *in vivo* 2-photon imaged neurons

For atlas registration of *in vivo* 2-photon imaged neurons (**Extended Data Fig. 7**), upon completion of imaging experiments, mice were anaesthetized with sodium pentobarbital and phenytoin sodium (0.2 mL, i.p.) and intracardially perfused with ice-cold PBS followed by 4% PFA. Mice were decapitated, and heads were post-fixed in PFA. After a post-fixation of 24-48 h, heads were rinsed with PBS. Because histology with prism lenses sufficient for post-hoc registration requires tissue to be collected parallel to the imaging face, care was taken to block the brain prior to slice collection. Bone and non-brain tissue below the headcap were removed, leaving the brain, the dorsal surface of the skull, and the headcap intact. The headcap was then rested atop a square mold with the brain suspended beneath. The mold was filled with a warmed 8% agarose solution and cooled. This concentration of agarose permitted removal of the headcap and implant while leaving the brain suspended in agarose. Coronal sections (200 μm) containing the imaged region in CeA were collected. Sections that contained a visible imprint of the prism face, or the first intact coronal sections when cutting anterior-to-posterior, were used for registration. Sections were mounted on glass slides and coverslipped with Fluoromount-G with DAPI (Southern Biotech).

Stitched images of the entire section were collected on a confocal microscope and used as an input in the registration pipeline, wherein cells imaged in vivo were mapped to Allen Mouse Brain Common Coordinate Framework (CCFv3) voxel coordinates and to bregma-relative stereotaxic coordinates using a custom open-source napari plugin (FOV-atlas-registration; https://github.com/kenjp1223/FOV-atlas-registration). Registration proceeded in three stages: *in vivo* field-of-view (FOV)-to-section alignment, confocal section-to-atlas alignment, and coordinate readout. The FOV was the mean *in vivo* fluorescence image extracted from the Suite2p output used for analysis, where the median pixel coordinate of each ROI was used as the cell centroid for all subsequent transformations.

First, the FOV image and the corresponding histological section were displayed side by side, and ∼10-30 corresponding anatomical landmark pairs were placed manually. A two-dimensional similarity transformation (rotation, isotropic scaling, and translation) was estimated from these pairs by least-squares orthogonal Procrustes analysis and applied to all cell centroids.

Because histological sections are rarely cut exactly in the atlas plane, an oblique virtual section was next extracted from the Allen CCFv3 template volume. Three Euler rotation angles and a slice index were adjusted until the resliced atlas plane visually matched the histological section. Residual local differences arising from tissue deformation and histological processing were then corrected by non-linear registration. Approximately 30 anatomical landmark pairs were placed between the resliced atlas plane and the section, targeting the outer brain contour, ventricles, and fibre tracts. A thin-plate spline (TPS) transformation was fitted from these pairs. Cell coordinates were mapped through this transformation into the resliced atlas plane and then returned to native CCFv3 voxel space by applying the inverse of the reslicing rotation.

Each cell’s CCFv3 voxel index was then used to query the structure ID to obtain region name, acronym, and parent structure. Voxel indices were additionally converted to Paxinos-Franklin bregma-relative stereotaxic coordinates following published CCF-to-Paxinos calibrations (Zhao, https://bohanzhao.com/atlas/; Cortex Lab allenCCF, https://github.com/cortex-lab/allenCCF). Coordinates are reported in millimetres relative to bregma. For each session, the plugin exports all landmark pairs, the fitted transformation parameters, and a per-cell table containing FOV, section, and CCFv3 coordinates together with assigned region identity and stereotaxic coordinates, allowing every registration to be reproduced from the saved parameters. Analysis was performed in Python, using napari, NumPy, SciPy, and scikit-image.

## Key resources table

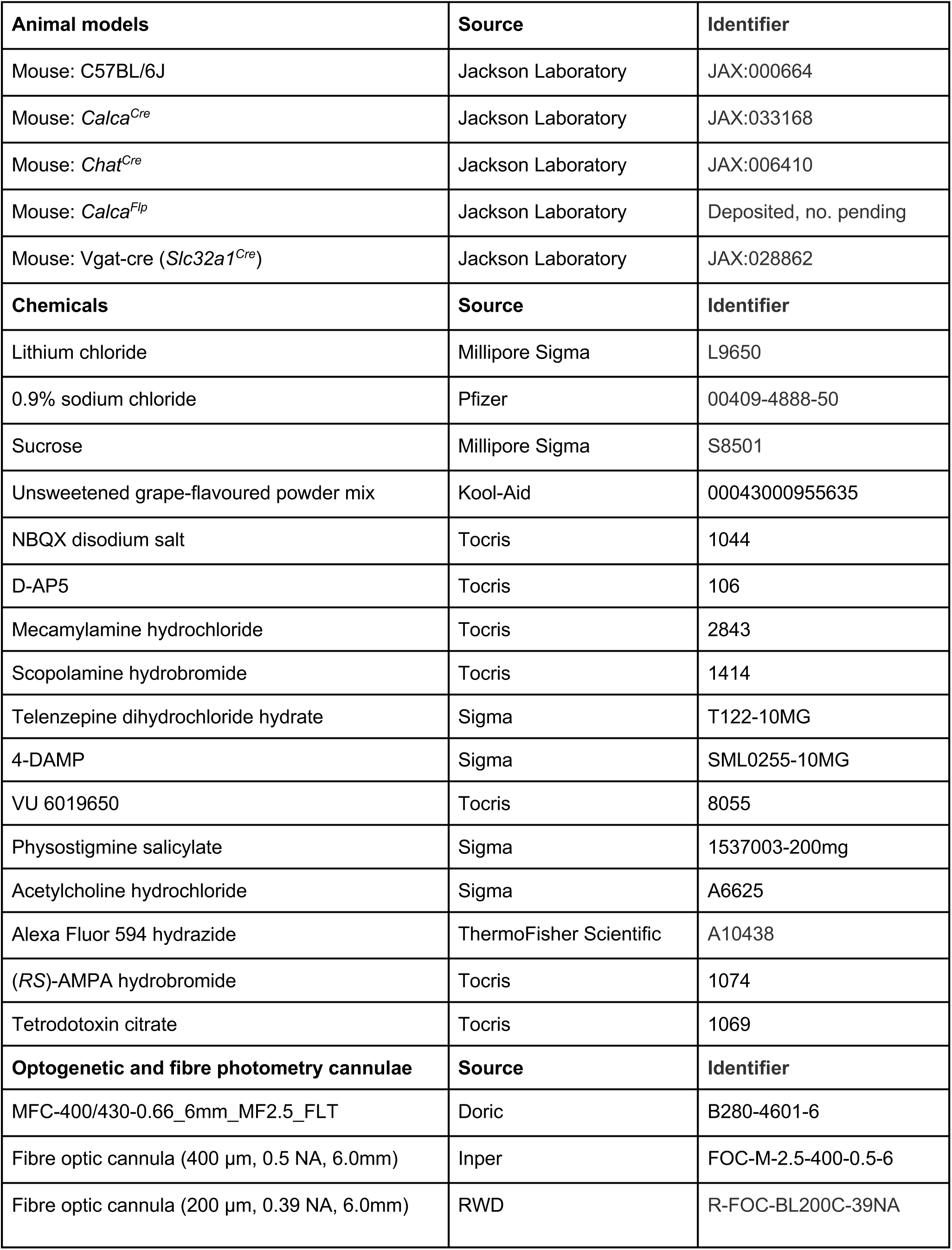
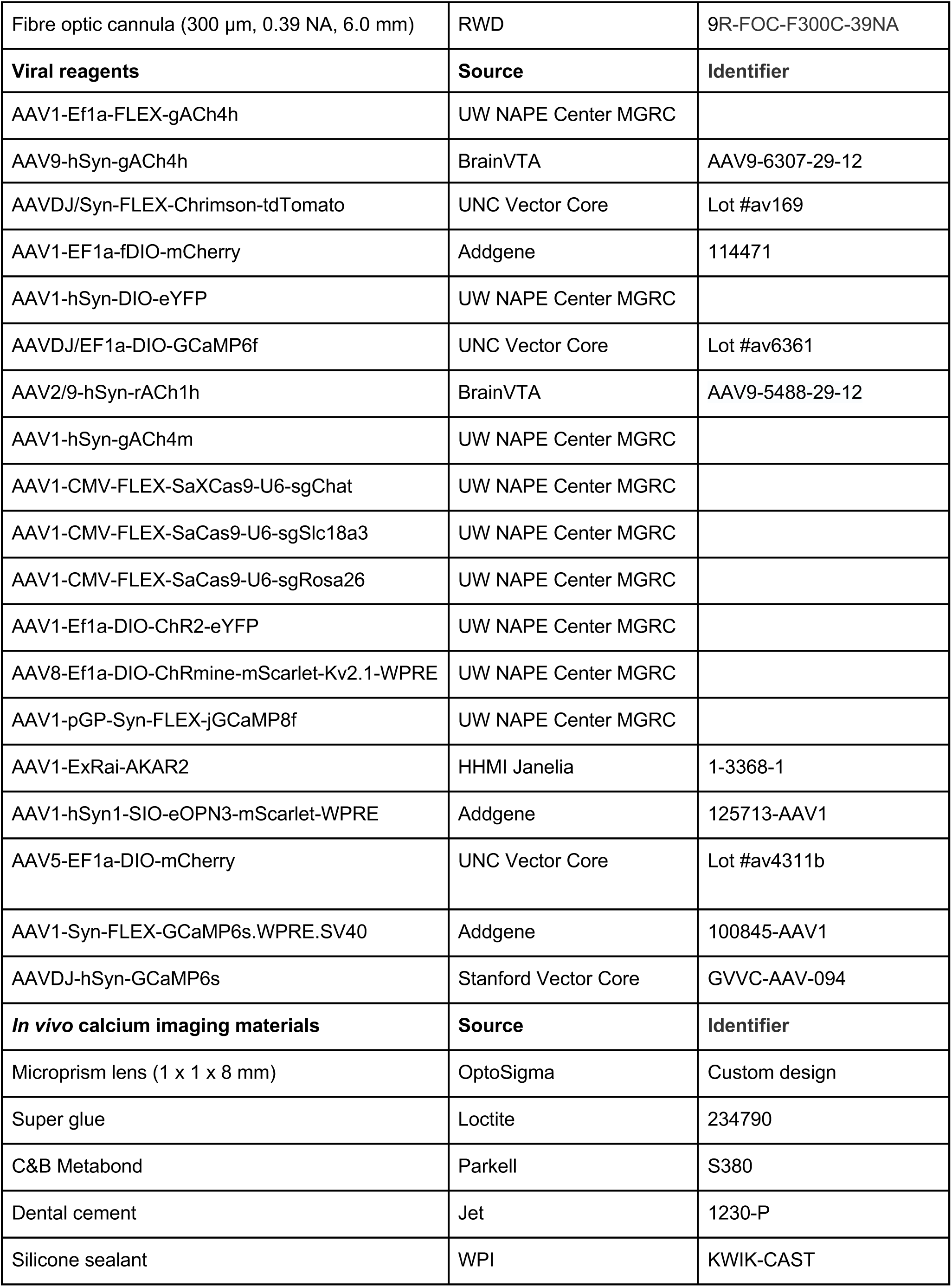

| Animal models | Source | Identifier |
| --- | --- | --- |
| Mouse: C57BL/6J | Jackson Laboratory | JAX:000664 |
| Mouse: *Calca*<sup>Cre</sup> | Jackson Laboratory | JAX:033168 |
| Mouse: *Chat*<sup>Cre</sup> | Jackson Laboratory | JAX:006410 |
| Mouse: *Calca*<sup>Flp</sup> | Jackson Laboratory | Deposited, no. pending |
| Mouse: Vgat-cre (*Slc32a1*<sup>Cre</sup>) | Jackson Laboratory | JAX:028862 |
| Chemicals | Source | Identifier |
| Lithium chloride | Millipore Sigma | L9650 |
| 0.9% sodium chloride | Pfizer | 00409-4888-50 |
| Sucrose | Millipore Sigma | S8501 |
| Unsweetened grape-flavoured powder mix | Kool-Aid | 00043000955635 |
| NBQX disodium salt | Tocris | 1044 |
| D-AP5 | Tocris | 106 |
| Mecamylamine hydrochloride | Tocris | 2843 |
| Scopolamine hydrobromide | Tocris | 1414 |
| Telenzepine dihydrochloride hydrate | Sigma | T122-10MG |
| 4-DAMP | Sigma | SML0255-10MG |
| VU 6019650 | Tocris | 8055 |
| Physostigmine salicylate | Sigma | 1537003-200mg |
| Acetylcholine hydrochloride | Sigma | A6625 |
| Alexa Fluor 594 hydrazide | ThermoFisher Scientific | A10438 |
| (RS)-AMPA hydrobromide | Tocris | 1074 |
| Tetrodotoxin citrate | Tocris | 1069 |
| Optogenetic and fibre photometry cannulae | Source | Identifier |
| MFC-400/430-0.66_6mm_MF2.5_FLT | Doric | B280-4601-6 |
| Fibre optic cannula (400 µm, 0.5 NA, 6.0mm) | Inper | FOC-M-2.5-400-0.5-6 |
| Fibre optic cannula (200 µm, 0.39 NA, 6.0mm) | RWD | R-FOC-BL200C-39NA |
| Fibre optic cannula (300 µm, 0.39 NA, 6.0 mm) | RWD | 9R-FOC-F300C-39NA |
| **Viral reagents** | **Source** | **Identifier** |
| AAV1-Ef1a-FLEX-gACh4h | UW NAPE Center MGRC |  |
| AAV9-hSyn-gACh4h | BrainVTA | AAV9-6307-29-12 |
| AAVDJ/Syn-FLEX-Chrimson-tdTomato | UNC Vector Core | Lot #av169 |
| AAV1-EF1a-fDIO-mCherry | Addgene | 114471 |
| AAV1-hSyn-DIO-eYFP | UW NAPE Center MGRC |  |
| AAVDJ/EF1a-DIO-GCaMP6f | UNC Vector Core | Lot #av6361 |
| AAV2/9-hSyn-rACh1h | BrainVTA | AAV9-5488-29-12 |
| AAV1-hSyn-gACh4m | UW NAPE Center MGRC |  |
| AAV1-CMV-FLEX-SaXCas9-U6-sgChat | UW NAPE Center MGRC |  |
| AAV1-CMV-FLEX-SaCas9-U6-sgSlc18a3 | UW NAPE Center MGRC |  |
| AAV1-CMV-FLEX-SaCas9-U6-sgRosa26 | UW NAPE Center MGRC |  |
| AAV1-Ef1a-DIO-ChR2-eYFP | UW NAPE Center MGRC |  |
| AAV8-Ef1a-DIO-ChRmine-mScarlet-Kv2.1-WPRE | UW NAPE Center MGRC |  |
| AAV1-pGP-Syn-FLEX-jGCaMP8f | UW NAPE Center MGRC |  |
| AAV1-ExRai-AKAR2 | HHMI Janelia | 1-3368-1 |
| AAV1-hSyn1-SIO-eOPN3-mScarlet-WPRE | Addgene | 125713-AAV1 |
| AAV5-EF1a-DIO-mCherry | UNC Vector Core | Lot #av4311b |
| AAV1-Syn-FLEX-GCaMP6s.WPRE.SV40 | Addgene | 100845-AAV1 |
| AAVDJ-hSyn-GCaMP6s | Stanford Vector Core | GVVC-AAV-094 |
| ***In vivo* calcium imaging materials** | **Source** | **Identifier** |
| Microprism lens (1 x 1 x 8 mm) | OptoSigma | Custom design |
| Super glue | Loctite | 234790 |
| C&B Metabond | Parkell | S380 |
| Dental cement | Jet | 1230-P |
| Silicone sealant | WPI | KWIK-CAST |

## Acknowledgements

We thank Y. Li for ACh sensor constructs; C. Zhou, S. Piantadosi, and M. Hjort for help troubleshooting deep brain prism experiments; S. Schattauer, and the staff at the Molecular Genetics Resource Core for the Center in Neurobiology of Addiction, Pain, and Emotion for AAV production; J. Chen, I. Witten, and C. Zimmerman for discussions on design of CFA experiments; and S. Phelps, L. Anastas, and K. Mandeville for technical support. This work was supported in part by NIH (R37 DA032750, R01 DA038168, G.D.S; F30 DA061559-03, C.T.B.) and NIDA (K99 DA059612-02, A.G.G.).

## Author Contributions

W.F. conceived the project with input from G.S. and R.P. W.F., G.S., and R.P. designed and interpreted the experiments. G.S. and R.P. supervised all aspects of the project. W.F. performed the experiments, analysed the data, and generated the figures with contributions from all the authors as described below. W.F., with support from A.G. and R.L., performed behavioural experiments. J.P. and K.I., with support from W.F., performed histology and developed the pipeline for post-hoc atlas registration of prism imaging ROIs. S.P., with support from W.F., performed the snRNAseq experiment. C.B. analysed the data from the snRNAseq experiment. M.T. and W.F. performed *ex vivo* patch clamp electrophysiology experiments. W.F., R.P, and G.S. wrote the paper with input from all the authors.

## Notice of pre-existing conditions, requirements and licenses for article submission

The article submitted together with this notice is subject to the Immediate Access to Research policy of the Howard Hughes Medical Institute (“HHMI”). In accordance with this policy: (i) a preprint of this article either has been, or will be, deposited on a preprint server under a Creative Commons Attribution 4.0 International (CC BY 4.0) license and (ii) an additional author-published revised version of this article incorporating peer review feedback and/or new results or analysis either has been, or prior to journal publication will be, deposited on a preprint server under a CC BY 4.0 license. In addition, a non-exclusive CC BY 4.0 license to this article has been granted to the public and HHMI has a sublicensable, non-exclusive license to this article.

**THIS ARTICLE IS SUBMITTED FOR REVIEW AND ACCEPTANCE SUBJECT TO THESE PRE-EXISTING CONDITIONS, REQUIREMENTS AND LICENSES.**

If you have any concerns with any of these pre-existing conditions, requirements or licenses, please contact the corresponding author immediately.

## Extended Data Figures

**Extended Data Fig. 1.**
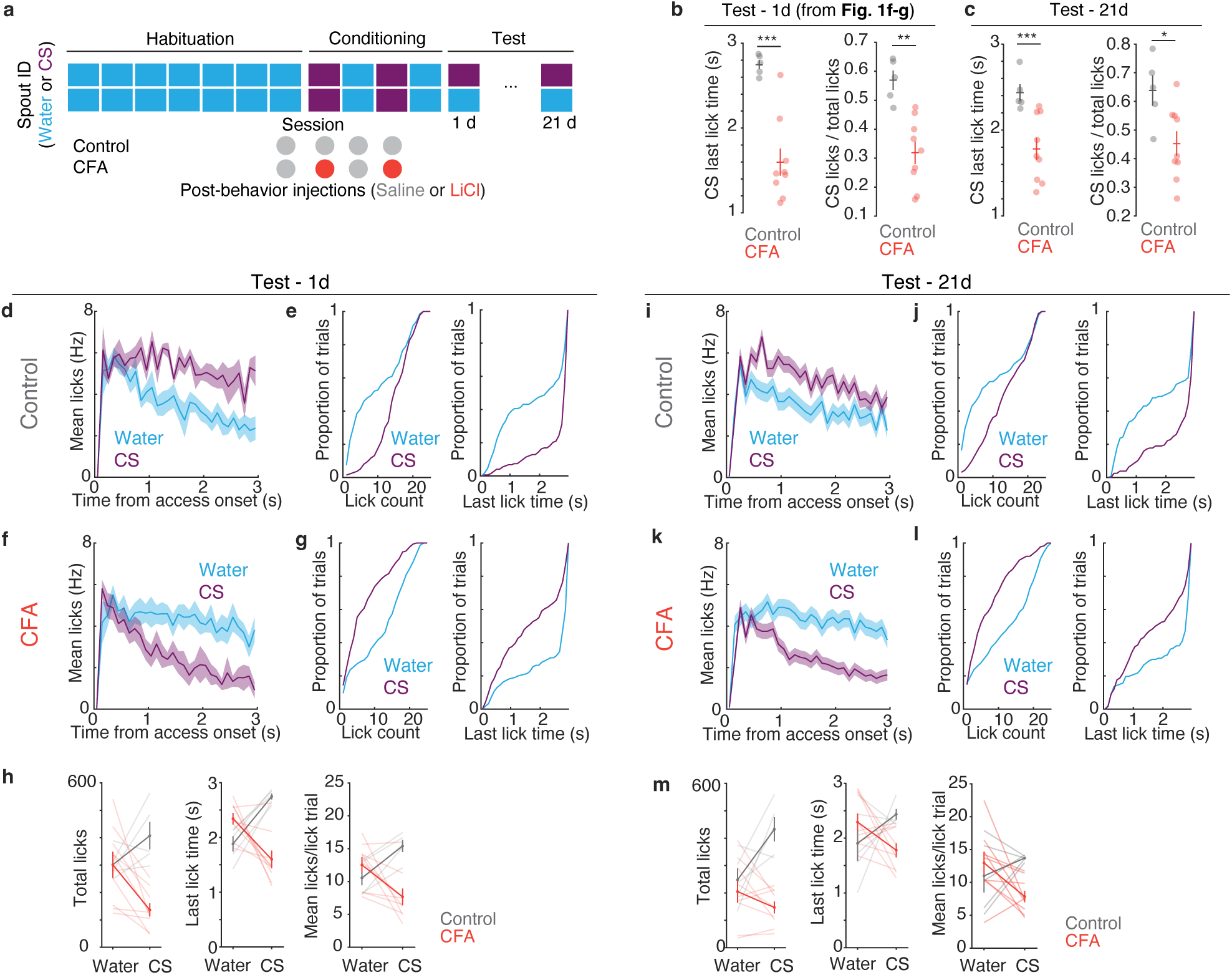
Head-fixed CFA statistics are stable 21 d after conditioning. **a,** Full timeline of CFA experiment. **b,** For visual comparison, from Fig. 1f**-g**, CS last-lick time (left) and CS preference (right) by group at Test 1 d after conditioning. **c,** CS last-lick time (left) and CS preference (right) at Test 21 d after conditioning. **d,** Mean lick rate during lick trials by solution in Control mice at 1 d Test. **e,** Left: Cumulative distribution of lick count in trials with a lick by solution in Control mice at 1 d Test. Right: Cumulative distribution of last-lick time by solution in Controls at 1 d Test. **f,** Mean lick rate during lick trials by solution in CFA mice at 1 d Test. **g,** Left: Cumulative distribution of lick count in trials with a lick by solution in CFA mice at 1 d Test. Right: Cumulative distribution of last-lick time by solution in CFA mice at 1 d Test. **h,** Left: Mean total lick count by solution. Middle: Mean last-lick time by solution. Right: Mean licks per lick trial by solution at 1 d Test. **i,** Mean lick rate during lick trials by solution in Control mice at 21 d Test. **j,** Left: Cumulative distribution of lick count in trials with a lick by solution in Control mice at 21 d Test. Right: Cumulative distribution of last-lick time by solution in Controls at 21 d Test. **k,** Mean lick rate during lick trials by solution in CFA mice at 21 d Test. **l,** Left: Cumulative distribution of lick count in trials with a lick by solution in CFA mice at 21 d Test. Right: Cumulative distribution of last-lick time by solution in CFA mice at 21 d Test. **m,** Left: Mean total lick count by solution. Middle: Mean last-lick time by solution. Right: Mean licks per lick trial by solution at 21 d Test. (Control, n=5 mice; CFA, n=9 mice). Bars represent SEM. \**P*<0.05, \*\**P*<0.01. \*\*\**P*<0.001. See **Supplementary Table 1** for details of statistical tests and exact *P* values.

**Extended Data Fig. 2.**
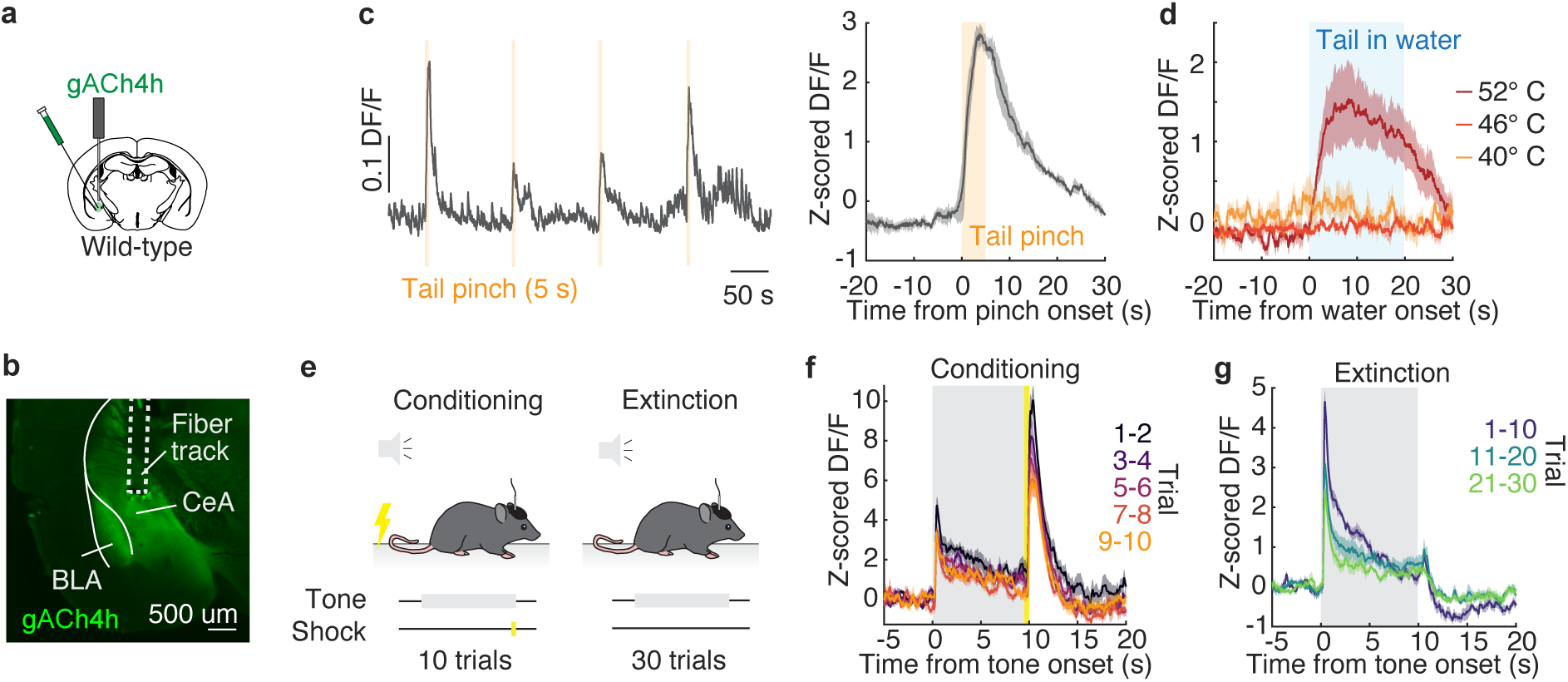
Noxious stimuli drive ACh release in CeA. **a,** Viral strategy for mechanical and noxious photometry experiments. **b,** Example histology of gACh4h and photometry fibre placement in CeA. **c,** Left, Example photometry trace during lightly anaesthetized tail pinch. Tail pinch was applied to the base of the tail via a small bulldog clamp approximately every 2 mins. Right, Average baseline-corrected ACh photometry trace in response to lightly anaesthetized tail pinch (n=3 mice). **d,** Averaged baseline-corrected ACh photometry trace in response to lightly anaesthetized hot water tail immersion (n=3 mice). **e,** Schematic of fear conditioning experiment. **f,** Mean baseline-subtracted ACh photometry trace during Conditioning, plotted by trial number. **g,** Mean baseline-subtracted ACh during Extinction, plotted by trial number (n=6 mice). Shaded regions represent SEM.

**Extended Data Fig. 3.**
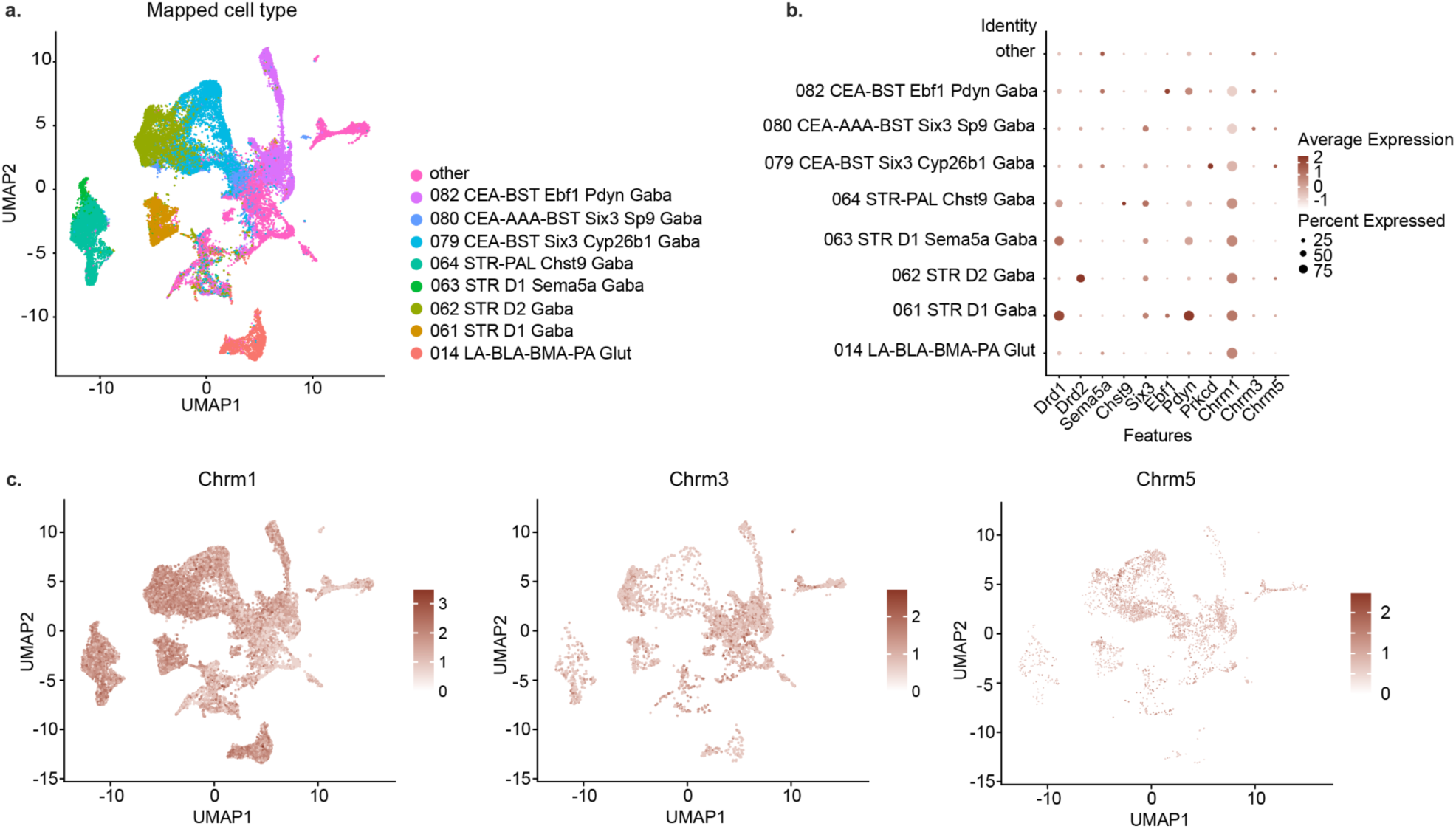
Single-nucleus RNA sequencing reveals G_q_-linked muscarinic receptor expression in CeA. **a,** UMAP plot illustrating nuclei clustered according to transcriptional similarities, as mapped with Allen Brain Institute’s MapMyCells algorithm (using 10x Whole mouse brain taxonomy (CCN20230722)). **b,** Disc plot illustrating the prevalence of selected candidate markers for each cell type, as well as genes for M1, M3, and M5 receptors (*Chrm1*, *Chrm3*, and *Chrm5*)**. c,** Feature plots indicating the expression of genes for M1, M3, and M5 receptors.

**Extended Data Fig. 4.**
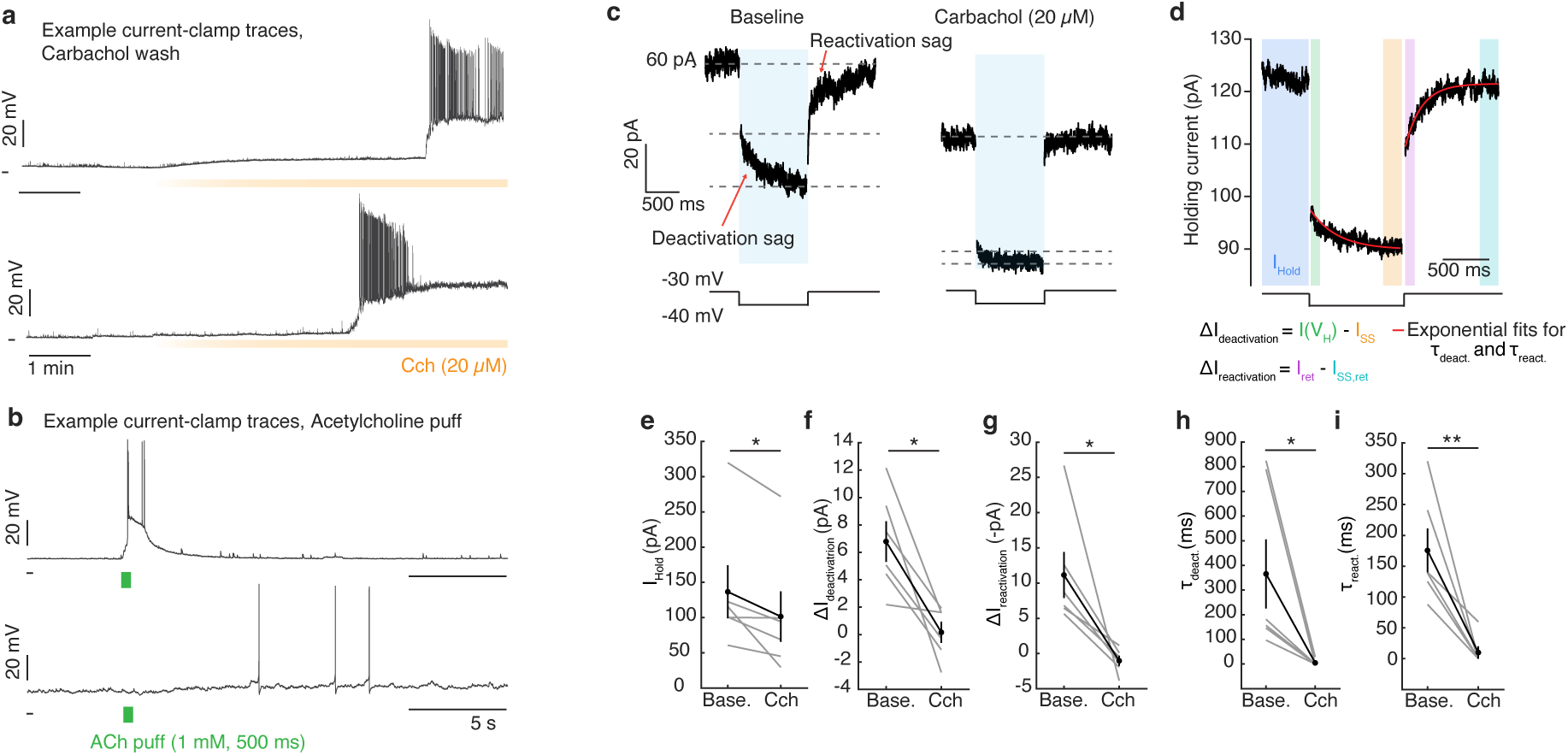
Cholinergic inactivation of M-currents at CeA neurons. **a,** Example traces from CeA neurons during bath application of carbachol (Cch, 20 µM). **b,** Example traces from CeA neurons during micro-puffs of acetylcholine (ACh, 1 mM). ACh puff experiments were performed in the presence of physostigmine (1 µM). **c,** Example traces from a CeA neuron during M-current assay illustrating characteristic sags resulting from slow deactivation and reactivation of Kv7 channels around a hyperpolarizing voltage step when held at -30 mV. Cch reduces the holding current and abolishes sags around the voltage step. **d,** Illustration of windows used for calculation of holding current (I_Hold_), deactivation current (ΔI_deactivation_), and reactivation current (ΔI_reactivation_); and exponential fit lines used to calculate time constant (τ) values around voltage step. **e,** Mean holding current prior to voltage step. **f,** Mean deactivation sag current during voltage step. **g,** Mean deactivation sag current following voltage step. **h,** Mean time constant of deactivation sag. **i,** Mean time constant of reactivation sag (n = 6 neurons, 3 mice). All experiments were performed in the presence of picrotoxin (100 µM). Bars represent SEM. \**P*<0.05, \*\**P*<0.01.

**Extended Data Fig. 5.**
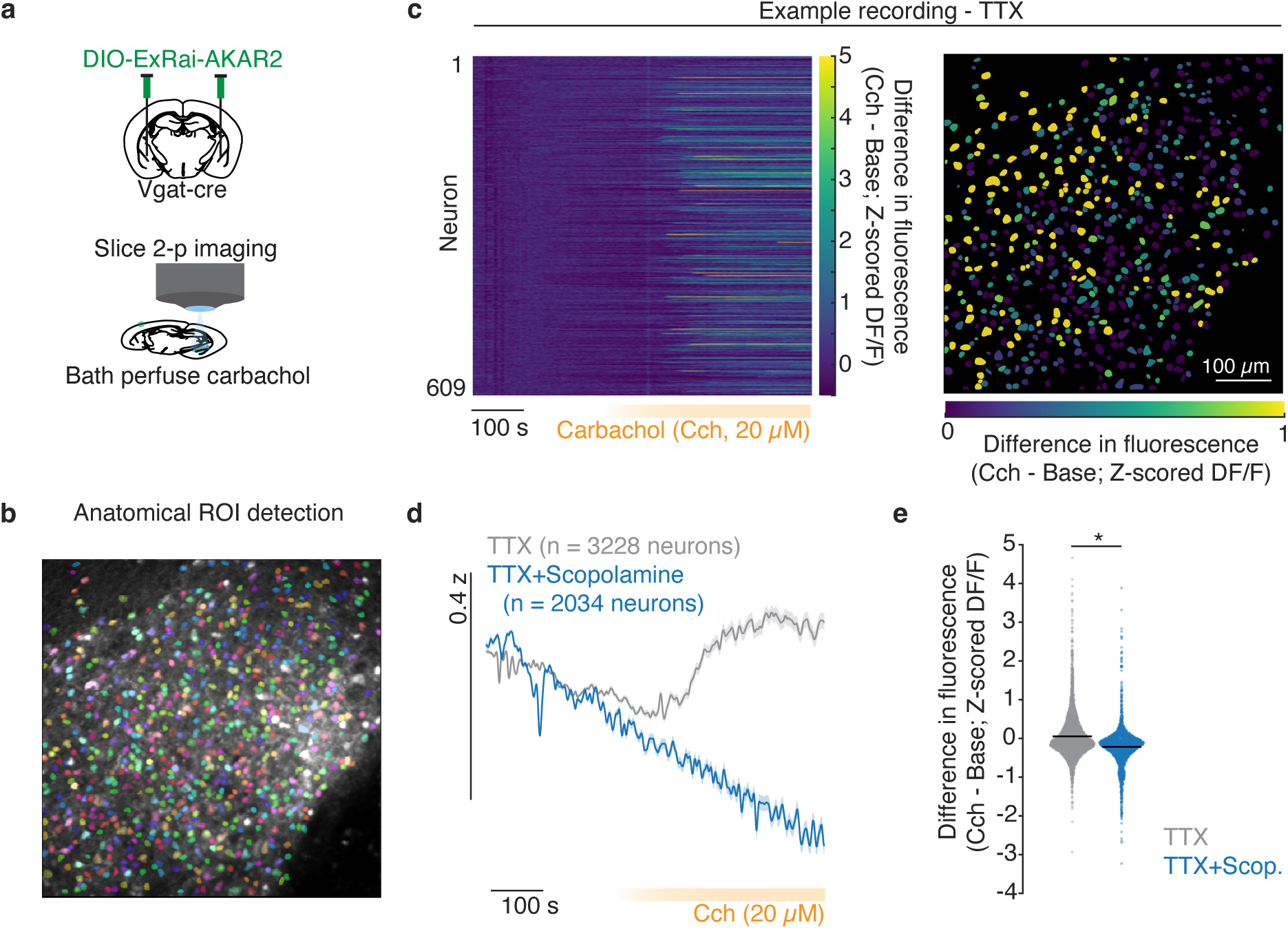
Muscarinic activation of protein kinase A (PKA) in CeA neurons. **a,** Schematic of approach for slice 2-photon PKA sensor imaging in CeA. **b,** Example mean fluorescence image of PKA sensor recording with coloured regions-of-interest identified with CellPose within Suite2p. **c,** Left: Example heatmap of PKA sensor fluorescence during Cch wash-in experiment in the presence of tetrodotoxin (TTX, 1 µM). Right: Example mean baseline-subtracted fluorescence during Cch wash by neuron. **d,** Mean fluorescence across all neurons by treatment group during Cch wash (TTX, n=3328 neurons, 8 slices; Scopolamine (10 µM) + TTX, n=2034 neurons, 5 slices). **e,** Mean baseline-subtracted fluorescence during Cch wash by group. Shaded regions represent SEM. \**P*<0.05. See **Supplementary Table 1** for details of statistical tests and exact *P* values.

**Extended Data Fig. 6.**
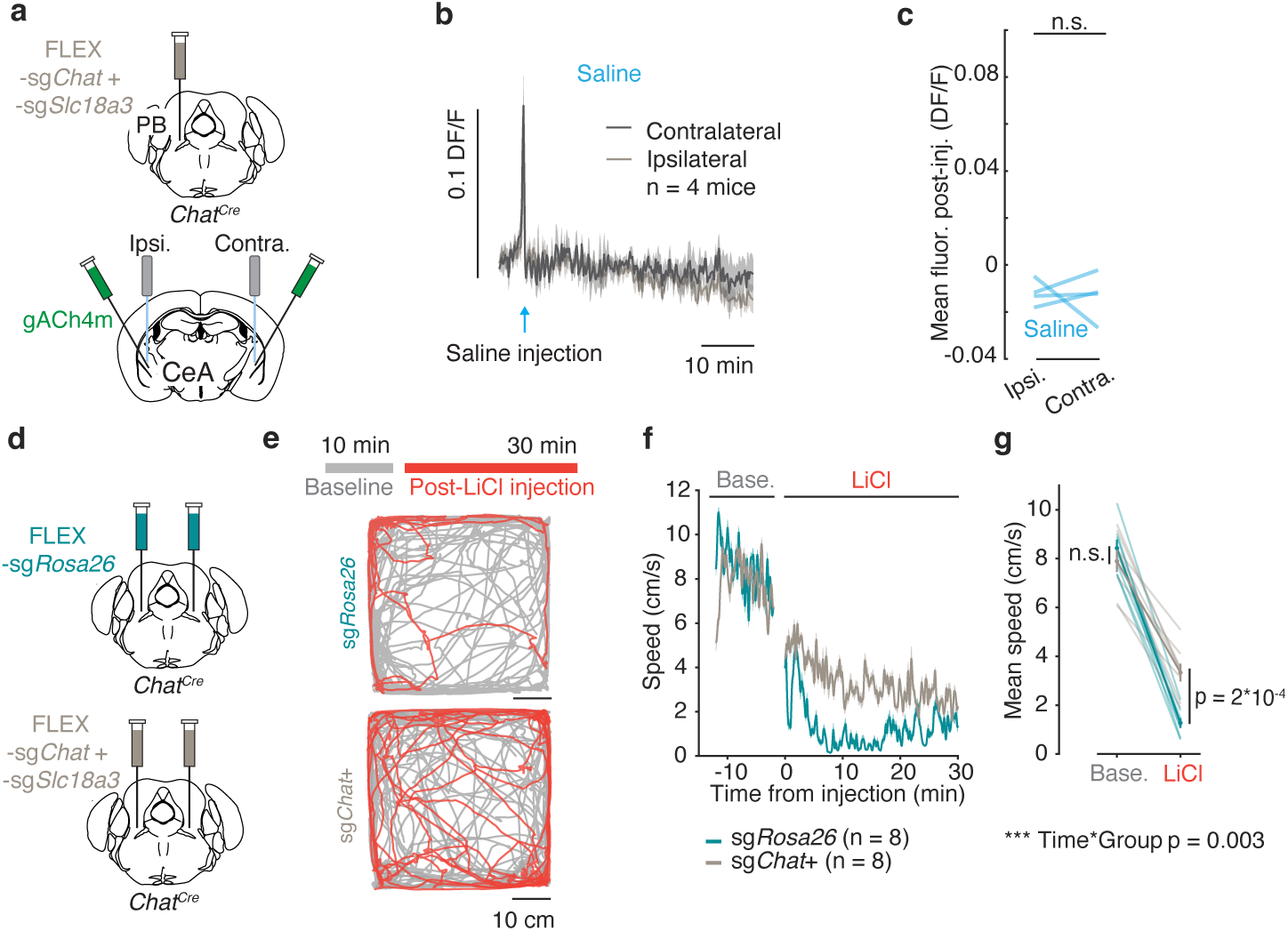
PB^ACh^ knockdown attenuates malaise phenotype following LiCl injection. **a,** Viral strategy for bilateral CeA ACh sensor recording following unilateral PB^ACh^ knockdown (from Fig. 4.) **b,** Mean ACh sensor fluorescence during saline injection in CeA ipsilateral (pink, Ipsi.) and contralateral (Contra., grey) to site of knockdown viral injection. **c,** Mean post-injection ACh sensor fluorescence by recording site and injection type. **d,** Viral strategy for bilateral knockdown of *Chat* and *Slc18a3* in PB*^Chat^* neurons for CFA behavioural testing. **e,** Top: Timeline of open field injection experiment. Bottom: Example tracks in open field chamber before (grey) and after (red) LiCl injection. **f,** Group mean speed during open field experiment. **g,** Mean speed before and after LiCl injection by group. \**P*<0.05, \*\**P*<0.01. \*\*\**P*<0.001. See **Supplementary Table 1** for details of statistical tests and exact *P* values.

**Extended Data Fig. 7.**
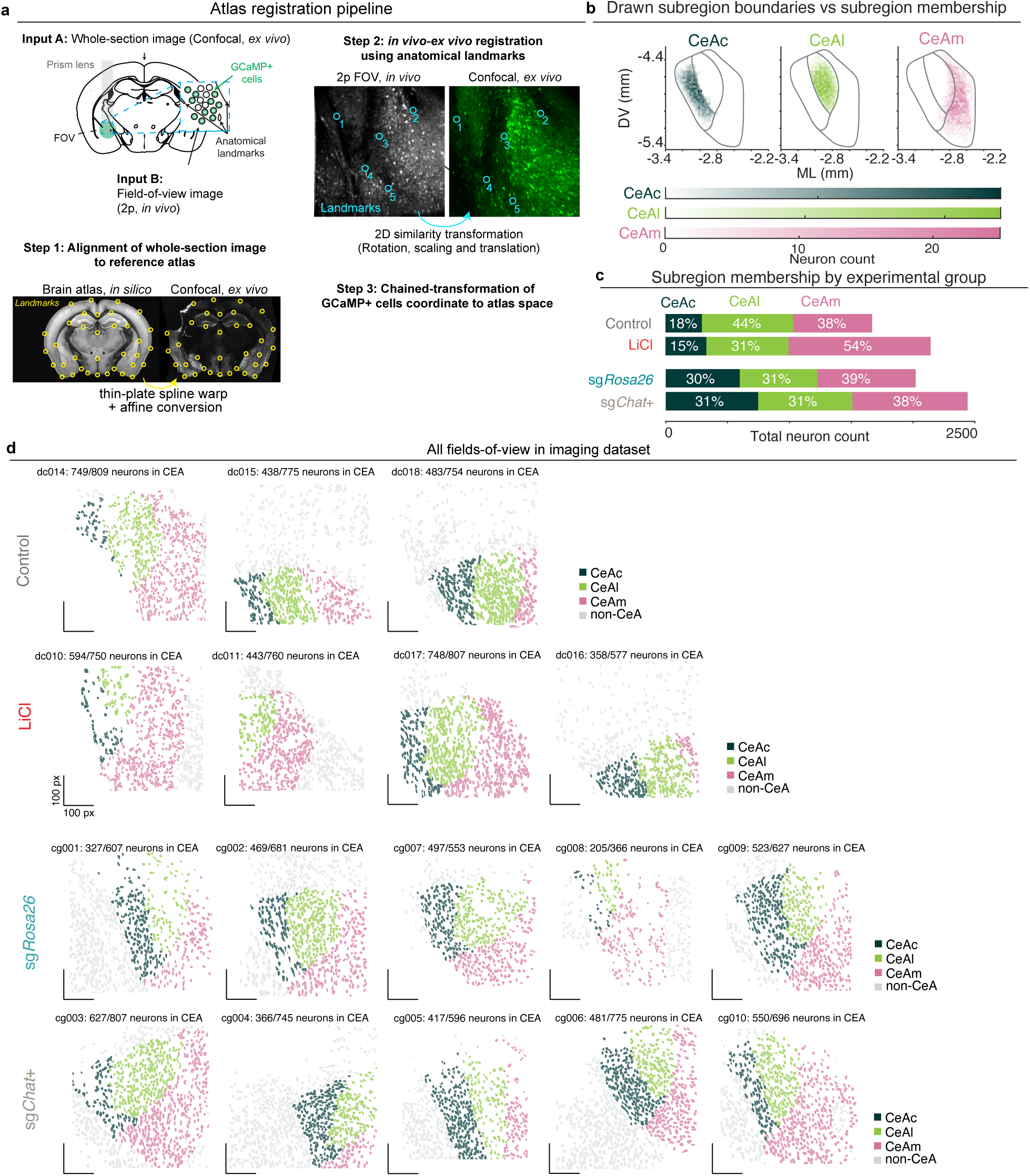
Post-hoc atlas registration of subcortical prism lens 2-photon imaging. **a,** Atlas registration pipeline for deep-brain prism imaging. **b,** Inferred CeA subregion boundaries based on neuronal densities in 0.025 mm^2^ spatial bins in coronal space. Because bins are collapsed along the anterior-posterior axis, spatial bin membership in visualizations does not determine subregion membership (i.e., some spatial bins contain both CeAl and CeAm neurons in heatmaps, but neurons are appropriately split by subregion for analyses). **c,** All fields-of-view from 2p imaging experiments. Neurons are coloured by CeA subregion. Grey neurons were determined to be outside of CeA and were excluded from all analyses.

**Extended Data Fig. 8.**
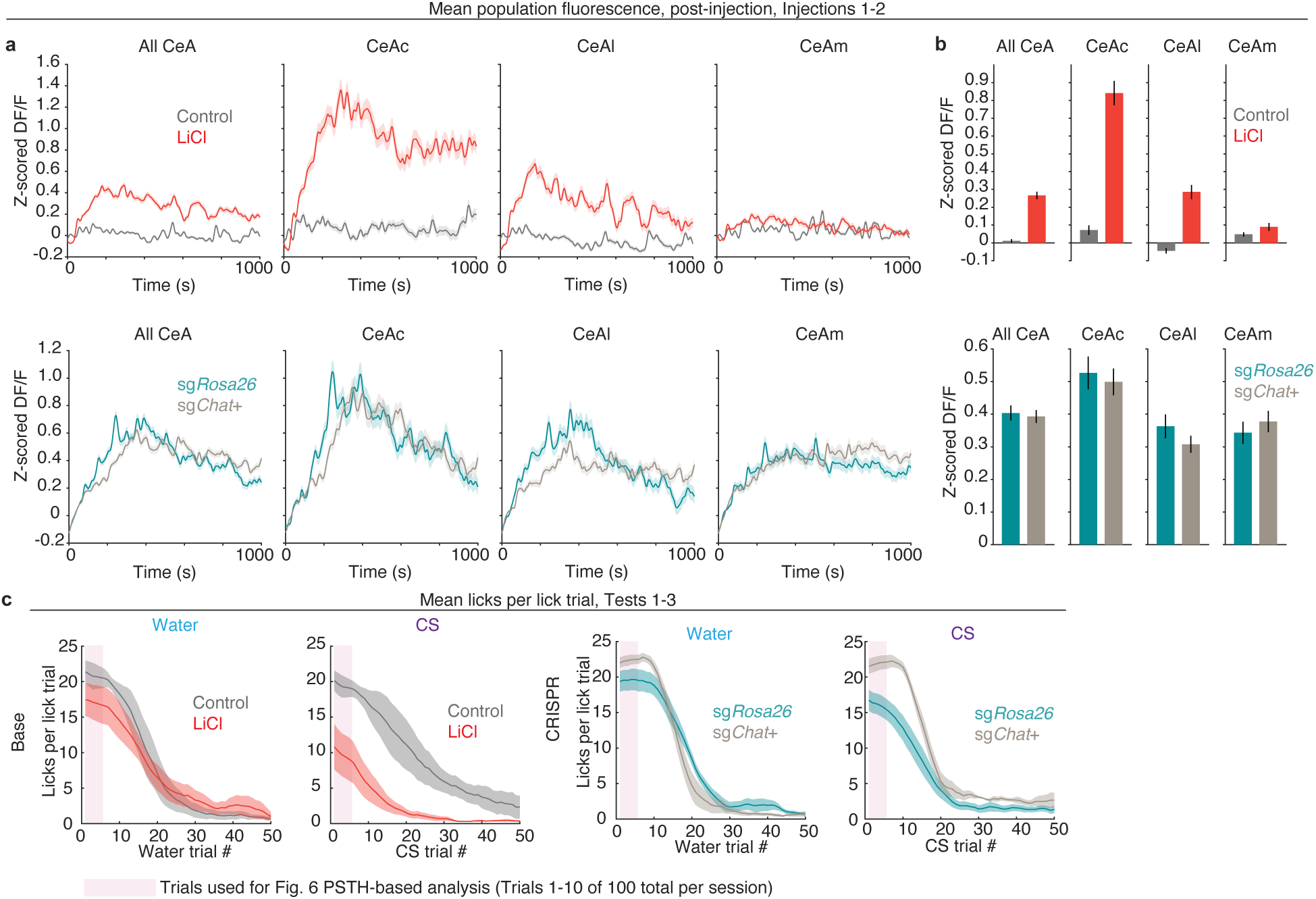
Mean population statistics from 2p experiment. **a,** Top, Mean population fluorescence during post-injection recordings in Base experiment. Bottom, Same as top, for CRISPR experiment. **b**, Top, Mean population fluorescence by CeA subregion. Shaded regions and bars represent SEM over all neurons. **c,** Mean licks per lick trial during Tests 1-3 by solution and group in Base experiment. Bottom, Same as top, for CRISPR experiment. Pink shading denotes trials used for analysis in Fig. 6. Shaded regions represent SEM over mice.

**Extended Data Fig. 9.**
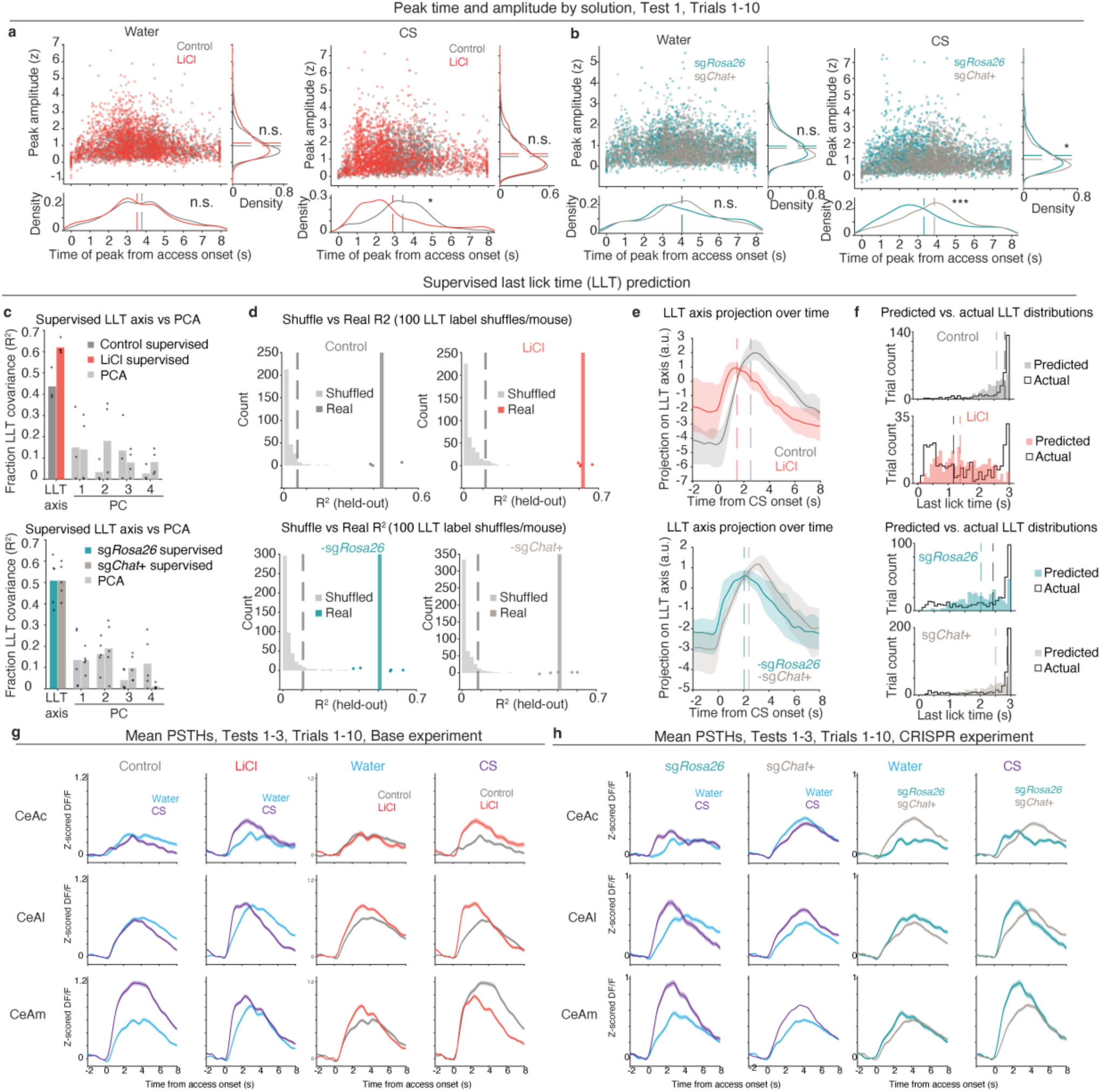
Peak PSTH statistics, validation of supervised last-lick time (LLT) prediction, and mean PSTHs by CeA subregion. **a,** Mean amplitude and time of peak responses during water (left) and CS (right) trials across all CeA neurons’ PSTHs in Base experiment. Line plots describe the relative distributions of peak statistics by group. Statistical tests were performed on actual peak statistic values. **b,** Same as **a,** for CRISPR experiment. **c,** Comparison of prediction of last-lick time (LLT) with neural data when using supervised approach (LLT axis) vs individual principal components (PCs) in PCA-based approach. Fraction LLT covariance describes mean trial-by-trial correlation of predicted vs actual LLTs. **d**, Comparison of supervised LLT prediction when last-lick time labels are shuffled vs when real labels are retained. Individual points represent mean values per mouse. Vertical dashed line represents 95th percentile of shuffled prediction performance; vertical solid line represents mean of performance using real labels. **e**, Visualization of LLT axis values when projected along trial time, using weights calculated from a single fixed window during access period. Vertical dashed lines represent real mean last-lick times. LLT axis value peaks correspond to most likely true last-lick time. **f**, Comparison of distributions of predicted and actual distributions of last-lick times. **g,** Solution and group comparisons of mean PSTHs by CeA subregion for Base experiment. **h,** Same as **g,** but for CRISPR experiment. \**P*<0.05, \*\**P*<0.01. \*\*\**P*<0.001. See **Supplementary Table 1** for details of statistical tests and exact *P* values.

**Extended Data Fig. 10.**
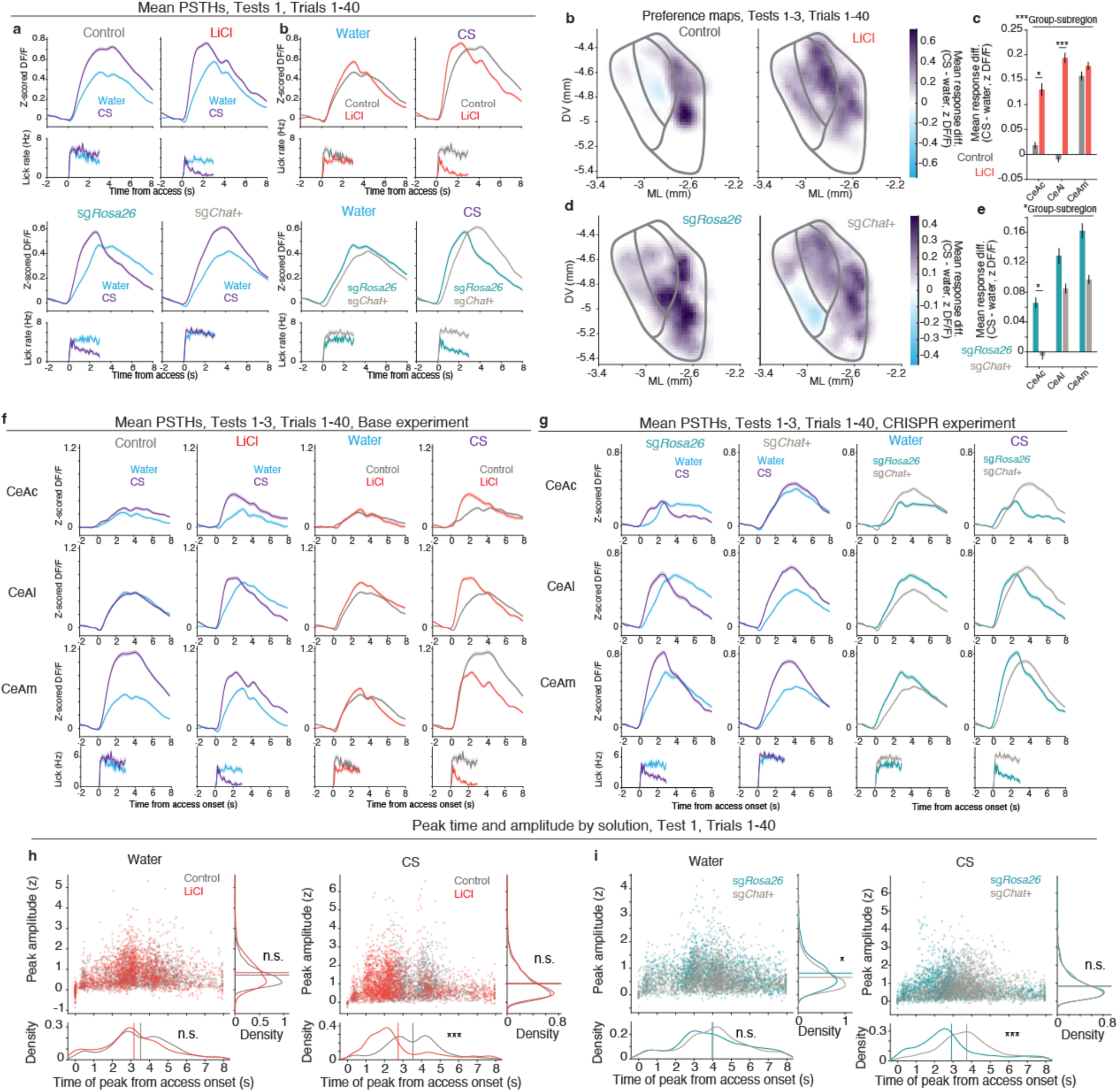
Analysis of CFA Test data using the first 40 trials of Test sessions. **a,** Mean neural responses across all CeA neurons by group during water and CS lick trials at Test 1 in Base (top) and CRISPR (bottom) experiments over trials 1-40 (analogous to Fig. 6c**,g**). **b,** Spatial maps of group mean difference in neuronal response to CS versus water (from 0 to 1.5 s after access onset) during Tests 1-3 in Base experiment over trials 1-40 in Base experiment (analogous to Fig. 6m). **c,** Group mean difference in neuronal response to CS versus water by CeA subregion in Base (top) and CRISPR (bottom) experiments (analogous to Fig. 6n). **d,** Same as **b**, for CRISPR experiment (analogous to Fig. 6o). **e,** Same as **c**, for CRISPR experiment (analogous to Fig. 6p). **f,** Solution and group comparisons of mean PSTHs by CeA subregion for Base experiment. **g,** Same as **f,** but for CRISPR experiment. **h,** Mean amplitude and time of peak responses during water (left) and CS (right) trials across all CeA neurons’ PSTHs in Base experiment. Line plots describe the relative distributions of peak statistics by group. Statistical tests were performed on actual peak statistic values. **i,** Same as **h,** for CRISPR experiment. \**P*<0.05, \*\**P*<0.01. \*\*\**P*<0.001. See **Supplementary Table 1** for details of statistical tests and exact *P* values.

**Supplementary Table 1.**
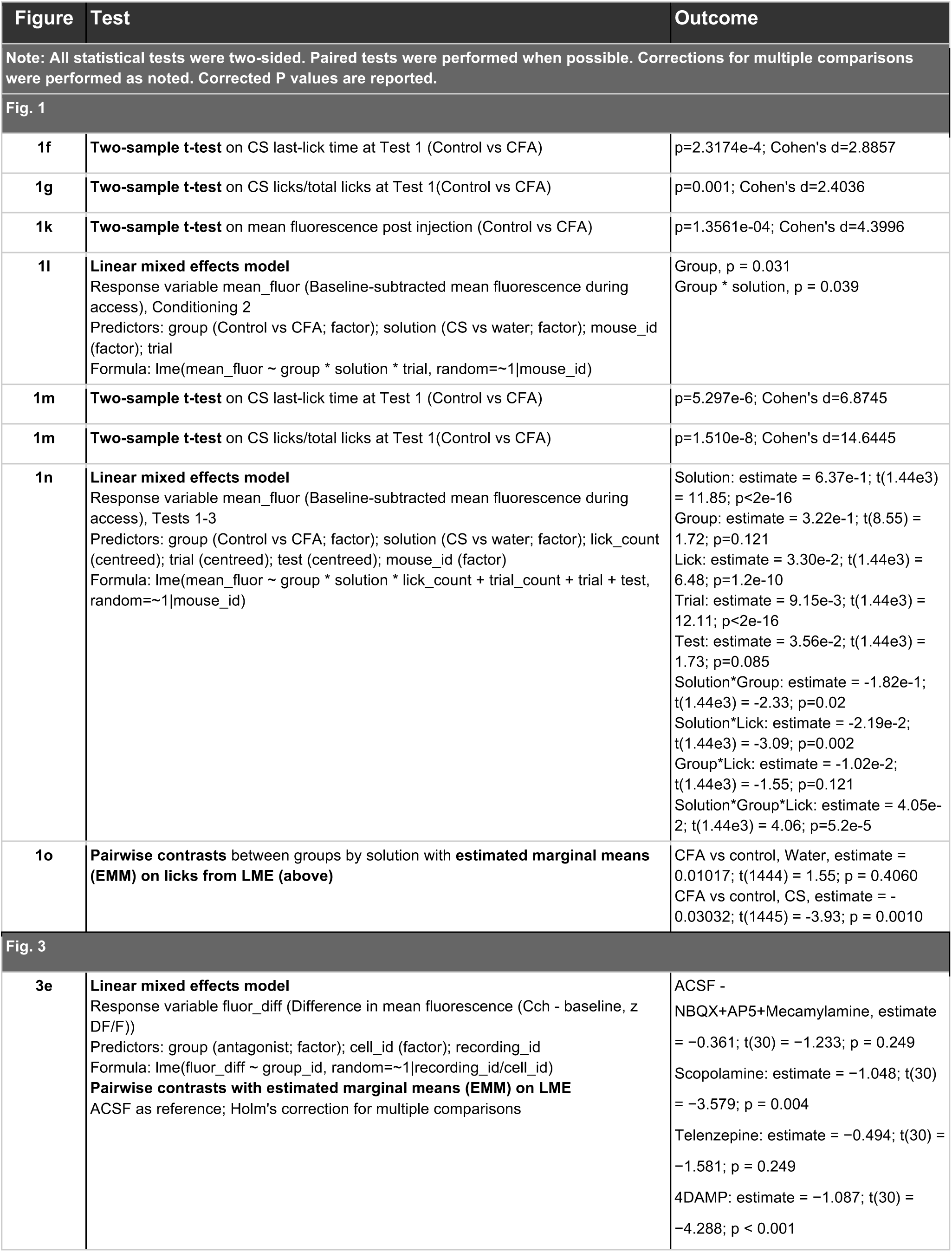
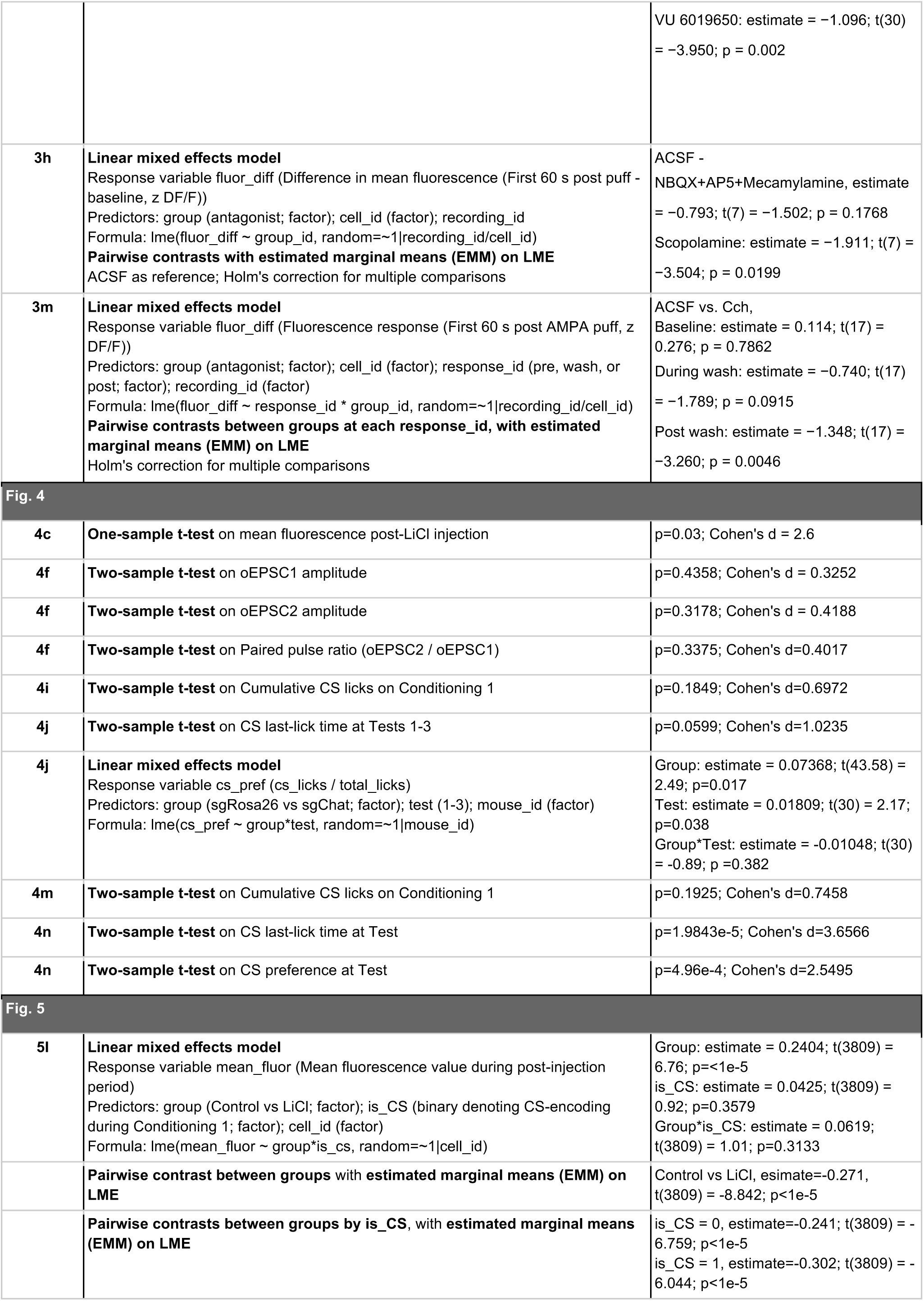
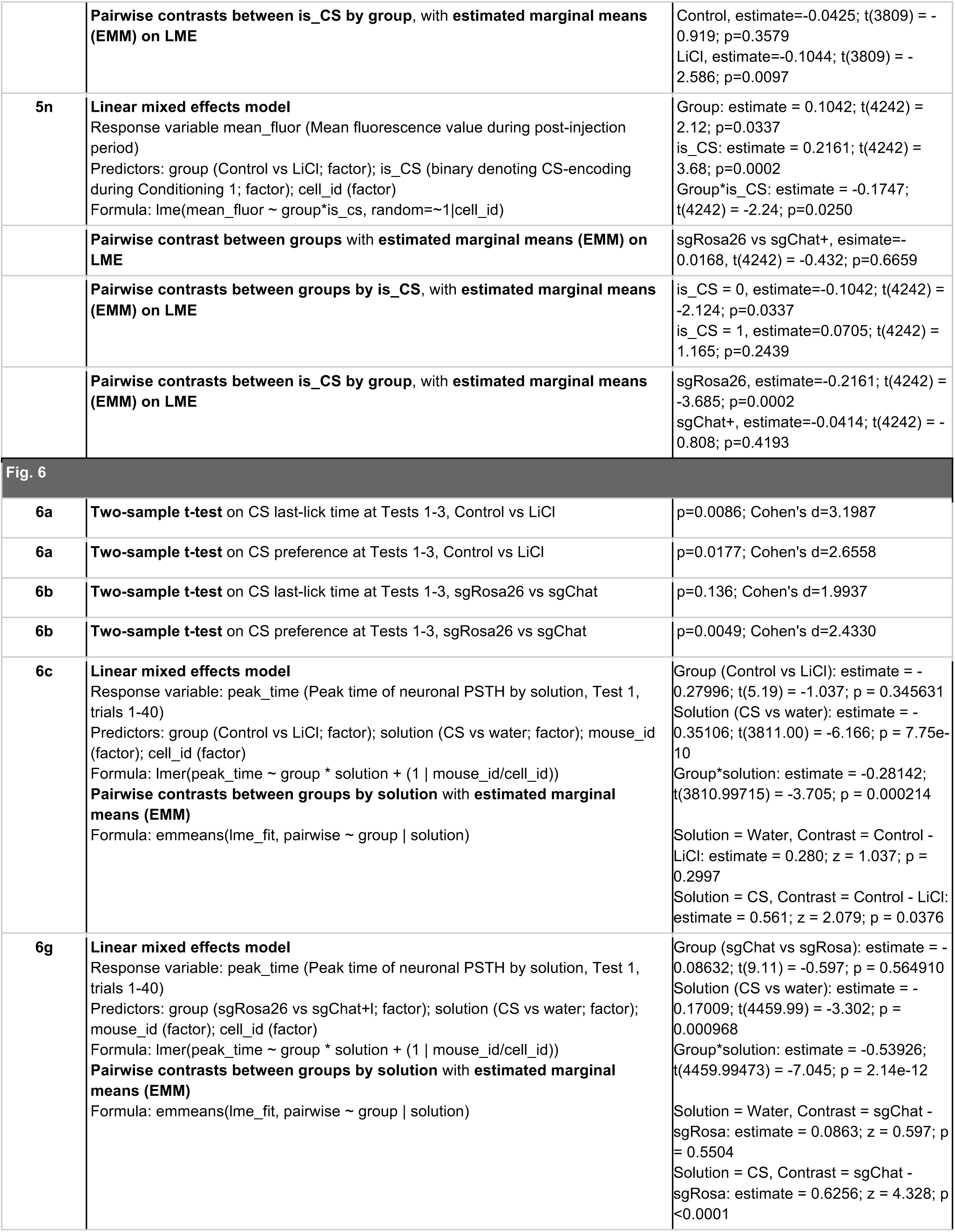
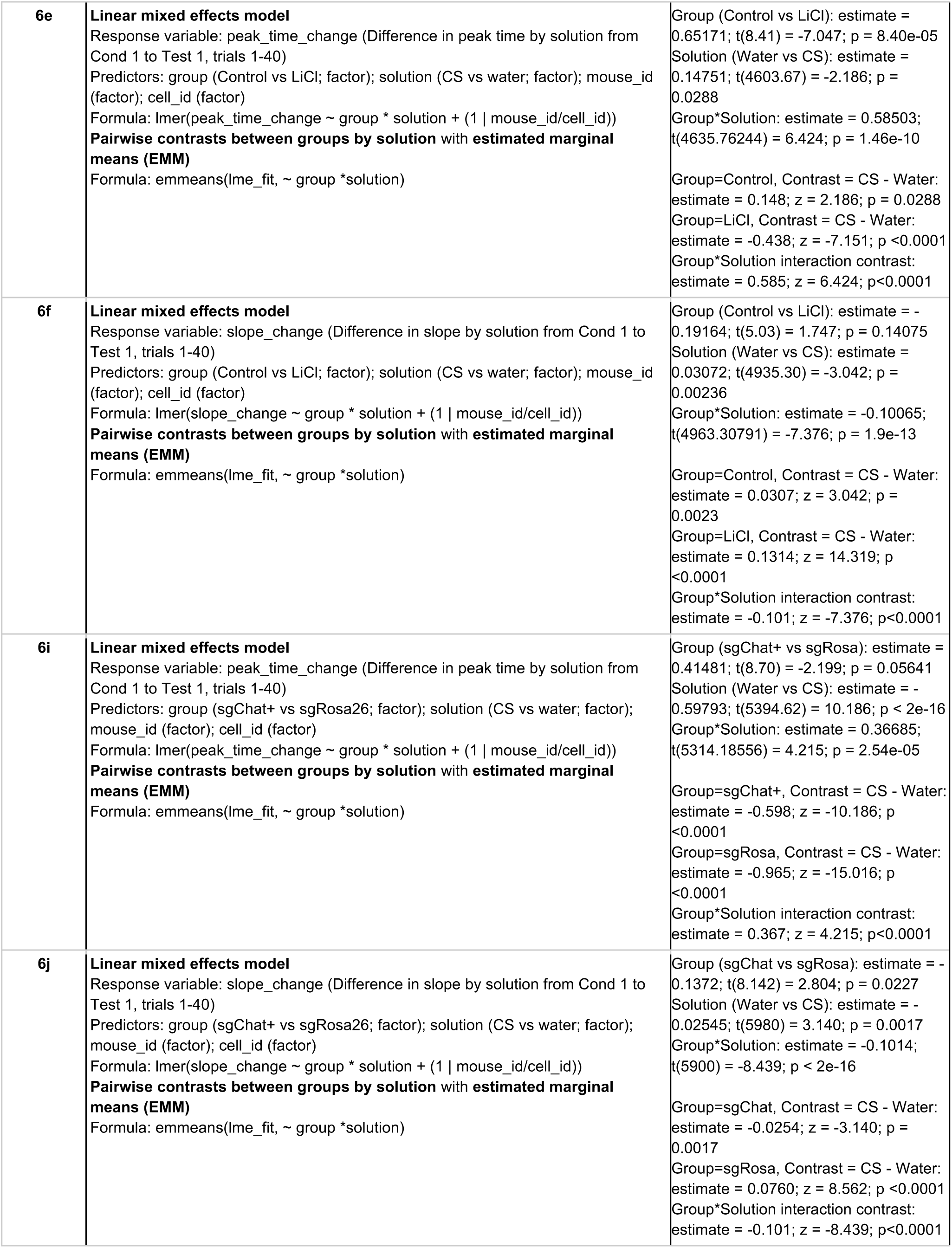
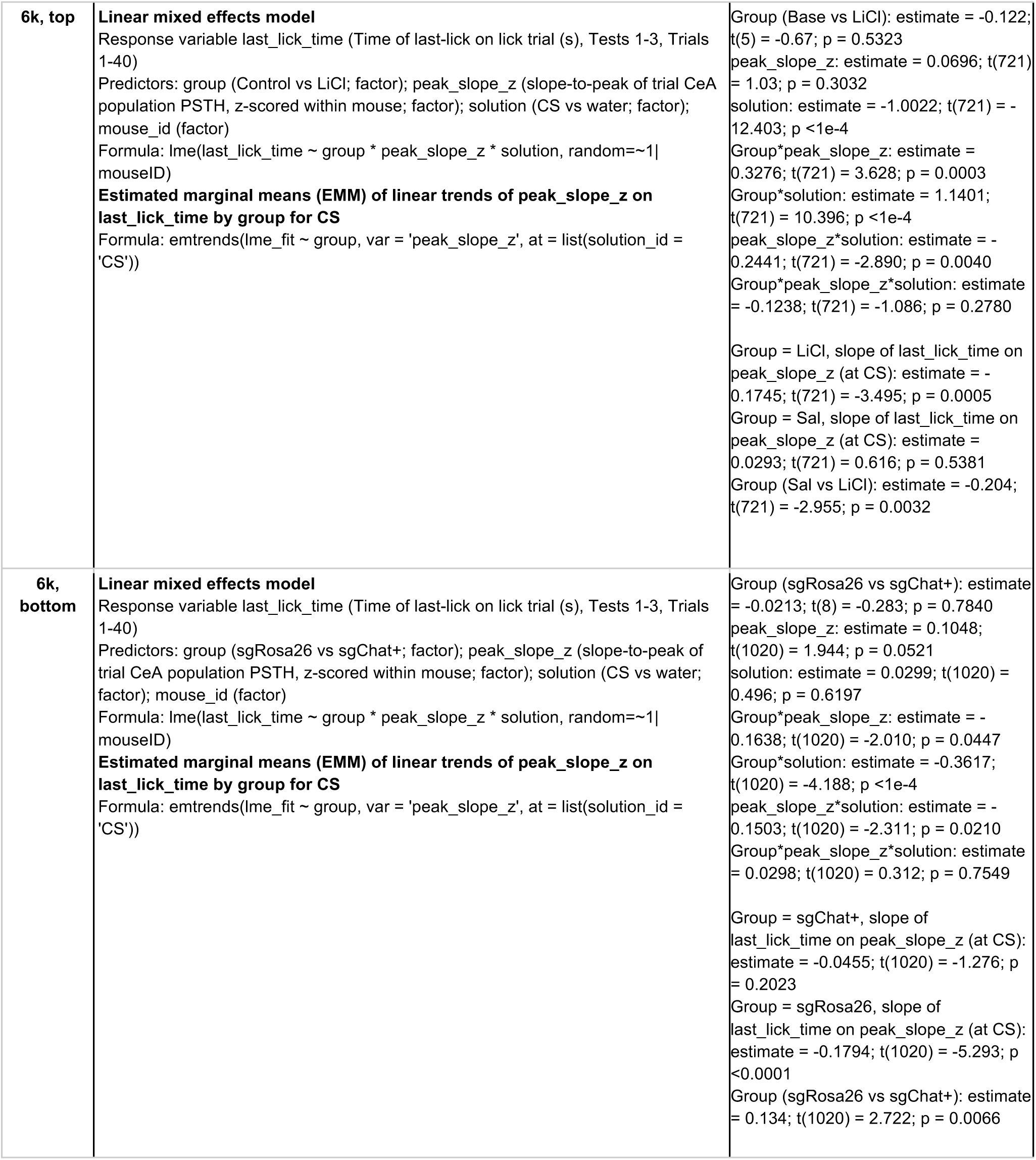
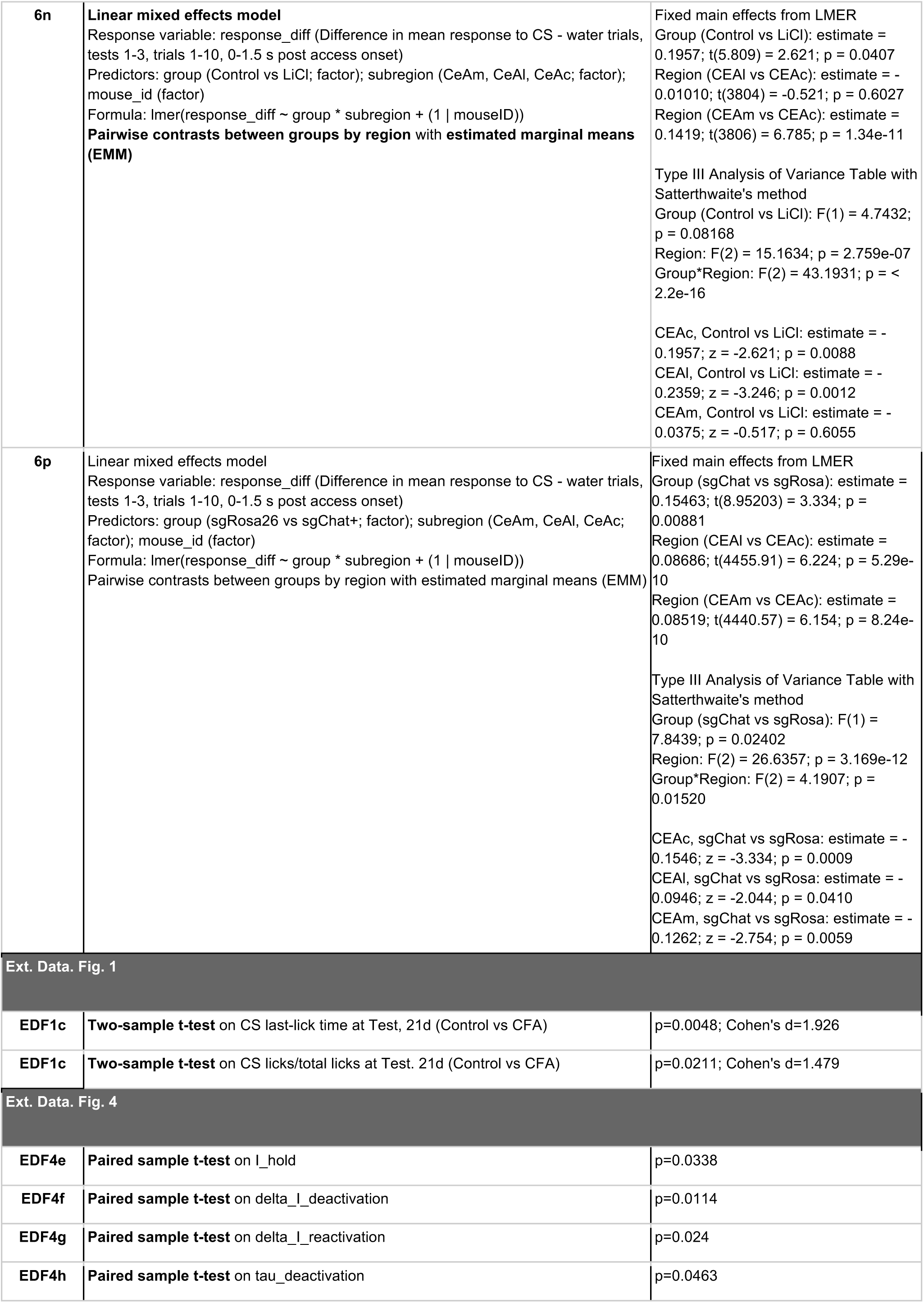
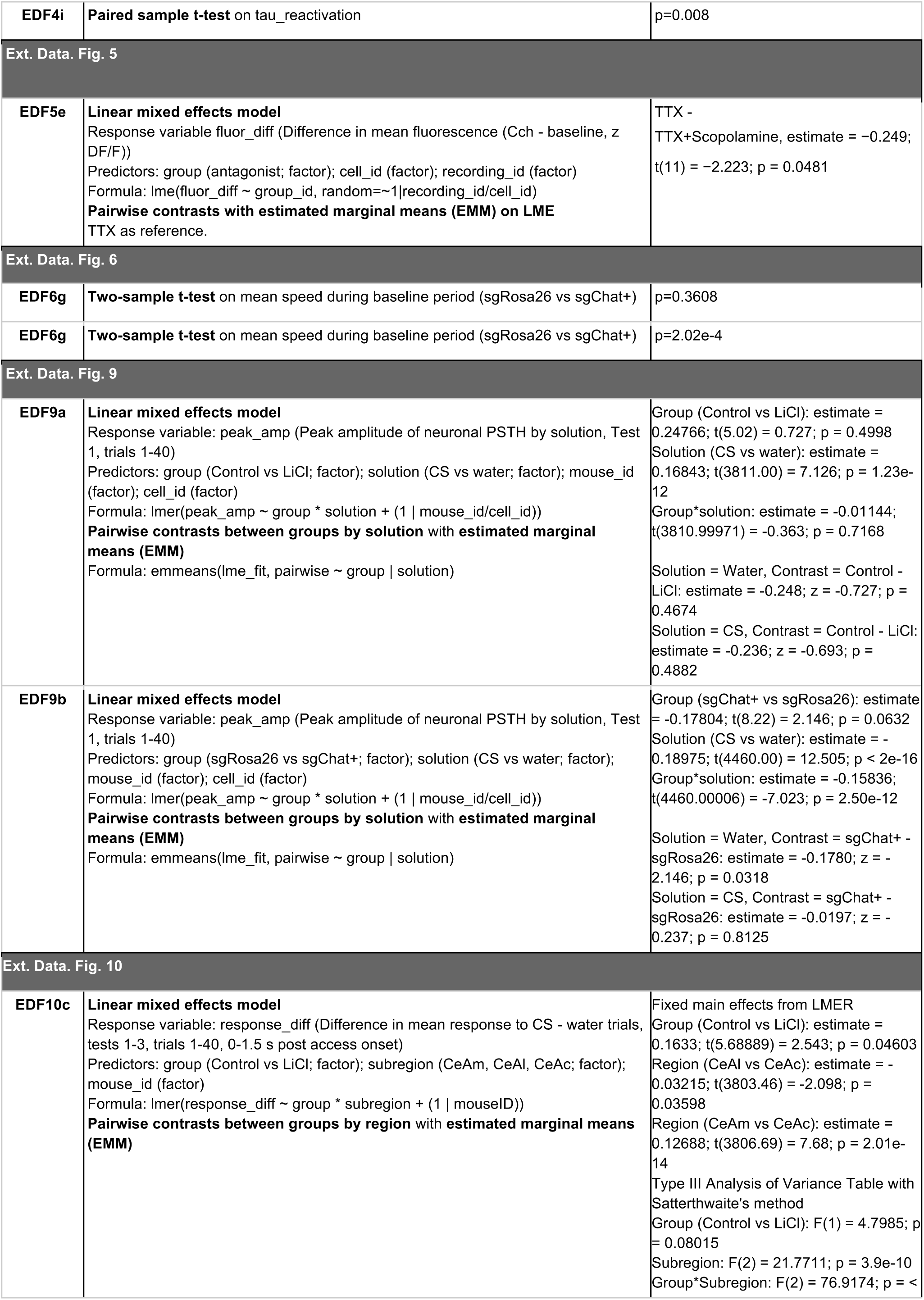
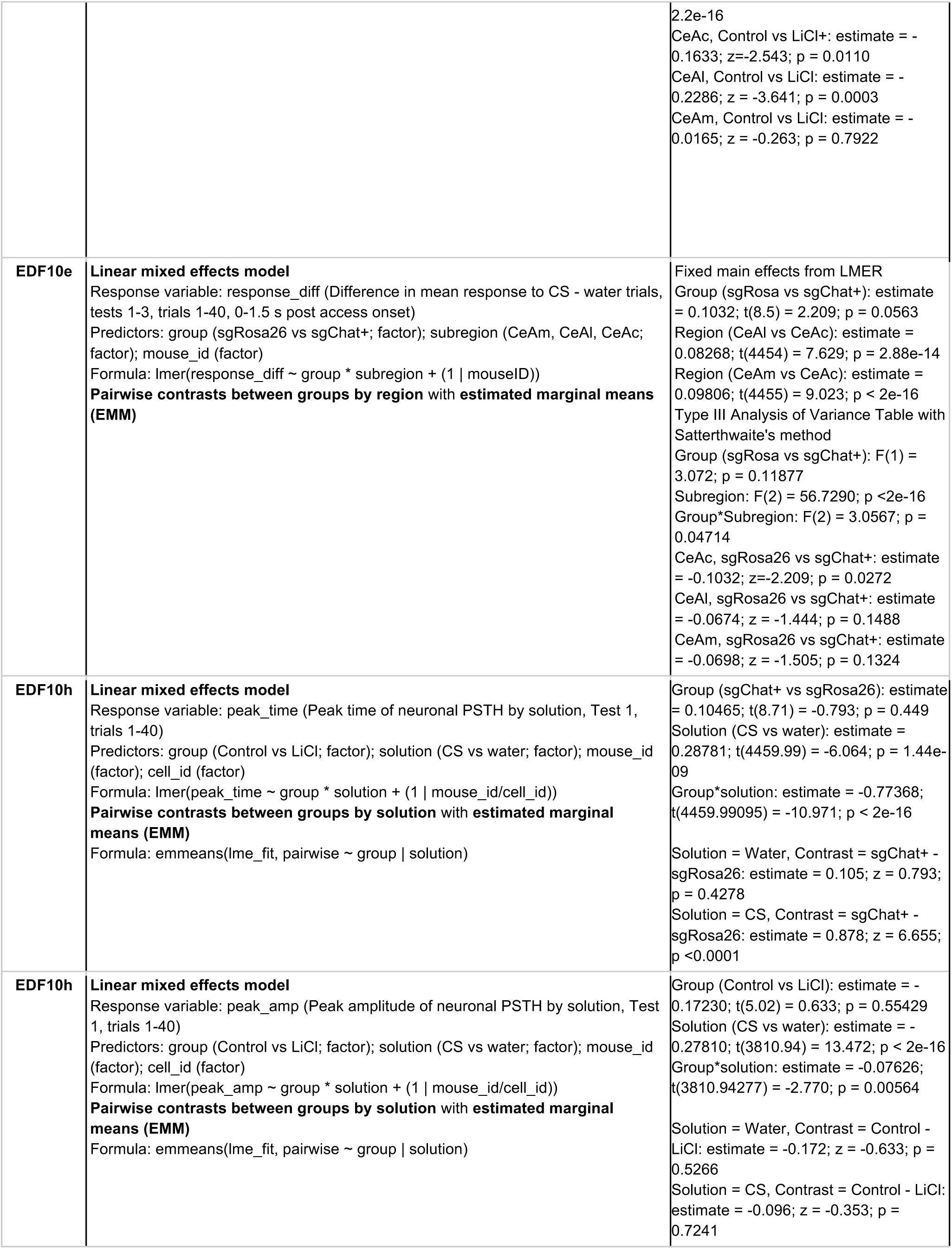
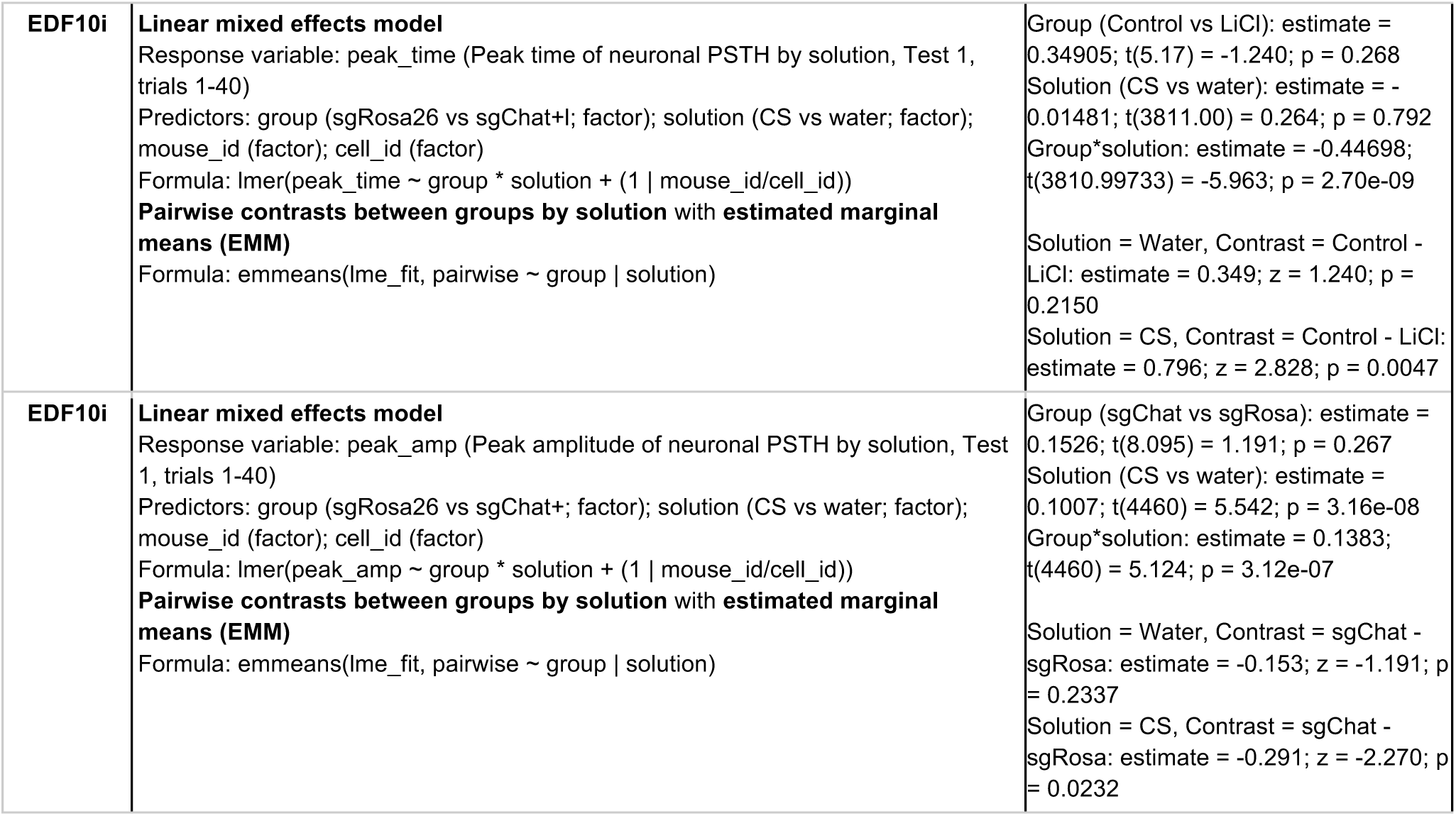

| Figure | Test | Outcome |
| --- | --- | --- |
| **Note:** All statistical tests were two-sided. Paired tests were performed when possible. Corrections for multiple comparisons were performed as noted. Corrected P values are reported. |  |  |
| **Fig. 1** |  |  |
| **1f** | **Two-sample t-test** on CS last-lick time at Test 1 (Control vs CFA) | p=2.3174e-4; Cohen's d=2.8857 |
| **1g** | **Two-sample t-test** on CS licks/total licks at Test 1(Control vs CFA) | p=0.001; Cohen's d=2.4036 |
| **1k** | **Two-sample t-test** on mean fluorescence post injection (Control vs CFA) | p=1.3561e-04; Cohen's d=4.3996 |
| **1l** | **Linear mixed effects model**<br>Response variable mean_fluor (Baseline-subtracted mean fluorescence during access), Conditioning 2<br>Predictors: group (Control vs CFA; factor); solution (CS vs water; factor); mouse_id (factor); trial<br>Formula: lme(mean_fluor ~ group * solution * trial, random=~1\|mouse_id) | Group, p = 0.031<br>Group * solution, p = 0.039 |
| **1m** | **Two-sample t-test** on CS last-lick time at Test 1 (Control vs CFA) | p=5.297e-6; Cohen's d=6.8745 |
| **1m** | **Two-sample t-test** on CS licks/total licks at Test 1(Control vs CFA) | p=1.510e-8; Cohen's d=14.6445 |
| **1n** | **Linear mixed effects model**<br>Response variable mean_fluor (Baseline-subtracted mean fluorescence during access), Tests 1-3<br>Predictors: group (Control vs CFA; factor); solution (CS vs water; factor); lick_count (centreed); trial (centreed); test (centreed); mouse_id (factor)<br>Formula: lme(mean_fluor ~ group * solution * lick_count + trial_count + trial + test, random=~1\|mouse_id) | Solution: estimate = 6.37e-1; t(1.44e3) = 11.85; p<2e-16<br>Group: estimate = 3.22e-1; t(8.55) = 1.72; p=0.121<br>Lick: estimate = 3.30e-2; t(1.44e3) = 6.48; p=1.2e-10<br>Trial: estimate = 9.15e-3; t(1.44e3) = 12.11; p<2e-16<br>Test: estimate = 3.56e-2; t(1.44e3) = 1.73; p=0.085<br>Solution*Group: estimate = -1.82e-1; t(1.44e3) = -2.33; p=0.02<br>Solution*Lick: estimate = -2.19e-2; t(1.44e3) = -3.09; p=0.002<br>Group*Lick: estimate = -1.02e-2; t(1.44e3) = -1.55; p=0.121<br>Solution*Group*Lick: estimate = 4.05e-2; t(1.44e3) = 4.06; p=5.2e-5 |
| **1o** | **Pairwise contrasts** between groups by solution with **estimated marginal means (EMM) on licks from LME (above)** | CFA vs control, Water, estimate = 0.01017; t(1444) = 1.55; p = 0.4060<br>CFA vs control, CS, estimate = -0.03032; t(1445) = -3.93; p = 0.0010 |
| **Fig. 3** |  |  |
| **3e** | **Linear mixed effects model**<br>Response variable fluor_diff (Difference in mean fluorescence (Cch - baseline, z DF/F))<br>Predictors: group (antagonist; factor); cell_id (factor); recording_id<br>Formula: lme(fluor_diff ~ group_id, random=~1\|recording_id/cell_id)<br>**Pairwise contrasts with estimated marginal means (EMM) on LME**<br>ACSF as reference; Holm's correction for multiple comparisons | ACSF -<br>NBQX+AP5+Mecamylamine, estimate = -0.361; t(30) = -1.233; p = 0.249<br>Scopolamine: estimate = -1.048; t(30) = -3.579; p = 0.004<br>Telenzepine: estimate = -0.494; t(30) = -1.581; p = 0.249<br>4DAMP: estimate = -1.087; t(30) = -4.288; p < 0.001 |
|  |  | VU 6019650: estimate = -1.096; t(30) = -3.950; p = 0.002 |
| 3h | **Linear mixed effects model**<br>Response variable fluor_diff (Difference in mean fluorescence (First 60 s post puff - baseline, z DF/F))<br>Predictors: group (antagonist; factor); cell_id (factor); recording_id<br>Formula: lme(fluor_diff ~ group_id, random=~1\|recording_id/cell_id)<br>**Pairwise contrasts with estimated marginal means (EMM) on LME**<br>ACSF as reference; Holm's correction for multiple comparisons | ACSF -<br>NBQX+AP5+Mecamylamine, estimate = -0.793; t(7) = -1.502; p = 0.1768<br>Scopolamine: estimate = -1.911; t(7) = -3.504; p = 0.0199 |
| 3m | **Linear mixed effects model**<br>Response variable fluor_diff (Fluorescence response (First 60 s post AMPA puff, z DF/F))<br>Predictors: group (antagonist; factor); cell_id (factor); response_id (pre, wash, or post; factor); recording_id (factor)<br>Formula: lme(fluor_diff ~ response_id * group_id, random=~1\|recording_id/cell_id)<br>**Pairwise contrasts between groups at each response_id, with estimated marginal means (EMM) on LME**<br>Holm's correction for multiple comparisons | ACSF vs. Cch,<br>Baseline: estimate = 0.114; t(17) = 0.276; p = 0.7862<br>During wash: estimate = -0.740; t(17) = -1.789; p = 0.0915<br>Post wash: estimate = -1.348; t(17) = -3.260; p = 0.0046 |
| 4c | One-sample t-test on mean fluorescence post-LiCl injection | p=0.03; Cohen's d = 2.6 |
| 4f | Two-sample t-test on oEPSC1 amplitude | p=0.4358; Cohen's d = 0.3252 |
| 4f | Two-sample t-test on oEPSC2 amplitude | p=0.3178; Cohen's d = 0.4188 |
| 4f | Two-sample t-test on Paired pulse ratio (oEPSC2 / oEPSC1) | p=0.3375; Cohen's d=0.4017 |
| 4i | Two-sample t-test on Cumulative CS licks on Conditioning 1 | p=0.1849; Cohen's d=0.6972 |
| 4j | Two-sample t-test on CS last-lick time at Tests 1-3 | p=0.0599; Cohen's d=1.0235 |
| 4j | **Linear mixed effects model**<br>Response variable cs_pref (cs_licks / total_licks)<br>Predictors: group (sgRosa26 vs sgChat; factor); test (1-3); mouse_id (factor)<br>Formula: lme(cs_pref ~ group*test, random=~1\|mouse_id) | Group: estimate = 0.07368; t(43.58) = 2.49; p=0.017<br>Test: estimate = 0.01809; t(30) = 2.17; p=0.038<br>Group*Test: estimate = -0.01048; t(30) = -0.89; p = 0.382 |
| 4m | Two-sample t-test on Cumulative CS licks on Conditioning 1 | p=0.1925; Cohen's d=0.7458 |
| 4n | Two-sample t-test on CS last-lick time at Test | p=1.9843e-5; Cohen's d=3.6566 |
| 4n | Two-sample t-test on CS preference at Test | p=4.96e-4; Cohen's d=2.5495 |
| 5l | **Linear mixed effects model**<br>Response variable mean_fluor (Mean fluorescence value during post-injection period)<br>Predictors: group (Control vs LiCl; factor); is_CS (binary denoting CS-encoding during Conditioning 1; factor); cell_id (factor)<br>Formula: lme(mean_fluor ~ group*is_cs, random=~1\|cell_id) | Group: estimate = 0.2404; t(3809) = 6.76; p<1e-5<br>is_CS: estimate = 0.0425; t(3809) = 0.92; p=0.3579<br>Group*is_CS: estimate = 0.0619; t(3809) = 1.01; p=0.3133 |
|  | **Pairwise contrast between groups with estimated marginal means (EMM) on LME** | Control vs LiCl, estimate=-0.271, t(3809) = -8.842; p<1e-5 |
|  | **Pairwise contrasts between groups by is_CS, with estimated marginal means (EMM) on LME** | is_CS = 0, estimate=-0.241; t(3809) = -6.759; p<1e-5<br>is_CS = 1, estimate=-0.302; t(3809) = -6.044; p<1e-5 |
|  | **Pairwise contrasts between is_CS by group, with estimated marginal means (EMM) on LME** | Control, estimate=-0.0425; t(3809) = -0.919; p=0.3579<br>LiCl, estimate=-0.1044; t(3809) = -2.586; p=0.0097 |
| **5n** | **Linear mixed effects model**<br>Response variable mean_fluor (Mean fluorescence value during post-injection period)<br>Predictors: group (Control vs LiCl; factor); is_CS (binary denoting CS-encoding during Conditioning 1; factor); cell_id (factor)<br>Formula: lme(mean_fluor ~ group*is_cs, random=~1\|cell_id) | Group: estimate = 0.1042; t(4242) = 2.12; p=0.0337<br>is_CS: estimate = 0.2161; t(4242) = 3.68; p=0.0002<br>Group*is_CS: estimate = -0.1747; t(4242) = -2.24; p=0.0250 |
|  | **Pairwise contrast between groups with estimated marginal means (EMM) on LME** | sgRosa26 vs sgChat+, estimate=-0.0168, t(4242) = -0.432; p=0.6659 |
|  | **Pairwise contrasts between groups by is_CS, with estimated marginal means (EMM) on LME** | is_CS = 0, estimate=-0.1042; t(4242) = -2.124; p=0.0337<br>is_CS = 1, estimate=0.0705; t(4242) = 1.165; p=0.2439 |
|  | **Pairwise contrasts between is_CS by group, with estimated marginal means (EMM) on LME** | sgRosa26, estimate=-0.2161; t(4242) = -3.685; p=0.0002<br>sgChat+, estimate=-0.0414; t(4242) = -0.808; p=0.4193 |
| **Fig. 6** |  |  |
| **6a** | **Two-sample t-test** on CS last-lick time at Tests 1-3, Control vs LiCl | p=0.0086; Cohen's d=3.1987 |
| **6a** | **Two-sample t-test** on CS preference at Tests 1-3, Control vs LiCl | p=0.0177; Cohen's d=2.6558 |
| **6b** | **Two-sample t-test** on CS last-lick time at Tests 1-3, sgRosa26 vs sgChat | p=0.136; Cohen's d=1.9937 |
| **6b** | **Two-sample t-test** on CS preference at Tests 1-3, sgRosa26 vs sgChat | p=0.0049; Cohen's d=2.4330 |
| **6c** | **Linear mixed effects model**<br>Response variable: peak_time (Peak time of neuronal PSTH by solution, Test 1, trials 1-40)<br>Predictors: group (Control vs LiCl; factor); solution (CS vs water; factor); mouse_id (factor); cell_id (factor)<br>Formula: lmer(peak_time ~ group * solution + (1 \| mouse_id/cell_id))<br>**Pairwise contrasts between groups by solution with estimated marginal means (EMM)**<br>Formula: emmeans(lme_fit, pairwise ~ group \| solution) | Group (Control vs LiCl): estimate = -0.27996; t(5.19) = -1.037; p = 0.345631<br>Solution (CS vs water): estimate = -0.35106; t(3811.00) = -6.166; p = 7.75e-10<br>Group*solution: estimate = -0.28142; t(3810.99715) = -3.705; p = 0.000214<br>Solution = Water, Contrast = Control - LiCl: estimate = 0.280; z = 1.037; p = 0.2997<br>Solution = CS, Contrast = Control - LiCl: estimate = 0.561; z = 2.079; p = 0.0376 |
| **6g** | **Linear mixed effects model**<br>Response variable: peak_time (Peak time of neuronal PSTH by solution, Test 1, trials 1-40)<br>Predictors: group (sgRosa26 vs sgChat+I; factor); solution (CS vs water; factor); mouse_id (factor); cell_id (factor)<br>Formula: lmer(peak_time ~ group * solution + (1 \| mouse_id/cell_id))<br>**Pairwise contrasts between groups by solution with estimated marginal means (EMM)**<br>Formula: emmeans(lme_fit, pairwise ~ group \| solution) | Group (sgChat vs sgRosa): estimate = -0.08632; t(9.11) = -0.597; p = 0.564910<br>Solution (CS vs water): estimate = -0.17009; t(4459.99) = -3.302; p = 0.000968<br>Group*solution: estimate = -0.53926; t(4459.99473) = -7.045; p = 2.14e-12<br>Solution = Water, Contrast = sgChat - sgRosa: estimate = 0.0863; z = 0.597; p = 0.5504<br>Solution = CS, Contrast = sgChat - sgRosa: estimate = 0.6256; z = 4.328; p <0.0001 |
| 6e | **Linear mixed effects model**<br>Response variable: peak_time_change (Difference in peak time by solution from Cond 1 to Test 1, trials 1-40)<br>Predictors: group (Control vs LiCl; factor); solution (CS vs water; factor); mouse_id (factor); cell_id (factor)<br>Formula: lmer(peak_time_change ~ group * solution + (1 \| mouse_id/cell_id))<br>**Pairwise contrasts between groups by solution with estimated marginal means (EMM)**<br>Formula: emmeans(lme_fit, ~ group * solution) | Group (Control vs LiCl): estimate = 0.65171; t(8.41) = -7.047; p = 8.40e-05<br>Solution (Water vs CS): estimate = 0.14751; t(4603.67) = -2.186; p = 0.0288<br>Group*Solution: estimate = 0.58503; t(4635.76244) = 6.424; p = 1.46e-10<br>Group=Control, Contrast = CS - Water: estimate = 0.148; z = 2.186; p = 0.0288<br>Group=LiCl, Contrast = CS - Water: estimate = -0.438; z = -7.151; p <0.0001<br>Group*Solution interaction contrast: estimate = 0.585; z = 6.424; p<0.0001 |
| 6f | **Linear mixed effects model**<br>Response variable: slope_change (Difference in slope by solution from Cond 1 to Test 1, trials 1-40)<br>Predictors: group (Control vs LiCl; factor); solution (CS vs water; factor); mouse_id (factor); cell_id (factor)<br>Formula: lmer(slope_change ~ group * solution + (1 \| mouse_id/cell_id))<br>**Pairwise contrasts between groups by solution with estimated marginal means (EMM)**<br>Formula: emmeans(lme_fit, ~ group * solution) | Group (Control vs LiCl): estimate = -0.19164; t(5.03) = 1.747; p = 0.14075<br>Solution (Water vs CS): estimate = 0.03072; t(4935.30) = -3.042; p = 0.00236<br>Group*Solution: estimate = -0.10065; t(4963.30791) = -7.376; p = 1.9e-13<br>Group=Control, Contrast = CS - Water: estimate = 0.0307; z = 3.042; p = 0.0023<br>Group=LiCl, Contrast = CS - Water: estimate = 0.1314; z = 14.319; p <0.0001<br>Group*Solution interaction contrast: estimate = -0.101; z = -7.376; p<0.0001 |
| 6i | **Linear mixed effects model**<br>Response variable: peak_time_change (Difference in peak time by solution from Cond 1 to Test 1, trials 1-40)<br>Predictors: group (sgChat+ vs sgRosa26; factor); solution (CS vs water; factor); mouse_id (factor); cell_id (factor)<br>Formula: lmer(peak_time_change ~ group * solution + (1 \| mouse_id/cell_id))<br>**Pairwise contrasts between groups by solution with estimated marginal means (EMM)**<br>Formula: emmeans(lme_fit, ~ group * solution) | Group (sgChat+ vs sgRosa): estimate = 0.41481; t(8.70) = -2.199; p = 0.05641<br>Solution (Water vs CS): estimate = -0.59793; t(5394.62) = 10.186; p < 2e-16<br>Group*Solution: estimate = 0.36685; t(5314.18556) = 4.215; p = 2.54e-05<br>Group=sgChat+, Contrast = CS - Water: estimate = -0.598; z = -10.186; p <0.0001<br>Group=sgRosa, Contrast = CS - Water: estimate = -0.965; z = -15.016; p <0.0001<br>Group*Solution interaction contrast: estimate = 0.367; z = 4.215; p<0.0001 |
| 6j | **Linear mixed effects model**<br>Response variable: slope_change (Difference in slope by solution from Cond 1 to Test 1, trials 1-40)<br>Predictors: group (sgChat+ vs sgRosa26; factor); solution (CS vs water; factor); mouse_id (factor); cell_id (factor)<br>Formula: lmer(slope_change ~ group * solution + (1 \| mouse_id/cell_id))<br>**Pairwise contrasts between groups by solution with estimated marginal means (EMM)**<br>Formula: emmeans(lme_fit, ~ group * solution) | Group (sgChat vs sgRosa): estimate = -0.1372; t(8.142) = 2.804; p = 0.0227<br>Solution (Water vs CS): estimate = -0.02545; t(5980) = 3.140; p = 0.0017<br>Group*Solution: estimate = -0.1014; t(5900) = -8.439; p < 2e-16<br>Group=sgChat, Contrast = CS - Water: estimate = -0.0254; z = -3.140; p = 0.0017<br>Group=sgRosa, Contrast = CS - Water: estimate = 0.0760; z = 8.562; p <0.0001<br>Group*Solution interaction contrast: estimate = -0.101; z = -8.439; p<0.0001 |
| 6k, top | **Linear mixed effects model**<br>Response variable last_lick_time (Time of last-lick on lick trial (s), Tests 1-3, Trials 1-40)<br>Predictors: group (Control vs LiCl; factor); peak_slope_z (slope-to-peak of trial CeA population PSTH, z-scored within mouse; factor); solution (CS vs water; factor); mouse_id (factor)<br>Formula: lme(last_lick_time ~ group * peak_slope_z * solution, random=~1\|mouseID)<br>**Estimated marginal means (EMM) of linear trends of peak_slope_z on last_lick_time by group for CS**<br>Formula: emtrends(lme_fit ~ group, var = 'peak_slope_z', at = list(solution_id = 'CS')) | Group (Base vs LiCl): estimate = -0.122; t(5) = -0.67; p = 0.5323<br>peak_slope_z: estimate = 0.0696; t(721) = 1.03; p = 0.3032<br>solution: estimate = -1.0022; t(721) = -12.403; p < 1e-4<br>Group*peak_slope_z: estimate = 0.3276; t(721) = 3.628; p = 0.0003<br>Group*solution: estimate = 1.1401; t(721) = 10.396; p < 1e-4<br>peak_slope_z*solution: estimate = -0.2441; t(721) = -2.890; p = 0.0040<br>Group*peak_slope_z*solution: estimate = -0.1238; t(721) = -1.086; p = 0.2780<br>Group = LiCl, slope of last_lick_time on peak_slope_z (at CS): estimate = -0.1745; t(721) = -3.495; p = 0.0005<br>Group = Sal, slope of last_lick_time on peak_slope_z (at CS): estimate = 0.0293; t(721) = 0.616; p = 0.5381<br>Group (Sal vs LiCl): estimate = -0.204; t(721) = -2.955; p = 0.0032 |
| 6k, bottom | **Linear mixed effects model**<br>Response variable last_lick_time (Time of last-lick on lick trial (s), Tests 1-3, Trials 1-40)<br>Predictors: group (sgRosa26 vs sgChat+; factor); peak_slope_z (slope-to-peak of trial CeA population PSTH, z-scored within mouse; factor); solution (CS vs water; factor); mouse_id (factor)<br>Formula: lme(last_lick_time ~ group * peak_slope_z * solution, random=~1\|mouseID)<br>**Estimated marginal means (EMM) of linear trends of peak_slope_z on last_lick_time by group for CS**<br>Formula: emtrends(lme_fit ~ group, var = 'peak_slope_z', at = list(solution_id = 'CS')) | Group (sgRosa26 vs sgChat+): estimate = -0.0213; t(8) = -0.283; p = 0.7840<br>peak_slope_z: estimate = 0.1048; t(1020) = 1.944; p = 0.0521<br>solution: estimate = 0.0299; t(1020) = 0.496; p = 0.6197<br>Group*peak_slope_z: estimate = -0.1638; t(1020) = -2.010; p = 0.0447<br>Group*solution: estimate = -0.3617; t(1020) = -4.188; p < 1e-4<br>peak_slope_z*solution: estimate = -0.1503; t(1020) = -2.311; p = 0.0210<br>Group*peak_slope_z*solution: estimate = 0.0298; t(1020) = 0.312; p = 0.7549<br>Group = sgChat+, slope of last_lick_time on peak_slope_z (at CS): estimate = -0.0455; t(1020) = -1.276; p = 0.2023<br>Group = sgRosa26, slope of last_lick_time on peak_slope_z (at CS): estimate = -0.1794; t(1020) = -5.293; p < 0.0001<br>Group (sgRosa26 vs sgChat+): estimate = 0.134; t(1020) = 2.722; p = 0.0066 |
| **6n** | **Linear mixed effects model**<br>Response variable: response_diff (Difference in mean response to CS - water trials, tests 1-3, trials 1-10, 0-1.5 s post access onset)<br>Predictors: group (Control vs LiCl; factor); subregion (CeAm, CeAl, CeAc; factor); mouse_id (factor)<br>Formula: lmer(response_diff ~ group * subregion + (1 \| mouseID))<br>**Pairwise contrasts between groups by region with estimated marginal means (EMM)** | Fixed main effects from LMER<br>Group (Control vs LiCl): estimate = 0.1957; t(5.809) = 2.621; p = 0.0407<br>Region (CEAl vs CEAc): estimate = -0.01010; t(3804) = -0.521; p = 0.6027<br>Region (CEAm vs CEAc): estimate = 0.1419; t(3806) = 6.785; p = 1.34e-11<br>Type III Analysis of Variance Table with Satterthwaite's method<br>Group (Control vs LiCl): F(1) = 4.7432; p = 0.08168<br>Region: F(2) = 15.1634; p = 2.759e-07<br>Group*Region: F(2) = 43.1931; p = < 2.2e-16<br>CEAc, Control vs LiCl: estimate = -0.1957; z = -2.621; p = 0.0088<br>CEAl, Control vs LiCl: estimate = -0.2359; z = -3.246; p = 0.0012<br>CEAm, Control vs LiCl: estimate = -0.0375; z = -0.517; p = 0.6055 |
| **6p** | **Linear mixed effects model**<br>Response variable: response_diff (Difference in mean response to CS - water trials, tests 1-3, trials 1-10, 0-1.5 s post access onset)<br>Predictors: group (sgRosa26 vs sgChat+; factor); subregion (CeAm, CeAl, CeAc; factor); mouse_id (factor)<br>Formula: lmer(response_diff ~ group * subregion + (1 \| mouseID))<br>Pairwise contrasts between groups by region with estimated marginal means (EMM) | Fixed main effects from LMER<br>Group (sgChat vs sgRosa): estimate = 0.15463; t(8.95203) = 3.334; p = 0.00881<br>Region (CEAl vs CEAc): estimate = 0.08686; t(4455.91) = 6.224; p = 5.29e-10<br>Region (CEAm vs CEAc): estimate = 0.08519; t(4440.57) = 6.154; p = 8.24e-10<br>Type III Analysis of Variance Table with Satterthwaite's method<br>Group (sgChat vs sgRosa): F(1) = 7.8439; p = 0.02402<br>Region: F(2) = 26.6357; p = 3.169e-12<br>Group*Region: F(2) = 4.1907; p = 0.01520<br>CEAc, sgChat vs sgRosa: estimate = -0.1546; z = -3.334; p = 0.0009<br>CEAl, sgChat vs sgRosa: estimate = -0.0946; z = -2.044; p = 0.0410<br>CEAm, sgChat vs sgRosa: estimate = -0.1262; z = -2.754; p = 0.0059 |
| **Ext. Data. Fig. 1** |  |  |
| **EDF1c** | **Two-sample t-test** on CS last-lick time at Test, 21d (Control vs CFA) | p=0.0048; Cohen's d=1.926 |
| **EDF1c** | **Two-sample t-test** on CS licks/total licks at Test. 21d (Control vs CFA) | p=0.0211; Cohen's d=1.479 |
| **Ext. Data. Fig. 4** |  |  |
| **EDF4e** | **Paired sample t-test** on I_hold | p=0.0338 |
| **EDF4f** | **Paired sample t-test** on delta_I_deactivation | p=0.0114 |
| **EDF4g** | **Paired sample t-test** on delta_I_reactivation | p=0.024 |
| **EDF4h** | **Paired sample t-test** on tau_deactivation | p=0.0463 |
| **EDF4i** | **Paired sample t-test** on tau_reactivation | p=0.008 |
| **Ext. Data. Fig. 5** |  |  |
| **EDF5e** | **Linear mixed effects model**<br>Response variable: fluor_diff (Difference in mean fluorescence (Cch - baseline, z DF/F))<br>Predictors: group (antagonist; factor); cell_id (factor); recording_id (factor)<br>Formula: lme(fluor_diff ~ group_id, random=~1\|recording_id/cell_id)<br>**Pairwise contrasts with estimated marginal means (EMM) on LME**<br>TTX as reference. | TTX -<br>TTX+Scopolamine, estimate = -0.249;<br>t(11) = -2.223; p = 0.0481 |
| **Ext. Data. Fig. 6** |  |  |
| **EDF6g** | **Two-sample t-test** on mean speed during baseline period (sgRosa26 vs sgChat+) | p=0.3608 |
| **EDF6g** | **Two-sample t-test** on mean speed during baseline period (sgRosa26 vs sgChat+) | p=2.02e-4 |
| **Ext. Data. Fig. 9** |  |  |
| **EDF9a** | **Linear mixed effects model**<br>Response variable: peak_amp (Peak amplitude of neuronal PSTH by solution, Test 1, trials 1-40)<br>Predictors: group (Control vs LiCl; factor); solution (CS vs water; factor); mouse_id (factor); cell_id (factor)<br>Formula: lmer(peak_amp ~ group * solution + (1 \| mouse_id/cell_id))<br>**Pairwise contrasts between groups by solution with estimated marginal means (EMM)**<br>Formula: emmeans(lme_fit, pairwise ~ group \| solution) | Group (Control vs LiCl): estimate = 0.24766; t(5.02) = 0.727; p = 0.4998<br>Solution (CS vs water): estimate = 0.16843; t(3811.00) = 7.126; p = 1.23e-12<br>Group*solution: estimate = -0.01144; t(3810.99971) = -0.363; p = 0.7168<br>Solution = Water, Contrast = Control - LiCl: estimate = -0.248; z = -0.727; p = 0.4674<br>Solution = CS, Contrast = Control - LiCl: estimate = -0.236; z = -0.693; p = 0.4882 |
| **EDF9b** | **Linear mixed effects model**<br>Response variable: peak_amp (Peak amplitude of neuronal PSTH by solution, Test 1, trials 1-40)<br>Predictors: group (sgRosa26 vs sgChat+; factor); solution (CS vs water; factor); mouse_id (factor); cell_id (factor)<br>Formula: lmer(peak_amp ~ group * solution + (1 \| mouse_id/cell_id))<br>**Pairwise contrasts between groups by solution with estimated marginal means (EMM)**<br>Formula: emmeans(lme_fit, pairwise ~ group \| solution) | Group (sgChat+ vs sgRosa26): estimate = -0.17804; t(8.22) = 2.146; p = 0.0632<br>Solution (CS vs water): estimate = -0.18975; t(4460.00) = 12.505; p < 2e-16<br>Group*solution: estimate = -0.15836; t(4460.00006) = -7.023; p = 2.50e-12<br>Solution = Water, Contrast = sgChat+ - sgRosa26: estimate = -0.1780; z = -2.146; p = 0.0318<br>Solution = CS, Contrast = sgChat+ - sgRosa26: estimate = -0.0197; z = -0.237; p = 0.8125 |
| **Ext. Data. Fig. 10** |  |  |
| **EDF10c** | **Linear mixed effects model**<br>Response variable: response_diff (Difference in mean response to CS - water trials, tests 1-3, trials 1-40, 0-1.5 s post access onset)<br>Predictors: group (Control vs LiCl; factor); subregion (CeAm, CeAl, CeAc; factor); mouse_id (factor)<br>Formula: lmer(response_diff ~ group * subregion + (1 \| mouseID))<br>**Pairwise contrasts between groups by region with estimated marginal means (EMM)** | Fixed main effects from LMER<br>Group (Control vs LiCl): estimate = 0.1633; t(5.68889) = 2.543; p = 0.04603<br>Region (CeAl vs CeAc): estimate = -0.03215; t(3803.46) = -2.098; p = 0.03598<br>Region (CeAm vs CeAc): estimate = 0.12688; t(3806.69) = 7.68; p = 2.01e-14<br>Type III Analysis of Variance Table with Satterthwaite's method<br>Group (Control vs LiCl): F(1) = 4.7985; p = 0.08015<br>Subregion: F(2) = 21.7711; p = 3.9e-10<br>Group*Subregion: F(2) = 76.9174; p = < |
|  |  | 2.2e-16<br>CeAc, Control vs LiCl+: estimate = -0.1633; z=-2.543; p = 0.0110<br>CeAl, Control vs LiCl: estimate = -0.2286; z = -3.641; p = 0.0003<br>CeAm, Control vs LiCl: estimate = -0.0165; z = -0.263; p = 0.7922 |
| **EDF10e** | **Linear mixed effects model**<br>Response variable: response_diff (Difference in mean response to CS - water trials, tests 1-3, trials 1-40, 0-1.5 s post access onset)<br>Predictors: group (sgRosa26 vs sgChat+; factor); subregion (CeAm, CeAl, CeAc; factor); mouse_id (factor)<br>Formula: lmer(response_diff ~ group * subregion + (1 \| mouseID))<br>**Pairwise contrasts between groups by region with estimated marginal means (EMM)** | Fixed main effects from LMER<br>Group (sgRosa vs sgChat+): estimate = 0.1032; t(8.5) = 2.209; p = 0.0563<br>Region (CeAl vs CeAc): estimate = 0.08268; t(4454) = 7.629; p = 2.88e-14<br>Region (CeAm vs CeAc): estimate = 0.09806; t(4455) = 9.023; p < 2e-16<br>Type III Analysis of Variance Table with Satterthwaite's method<br>Group (sgRosa vs sgChat+): F(1) = 3.072; p = 0.11877<br>Subregion: F(2) = 56.7290; p < 2e-16<br>Group*Subregion: F(2) = 3.0567; p = 0.04714<br>CeAc, sgRosa26 vs sgChat+: estimate = -0.1032; z=-2.209; p = 0.0272<br>CeAl, sgRosa26 vs sgChat+: estimate = -0.0674; z = -1.444; p = 0.1488<br>CeAm, sgRosa26 vs sgChat+: estimate = -0.0698; z = -1.505; p = 0.1324 |
| **EDF10h** | **Linear mixed effects model**<br>Response variable: peak_time (Peak time of neuronal PSTH by solution, Test 1, trials 1-40)<br>Predictors: group (Control vs LiCl; factor); solution (CS vs water; factor); mouse_id (factor); cell_id (factor)<br>Formula: lmer(peak_time ~ group * solution + (1 \| mouse_id/cell_id))<br>**Pairwise contrasts between groups by solution with estimated marginal means (EMM)**<br>Formula: emmeans(lme_fit, pairwise ~ group \| solution) | Group (sgChat+ vs sgRosa26): estimate = 0.10465; t(8.71) = -0.793; p = 0.449<br>Solution (CS vs water): estimate = 0.28781; t(4459.99) = -6.064; p = 1.44e-09<br>Group*solution: estimate = -0.77368; t(4459.99095) = -10.971; p < 2e-16<br>Solution = Water, Contrast = sgChat+ - sgRosa26: estimate = 0.105; z = 0.793; p = 0.4278<br>Solution = CS, Contrast = sgChat+ - sgRosa26: estimate = 0.878; z = 6.655; p < 0.0001 |
| **EDF10h** | **Linear mixed effects model**<br>Response variable: peak_amp (Peak amplitude of neuronal PSTH by solution, Test 1, trials 1-40)<br>Predictors: group (Control vs LiCl; factor); solution (CS vs water; factor); mouse_id (factor); cell_id (factor)<br>Formula: lmer(peak_amp ~ group * solution + (1 \| mouse_id/cell_id))<br>**Pairwise contrasts between groups by solution with estimated marginal means (EMM)**<br>Formula: emmeans(lme_fit, pairwise ~ group \| solution) | Group (Control vs LiCl): estimate = -0.17230; t(5.02) = 0.633; p = 0.55429<br>Solution (CS vs water): estimate = -0.27810; t(3810.94) = 13.472; p < 2e-16<br>Group*solution: estimate = -0.07626; t(3810.94277) = -2.770; p = 0.00564<br>Solution = Water, Contrast = Control - LiCl: estimate = -0.172; z = -0.633; p = 0.5266<br>Solution = CS, Contrast = Control - LiCl: estimate = -0.096; z = -0.353; p = 0.7241 |
| **EDF10i** | **Linear mixed effects model**<br>Response variable: peak_time (Peak time of neuronal PSTH by solution, Test 1, trials 1-40)<br>Predictors: group (sgRosa26 vs sgChat+; factor); solution (CS vs water; factor); mouse_id (factor); cell_id (factor)<br>Formula: lmer(peak_time ~ group * solution + (1 \| mouse_id/cell_id))<br>**Pairwise contrasts between groups by solution with estimated marginal means (EMM)**<br>Formula: emmeans(lme_fit, pairwise ~ group \| solution) | Group (Control vs LiCl): estimate = 0.34905; t(5.17) = -1.240; p = 0.268<br>Solution (CS vs water): estimate = -0.01481; t(3811.00) = 0.264; p = 0.792<br>Group*solution: estimate = -0.44698; t(3810.99733) = -5.963; p = 2.70e-09<br>Solution = Water, Contrast = Control - LiCl: estimate = 0.349; z = 1.240; p = 0.2150<br>Solution = CS, Contrast = Control - LiCl: estimate = 0.796; z = 2.828; p = 0.0047 |
| **EDF10i** | **Linear mixed effects model**<br>Response variable: peak_amp (Peak amplitude of neuronal PSTH by solution, Test 1, trials 1-40)<br>Predictors: group (sgRosa26 vs sgChat+; factor); solution (CS vs water; factor); mouse_id (factor); cell_id (factor)<br>Formula: lmer(peak_amp ~ group * solution + (1 \| mouse_id/cell_id))<br>**Pairwise contrasts between groups by solution with estimated marginal means (EMM)**<br>Formula: emmeans(lme_fit, pairwise ~ group \| solution) | Group (sgChat vs sgRosa): estimate = 0.1526; t(8.095) = 1.191; p = 0.267<br>Solution (CS vs water): estimate = 0.1007; t(4460) = 5.542; p = 3.16e-08<br>Group*solution: estimate = 0.1383; t(4460) = 5.124; p = 3.12e-07<br>Solution = Water, Contrast = sgChat - sgRosa: estimate = -0.153; z = -1.191; p = 0.2337<br>Solution = CS, Contrast = sgChat - sgRosa: estimate = -0.291; z = -2.270; p = 0.0232 |

## References

1. Garcia, J., Kimeldorf, D. J. & Koelling, R. A. Conditioned aversion to saccharin resulting from exposure to gamma radiation. Science 122, 157–158 (1955).

2. Garcia, J. & Koelling, R. A. Relation of cue to consequence in avoidance learning. Psychon Sci 4, 123–124 (1966).

3. Garcia, J., McGowan, B. K., Ervin, F. R. & Koelling, R. A. Cues: their relative effectiveness as a function of the reinforcer. Science 160, 794–795 (1968).

4. Garcia, J., Hankins, W. G. & Rusiniak, K. W. Behavioral regulation of the milieu interne in man and rat. Science 185, 824–831 (1974).

5. Adams, C. D. & Dickinson, A. Instrumental Responding following Reinforcer Devaluation. Q J Exp Psychol B 33, 109–121 (1981).

6. Wilcoxon, H. C., Dragoin, W. B. & Kral, P. A. Illness-induced aversions in rat and quail: relative salience of visual and gustatory cues. Science 171, 826–828 (1971).

7. Gustavson, C. R., Garcia, J., Hankins, W. G. & Rusiniak, K. W. Coyote predation control by aversive conditioning. Science 184, 581–583 (1974).

8. Garb, J. L. & Stunkard, A. J. Taste aversions in man. Am J Psychiatry 131, 1204–1207 (1974).

9. Bernstein, I. L. Learned taste aversions in children receiving chemotherapy. Science 200, 1302–1303 (1978).

10. Bernstein, I. L. & Webster, M. M. Learned taste aversions in humans. Physiol Behav 25, 363– 366 (1980).

11. Boakes, R. A. & Costa, D. S. J. Temporal contiguity in associative learning: Interference and decay from an historical perspective. J Exp Psychol Anim Learn Cogn 40, 381–400 (2014).

12. Seol, G. H. et al. Neuromodulators Control the Polarity of Spike-Timing-Dependent Synaptic Plasticity. Neuron 55, 919–929 (2007).

13. Zimmerman, C. A. et al. A neural mechanism for learning from delayed postingestive feedback. Nature 642, 700–709 (2025).

14. Schiff, H. C. et al. An Insula–Central Amygdala Circuit for Guiding Tastant-Reinforced Choice Behavior. J. Neurosci. 38, 1418–1429 (2018).

15. Wang, L. et al. The coding of valence and identity in the mammalian taste system. Nature 558, 127–131 (2018).

16. Ponserre, M., Peters, C., Fermani, F., Conzelmann, K.-K. & Klein, R. The insula cortex contacts distinct output streams of the central amygdala. J Neurosci 40, 8870–8882 (2020).

17. Kim, J., Zhang, X., Muralidhar, S., LeBlanc, S. A. & Tonegawa, S. Basolateral to central amygdala neural circuits for appetitive behaviors. Neuron 93, 1464–1479 5 (2017).

18. Li, B. Central amygdala cells for learning and expressing aversive emotional memories. Curr Opin Behav Sci 26, 40–45 (2019).

19. Yang, T. et al. Plastic and stimulus-specific coding of salient events in the central amygdala. Nature 616, 510–519 (2023).

20. Janak, P. H. & Tye, K. M. From circuits to behaviour in the amygdala. Nature 517, 284–292 (2015).

21. Douglass, A. M. et al. Central amygdala circuits modulate food consumption through a positive-valence mechanism. Nat. Neurosci. 20, 1384–1394 (2017).

22. Cai, H., Haubensak, W., Anthony, T. E. & Anderson, D. J. Central amygdala PKC-δ(+) neurons mediate the influence of multiple anorexigenic signals. Nat Neurosci 17, 1240–1248 (2014).

23. Hardaway, J. A. et al. Central Amygdala Prepronociceptin-Expressing Neurons Mediate Palatable Food Consumption and Reward. Neuron 102, 1037–1052.e7 (2019).

24. Ding, W., Weltzien, H., Peters, C. & Klein, R. Nausea-induced suppression of feeding is mediated by central amygdala Dlk1-expressing neurons. Cell Rep 43, 113990 (2024).

25. Ding, W., Almeida, A. P., Wang, W., Qu, C. & Klein, R. Central amygdala Isl1 neurons control biting by integrating sensory and motivational signals. bioRxiv. (2026) doi:10.64898/2026.02.03.703447.

26. Moscarello, J. M. & Penzo, M. A. The central nucleus of the amygdala and the construction of defensive modes across the threat-imminence continuum. Nat Neurosci 25, 999–1008 (2022).

27. Pauli, J. L. et al. Molecular and anatomical characterization of parabrachial neurons and their axonal projections. eLife 11, e81868 (2022).

28. Campos, C. A., Bowen, A. J., Roman, C. W. & Palmiter, R. D. Encoding of danger by parabrachial CGRP neurons. Nature 555, 617–622 (2018).

29. Condon, L. F. et al. Parabrachial Calca neurons drive nociplasticity. Cell Rep. 43, (2024).

30. Chen, J. Y., Campos, C. A., Jarvie, B. C. & Palmiter, R. D. Parabrachial CGRP Neurons Establish and Sustain Aversive Taste Memories. Neuron 100, 891–899 5 (2018).

31. Reilly, S. The parabrachial nucleus and conditioned taste aversion. Brain Res Bull 48, 239–254 (1999).

32. Carter, M. E., Soden, M. E., Zweifel, L. S. & Palmiter, R. D. Genetic identification of a neural circuit that suppresses appetite. Nature 503, 111–114 (2013).

33. Carter, M. E., Han, S. & Palmiter, R. D. Parabrachial calcitonin gene-related peptide neurons mediate conditioned taste aversion. J Neurosci 35, 4582–4586 (2015).

34. Jarvie, B. C. et al. Ingestion-activated CGRP neurons control learning but not satiety. Neuron (2026) 10.1016/j.neuron.2026.08.001.

35. Zajdel, J., Sköld, J., Jaarola, M., Singh, A. K. & Engblom, D. Calcitonin gene related peptide α is dispensable for many danger-related motivational responses. Sci Rep 11, 16204 (2021).

36. Chen, W. et al. Distinct eLPB^ChAT^ projections for methamphetamine withdrawal anxiety and primed reinstatement of conditioned place preference. Theranostics 14, 2881–2896 (2024).

37. Aitta-aho, T. et al. Basal Forebrain and Brainstem Cholinergic Neurons Differentially Impact Amygdala Circuits and Learning-Related Behavior. Curr. Biol. 28, 2557–2569.e4 (2018).

38. Hasselmo, M. E. The role of acetylcholine in learning and memory. Curr Opin Neurobiol 16, 710–715 (2006).

39. Deutsch, R. Effects of atropine on conditioned taste aversion. Pharmacol Biochem Behav 8, 685–694 (1978).

40. Morin, J.-P. et al. Muscarinic receptor signaling in the amygdala is required for conditioned taste aversion. Neurosci. Lett. 740, 135466 (2021).

41. Clark, E. W. & Bernstein, I. L. Boosting cholinergic activity in gustatory cortex enhances the salience of a familiar conditioned stimulus in taste aversion learning. Behav Neurosci 123, 764– 771 (2009).

42. Gutiérrez, R., Rodriguez-Ortiz, C. J., Cruz, V., Núñez-Jaramillo, L. & Bermudez-Rattoni, F. Cholinergic dependence of taste memory formation: evidence of two distinct processes. Neurobiol Learn Mem 80, 323–331 (2003).

43. Gordon-Fennell, A. et al. An open-source platform for head-fixed operant and consummatory behavior. eLife 12, e86183 (2023).

44. Nachman, M. Learned taste and temperature aversions due to lithium chloride sickness after temporal delays. J Comp Physiol Psychol 73, 22–30 (1970).

45. Dwyer, D. M. Microstructural analysis of ingestive behaviour reveals no contribution of palatability to the incomplete extinction of a conditioned taste aversion. Q J Exp Psychol Hove 62, 9–17 (2009).

46. Jing, M. et al. An optimized acetylcholine sensor for monitoring in vivo cholinergic activity. Nat. Methods 17, 1139–1146 (2020).

47. Miranda, M. I., Ramírez-Lugo, L. & Bermúdez-Rattoni, F. Cortical cholinergic activity is related to the novelty of the stimulus. Brain Res 882, 230–235 (2000).

48. Klapoetke, N. C. et al. Independent optical excitation of distinct neural populations. Nat. Methods 11, 338–346 (2014).

49. Marshel, J. H. et al. Cortical layer–specific critical dynamics triggering perception. Science 365, eaaw5202 (2019).

50. Xie, S. et al. Red-shifted GRAB acetylcholine sensors for multiplex imaging in vivo. Nat. Neurosci. 29, 2012–2022 (2026).

51. Goldberg, J. A., Ding, J. B. & Surmeier, D. J. Muscarinic modulation of striatal function and circuitry. Handb Exp Pharmacol 223–241, (2012).

52. Bowen, A. J. et al. Topographic representation of current and future threats in the mouse nociceptive amygdala. Nat. Commun. 14, 196 (2023).

53. Garrison, A. T. et al. Development of VU6019650: A Potent, Highly Selective, and Systemically Active Orthosteric Antagonist of the M5 Muscarinic Acetylcholine Receptor for the Treatment of Opioid Use Disorder. J. Med. Chem. 65, 6273–6286 (2022).

54. Suh, B.-C. & Hille, B. Recovery from muscarinic modulation of M current channels requires phosphatidylinositol 4,5-bisphosphate synthesis. Neuron 35, 507–520 (2002).

55. Brown, D. A. & Adams, P. R. Muscarinic suppression of a novel voltage-sensitive K+ current in a vertebrate neurone. Nature 283, 673–676 (1980).

56. Wu, W. W., Chan, C. S. & Disterhoft, J. F. Slow afterhyperpolarization governs the development of NMDA receptor-dependent afterdepolarization in CA1 pyramidal neurons during synaptic stimulation. J Neurophysiol 92, 2346–2356 (2004).

57. Chen, Y. et al. Endogenous Gαq-Coupled Neuromodulator Receptors Activate Protein Kinase A. Neuron 96, 1070–1083.e5 (2017).

58. Liu, B. et al. An Ultrasensitive PKA Biosensor for Multi-modal Kinase Activity Detection and High-Resolution Imaging in Awake Mice. FASEB J. 35, (2021).

59. Razidlo, J. A. et al. Chronic Loss of Muscarinic M5 Receptor Function Manifests Disparate Impairments in Exploratory Behavior in Male and Female Mice despite Common Dopamine Regulation. J. Neurosci. 42, 6917–6930 (2022).

60. Markram, H. & Segal, M. Acetylcholine potentiates responses to N-methyl-D-aspartate in the rat hippocampus. Neurosci Lett 113, 62–65 (1990).

61. Markram, H. & Segal, M. Long-lasting facilitation of excitatory postsynaptic potentials in the rat hippocampus by acetylcholine. J Physiol 427, 381–393 (1990).

62. Sugisaki, E., Fukushima, Y., Tsukada, M. & Aihara, T. Cholinergic modulation on spike timing-dependent plasticity in hippocampal CA1 network. Neuroscience 192, 91–101 (2011).

63. Tenk, C. M., Kavaliers, M. & Ossenkopp, K.-P. Dose response effects of lithium chloride on conditioned place aversions and locomotor activity in rats. Eur J Pharmacol 515, 117–127 (2005).

64. Hunker, A. C. et al. Conditional Single Vector CRISPR/SaCas9 Viruses for Efficient Mutagenesis in the Adult Mouse Nervous System. Cell Rep. 30, 4303–4316.e6 (2020).

65. Mahn, M. et al. Efficient optogenetic silencing of neurotransmitter release with a mosquito rhodopsin. Neuron 109, 1621–1635.e8 (2021).

66. Hjort, M. M. et al. Microprisms enable enhanced throughput and resolution for longitudinal tracking of neuronal ensembles in deep brain structures. Neurophotonics 11, 033407 (2024).

67. Fleming, W., Jewell, S., Engelhard, B., Witten, D. M. & Witten, I. B. Inferring spikes from calcium imaging in dopamine neurons. PLoS One 16, 0252345 (2021).

68. Fermani, F. et al. Food and water intake are regulated by distinct central amygdala circuits revealed using intersectional genetics. Nat. Commun. 16, 3072 (2025).

69. Higley, M. J., Soler-Llavina, G. J. & Sabatini, B. L. Cholinergic modulation of multivesicular release regulates striatal synaptic potency and integration. Nat Neurosci 12, 1121–1128 (2009).

70. Kayyal, H. et al. Activity of Insula to Basolateral Amygdala Projecting Neurons is Necessary and Sufficient for Taste Valence Representation. J. Neurosci. 39, 9369–9382 (2019).

71. Yiannakas, A. & Rosenblum, K. The insula and taste learning. Front Mol Neurosci 10, 335 (2017).

72. Arthurs, J. & Reilly, S. Role of the gustatory thalamus in taste learning. Behav Brain Res 250, 9–17 (2013).

73. Casey, E., Avale, M. E., Kravitz, A. & Rubinstein, M. Dopaminergic innervation at the central nucleus of the amygdala reveals distinct topographically segregated regions. Brain Struct Funct 228, 663–675 (2023).

74. Bernanke, A. et al. Behavior and Fos activation reveal that male and female rats differentially assess affective valence during CTA learning and expression. PLOS ONE 16, 1–34 (2021).

75. Kim, D.-I. et al. Presynaptic sensor and silencer of peptidergic transmission reveal neuropeptides as primary transmitters in pontine fear circuit. Cell 187, 5102–5117.e16 (2024).

76. Jeong, H. et al. Mesolimbic dopamine release conveys causal associations. Science 378, eabq6740 (2022).

77. Pachitariu, M. et al. Suite2p: beyond 10,000 neurons with standard two-photon microscopy. bioRxiv (2017). 10.1101/061507

78. Stringer, C., Wang, T., Michaelos, M. & Pachitariu, M. Cellpose: a generalist algorithm for cellular segmentation. Nat. Methods 18, 100–106 (2021).

79. Yu, C.-H. et al. The Cousa objective: a long-working distance air objective for multiphoton imaging in vivo. Nat. Methods 21, 132–141 (2024).

